# Cadmium-induced gut dysbiosis precedes the onset of hippocampus-dependent learning and memory deficits in mice

**DOI:** 10.1101/2025.04.07.647608

**Authors:** Hao Wang, Joe Jongpyo Lim, Haiwei Gu, Zhengui Xia, Julia Yue Cui

## Abstract

**Background:** Cadmium (Cd) is a heavy metal recognized as a neurotoxicant. However, the mechanisms underlying its neurotoxicity remain poorly understood. The gut-brain axis, a bidirectional communication pathway between the central nervous system and the gut microbiome, has been linked to various neurological disorders. Because the gut microbiome is a known target of Cd, it is important to investigate whether the gut-brain axis mechanistically contributes to the Cd-induced neurotoxicity.

**Objective:** In our initial exploration of the role of the gut-brain axis in modulating Cd neurotoxicity on cognition, we investigated whether Cd exposure induces gut dysbiosis before the onset of cognitive deficits and explored the potential link between gut microbiome alterations and Cd-induced cognitive deficits.

**Methods:** Adult male mice were exposed to 3 mg/L Cd via drinking water for nine weeks. Behavioral assessments were conducted throughout the exposure period to evaluate cognitive function. Fresh fecal pellets were collected weekly to monitor changes in the gut microbiome composition. The effects of Cd on the hippocampus and intestine were analyzed using transcriptomics and mass spectrometry (MS)-based metabolomics.

**Results:** Cd exposure resulted in hippocampus-dependent learning and memory deficits, first observed at four weeks into exposure. RNA sequencing of the hippocampus at the terminal time point revealed reduced expression of genes involved in cognition and neuroinflammation in Cd exposed mice. Metagenomic shotgun sequencing showed that Cd-induced gut dysbiosis preceded the onset of cognitive impairments, with specific bacterial species associated with Cd-induced cognitive deficits. Furthermore, Cd exposure reduced the expression of genes involved in intestinal barrier integrity, increased inflammatory cytokines levels, and altered the levels of neuroactive microbial metabolites.

**Conclusion:** Our study is the first to show that Cd exposure triggers gut microbial shifts before the onset of cognitive deficits, accompanied by increased intestinal permeability and elevated proinflammatory biomarkers in both the intestine and brain at the terminal time point. These findings suggest a potential critical role of gut-brain axis in modulating Cd neurotoxicity and underscore the need for future research to elucidate the mechanistic involvement of gut microbiome as a potential target for mitigating Cd-induced cognitive decline.

## Introduction

Cadmium (Cd) is a naturally occurring heavy metal widely used in various commercial and industrial applications ^1^. It is among the top 10 priority environmental contaminants identified by the World Health Organization (WHO) due to its significant public health risks and substantial release into the environment. The primary routes of exposure for the general population are ingestion and cigarette smoking. Because Cd has a long biological half-life in humans, it accumulates in the body and produces toxicity in multiple organs, including the liver, kidney, testis, and lung ^2,3^.

Recent studies have provided increasing evidence that Cd is a neurotoxicant ^4,5^. Epidemiological research has linked blood Cd levels to multiple neurodegenerative diseases, including Alzheimer’s disease (AD) ^6,7^ and Parkinson’s disease (PD) ^8^. Hippocampus is a major region in the brain that is responsible for learning and memory ^9^. Our previous work has demonstrated that Cd exposure via drinking water during adolescent and adulthood impairs hippocampus-dependent learning and memory in mice ^10–13^. However, the molecular and cellular mechanisms underlying Cd neurotoxicity in learning and memory remain poorly understood. Most research exploring the mechanisms of Cd neurotoxicity has focused on its direct effects on the central nervous system (CNS), including oxidative stress, neuroinflammation, as well as the disruption of blood-brain barrier (BBB) and synapse function ^14^. These studies have significantly improved our understanding of the direct impact of Cd on the CNS, however, it remains to be determined whether Cd-mediated toxicity in other bio-compartments, such as the gut microbiome, may indirectly contribute to neurotoxicity through remote-sensing mechanisms ^15^.

Mounting evidence suggests that the gut microbiome plays an important role in regulating cognitive functions. Human studies reported that AD patients exhibit reduced microbial diversity, with their gut microbiome significantly differing from individuals with normal cognition ^16,17^. Similarly, dysbiosis of the gut microbiota has been observed in a transgenic mouse model of AD ^18,19^. Limited human case reports also suggest that Fecal microbiome transplantation (FMT) may enhance memory and cognitive performance in AD patients, but robust clinical trials are needed to validate these findings ^20,21^. Furthermore, multiple studies have found that probiotics treatment can improve learning and memory both in mice ^22^ and humans ^23–25^. Mechanistically, the gut microbiome is suggested to regulate the CNS functions through the gut-brain axis, a bidirectional signaling mechanism between the gastrointestinal (GI) tract and the CNS ^15^. Neuroactive microbial metabolites, such as short-chain fatty acids (SCFAs) and secondary bile acids (BAs), play a pivotal role in these “remote-sensing” mechanisms ^26^. For example, butyrate, a neuroprotective SCFA produced through bacterial fermentation in the GI tract, can improve learning and memory in mice ^15,27–29^. Moreover, certain microbial-derived secondary BAs have been implicated in the pathogenesis of AD ^28,29^.

In addition to the production of neuroactive microbial metabolites, the gut microbiome can influence the CNS functions by modulating intestinal barrier integrity and inflammation. Alteration in gut microbiome-derived metabolites (such as SCFAs) and/or pathogenic bacterial products (such as lipopolysaccharides) can compromise the integrity of the intestinal barrier ^26,30^. Disruption of the gut barrier can further facilitate the direct translocation of gut microbiome components and their products into the bloodstream, allowing them to reach the brain, where they can impair the integrity of the BBB and impair the CNS functions ^26^. Furthermore, changes in the microbiome and its metabolites can trigger the inflammatory responses in intestinal and circulating immune cells, further contributing to the induction and exacerbation of central neurogenic and inflammatory responses in the CNS, ultimately leading to the CNS dysfunction ^31,32^. In addition, increasing studies have demonstrated that morphological and functional changes in the intestinal barrier, as well as systemic inflammatory response are associated with various neurological disorders, including AD, PD, multiple sclerosis (MS), autism spectrum disorder, mild cognitive impairments, and depression ^32–37^.

Previous studies have examined the effects of Cd exposure on the gut environment ^38^. Epidemiologic research has found that Cd exposure is associated with altered fecal microbiome diversity and composition in humans ^39^. Animal studies also have shown that Cd exposure can impair intestinal integrity and alter the gut microbiome composition in mice ^40–42^. In addition, Cd exposure reduces SCFA-producing bacteria in the gut, leading to decrease in the SCFAs production ^41,43^. Together, these findings suggest that the gut microbiome may contribute to Cd-induced neurotoxicity in learning and memory. However, most previous animal studies have examined the toxic effects of Cd on the gut microbiome at a single terminal time point. To date, no studies have reported the effects of Cd exposure on the gut microbiome over a time course and explored the potential link between the gut microbiome changes and the Cd-induced neurotoxicity.

Despite growing evidence linking environmental toxicants to gut microbiome disruptions and neurological dysfunction, the temporal relationship between gut dysbiosis and Cd-induced cognitive deficits remains poorly understood. While Cd exposure has been shown to impair cognitive functions and alter gut microbial composition separately, it remains unclear whether gut dysbiosis precedes cognitive deficits or arises as a secondary consequence of Cd neurotoxicity. Furthermore, the specific microbial taxa and metabolic pathways involved in Cd neurotoxicity have not been well characterized, limiting our understanding of potential microbiome-mediated mechanisms. To address this critical knowledge gap, we hypothesized that gut microbiome dysbiosis precedes the onset of Cd-induced cognitive deficits and contributes to Cd neurotoxicity. As an initial step in exploring the role of the gut-brain axis in Cd neurotoxicity, the present study aimed to characterize the compositional and metabolic changes of the gut microbiome over a time course following Cd exposure and to identify distinct Cd-regulated gut microbial species associated with Cd-induced hippocampus-dependent learning and memory deficits in mice.

## Materials and Methods

### Chemicals

Cadmium Chloride (CdCl_2_) was purchased from MilliporeSigma (cat # 202980, Burlington, MA). Unless otherwise noted, all other chemicals were also purchased from MilliporeSigma (Burlington, MA).

### Animals and exposures

Six-week-old male C57BL/6 mice were purchased from Charles River Laboratories and housed (5 animals per cage) under standard conditions (12-hour light/dark cycle) with ad libitum access to feed (Picolab Rodent Diet 20, LabDiet, St. Louis, MO) and water (tap water purified by reverse osmosis, acidified with 2.4-2.8% HCl, and autoclaved) provided *ad libitum.* Starting at 8 weeks of age, mice received normal drinking water or drinking water with 3 mg/L Cd (in the form of CdCl_2._) for 9 weeks (Fig. 1). Behavioral tests were conducted before and during the whole exposure period to investigate the effects of Cd exposure on the mice. Feces were collected before exposure (week 0) and at weeks 1, 3, 5, 7, and 9 during the exposure period. Tissue collection was performed at the end of the study. The preparation, use, and disposal of hazardous reagents were conducted according to the guidelines set forth by the Environmental Health and Safety Office at the University of Washington. All animal care and exposure were approved by the University of Washington Institutional Animal Care and Use Committee (IACUC).

**Figure 1.**
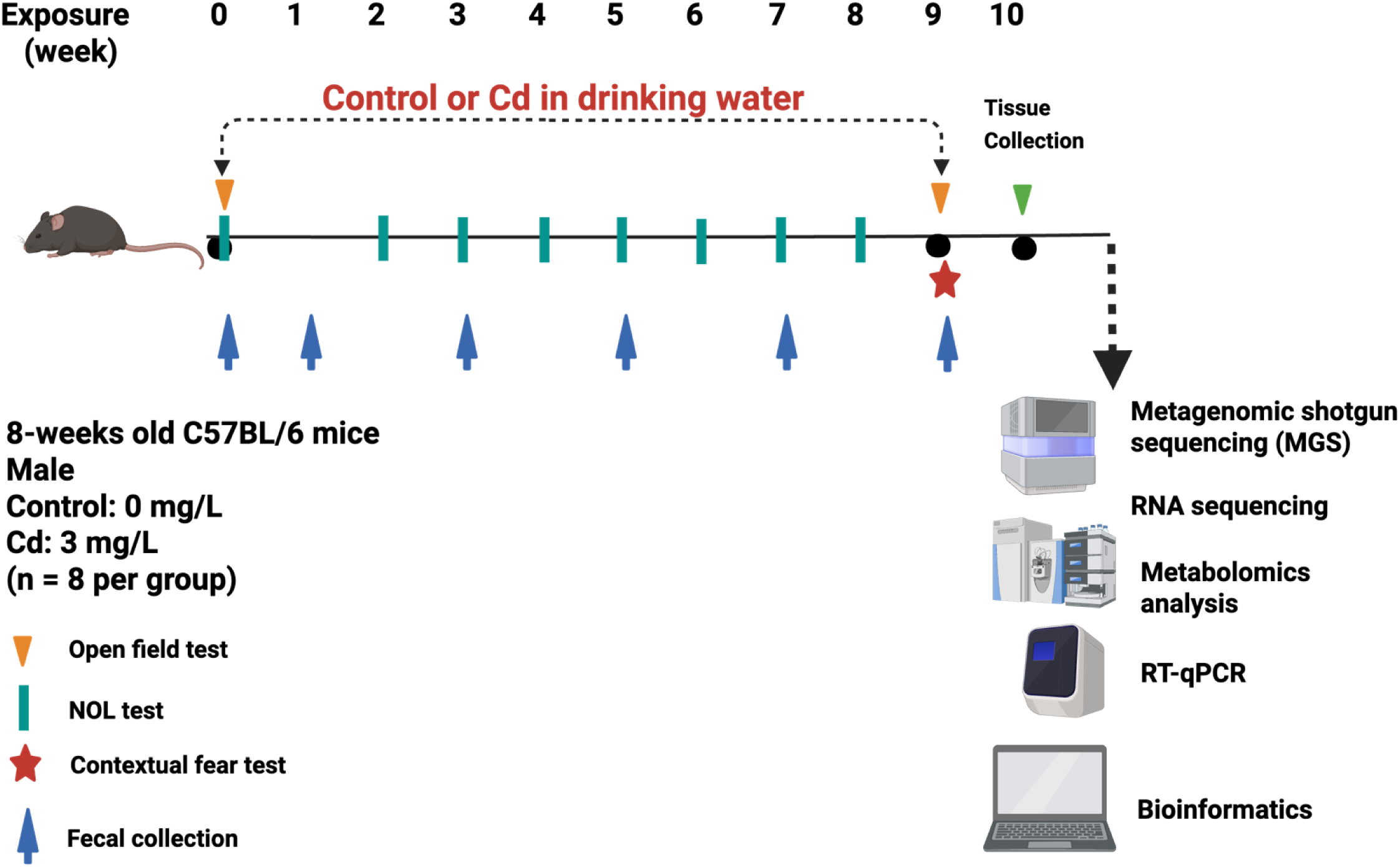
Experiment design and timeline for different tests and data analysis.

### Open field test

The open field test was used to assess the effects of Cd on locomotor activity and anxiety ^44^. Mice were placed into a 10” (width) × 10” (depth) × 16” (height) TruScan Photo Beam Tracking arena (Coulbourn Instruments, Whitehall, PA) with clear Plexiglas sidewalls, and their movement was monitored with two sets of infrared breams spaced 0.6” apart during the test. Mice were allowed to freely explore the arena without pre-habituation for 20 min, and data were collected by TruScan 2.0 software (Coulbourn Instruments). The total move distance, total move time, and average speed were used to assess locomotor activity. In addition, the time spent in the margin and center, the margin and center distance, and center entries were used to examine anxiety.

### Novel object location test

The Novel object location test (NOL) was used to assess the effects of Cd on hippocampus-dependent spatial memory ^13^. This assay was performed as previously described. Briefly, mice were placed into an open field arena (Coulbourn Instruments) with two identical objects placed in 2 different corners. During the training session, mice were allowed to freely explore the two objects for 5 min and then returned to their home cages (Fig. 2A). To exclude preference for a specific location, alternating corners were used for object presentation. The testing session was performed 1 h after the training session, and animals were returned to the arena with the same two objects; one object remained in its original location, and the other had been moved to a new location. The time animals spent actively investigating each object during the training and testing sessions was recorded by cameras and quantified by an experimenter blinded to the animal’s treatment after the test. Although our previous studies have demonstrated that memory established during the training and testing session typically lasts less than 24 hours, and the results of the NOL tests remain consistent when conducted weekly^11,13,44^, we switched to different objects during the consecutive two weeks of testing to minimize the potential effects of repeated NOL tests on mice performance.

**Figure 2.**
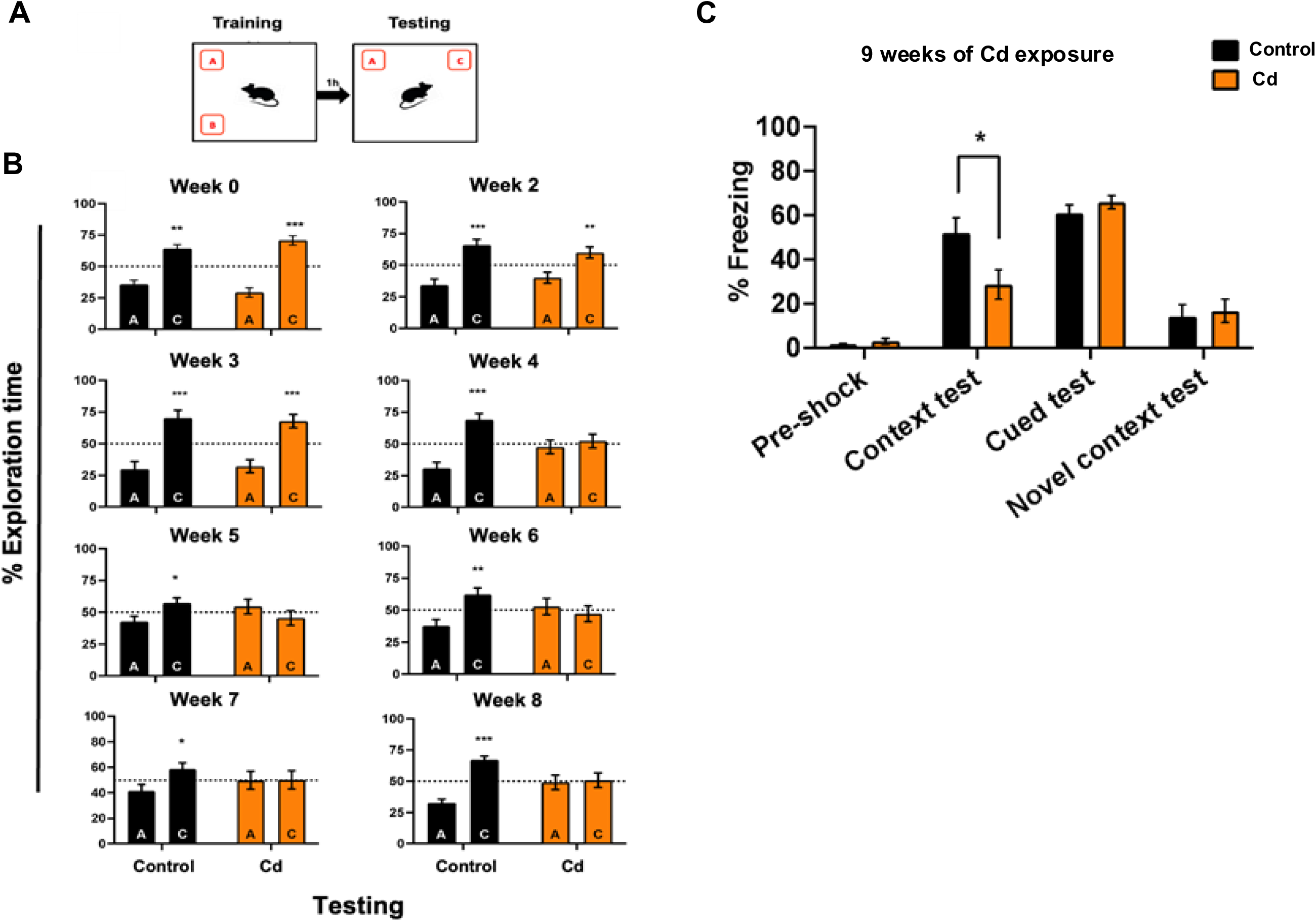
Cd exposure impairs the hippocampus-dependent learning and memory in mice. **A.** Schematic of 1hour Novel Object Location (NOL) test. **B.** The results of testing sessions at different time points. Cd-exposed mice exhibited spatial memory deficits starting at 4 weeks into Cd exposure. **C.** Cd exposure impairs the contextual fear memory in mice. However, auditory-cued fear memory and general freezing behavior was not affected in Cd-exposed mice. Data are presented as mean ± SEM with n = 8 in each group.

### Cued and contextual fear-conditioning tests

In this study, a modified cued and contextual fear conditioning test using the weak foot shock conditioning paradigm (3 × 0.3 mA, 2 s shocks with 2 min inter-trial intervals) was used as previously described ^11,13,44^. For the conditioning session, mice were placed into the foot shock context (10” × 10”× 16” arena with grid shock floor; Coulbourn Instruments) and allowed to explore the arena for 2 min until the presentation of a 90 dB, 30 s tone (conditioned stimulus, [CS]). During the last 2 s of the tone, a 0.3 mA foot shock (unconditioned stimulus, [US]) was delivered. This cycle was repeated two more times before the mice were returned to their home cages. The CS and US were automatically delivered by the TruScan software (Coulbourn Instruments). The contextual fear memory test was performed 24 h after the conditioning session. Mice were placed back into the same foot shock context for 2 min in the absence of tone or shock. For the cued test, which was performed 2 h after the context test, mice were placed into a novel context (new room; hexagonal Plexiglas arena) and allowed to freely explore for 2 min. The CS (tone) was then presented to them for 2 min. For the novel context test, which was performed 2 h after the cued test, mice were placed into another novel context (new room; rat cage) and allowed to freely explore for 2 min with no presentation of either tone or foot shock. In all three tests, persistent freezing behavior (four paws on the ground, no head or body movement besides breathing) was recorded by video cameras and manually quantified by an experimenter blinded to animal exposure.

### Fecal DNA extraction and shallow metagenomic shotgun sequencing

DNA was extracted from 100 - 200 mg fecal samples by using an AllPrep PowerFecal DNA/RNA Kit (Qiagen, Hilden, Germany) according to the manufacturer’s protocol. DNA concentration was quantified using a Qubit 2.0 Fluorometer (Life Technologies, Grand Island, NY). The integrity and quality of DNA samples were confirmed by using an Agilent 2100 Bioanalyzer (Agilent Technologies Inc., Santa Clara, CA). Shallow metagenomic shotgun sequencing was performed at 2 million reads by Diversigen (New Brighton, MN). DNA sequences were aligned to a curated database containing all representative genomes in RefSeq for bacteria with additional metagenomically assembled genomes (MAGs) and cell-cultured genomes. Only high-quality MAGs (Completeness > 90% and Contamination < 5% via checkm) were included for downstream analysis. Alignments were made against all reference genomes at 97% identity against all reference genomes. Every input sequence was compared to every reference sequence in the Diversigen Venti database using fully gapped alignment with BURST. Ties were broken by minimizing the overall number of unique Operational Taxonomic Units (OTUs). For taxonomy assignment, each input sequence was assigned the lowest common ancestor that was consistent across at least 80% of all reference sequences tied for best hit. The taxonomy data were based on the Genome Taxonomy Database (GTDB r95). Samples with fewer than 10,000 sequences were removed from further analysis. OTUs accounting for less than one-millionth of all strain-level markers and those with less than 0.01% of their unique genome regions covered (and < 0.1% of the whole genome) at the species level were also discarded. Differential abundance testing for the taxonomic count data was analyzed using a mixed effects model adjusting for baseline at week 0 using MaAsLin2 ^45^ with default parameters. Alpha (Chao1 index and Shannon index) and beta (Bray-Curtis) diversity were calculated using the taxonomy table using QIIME 1.9.1^46^. Taxa were normalized using the centered-log ratio (CLR) method ^47^ and plotted using the R packages ComplexHeatmap (Version 2.16.0) ^48,49^ and ggplot2 (version 3.4.4) (Wickham, 2016). Taxa that were differentially abundant between control and Cd exposure group in at least 2 time points were correlated with the novel object location test results using the cor.test function (Spearman correlation, p < 0.1) in R. Raw sequencing files can be found in Sequence Read Archive (SRA) with accession PRJNA1245492.

### RNA-Seq of hippocampus transcriptome

Total RNA was isolated from the hippocampus of control or Cd exposure mice after exposure using the RNA-Bee reagent (Tel-Test Inc, Friendswood, TX) according to the manufacturer’s protocol. The concentrations of RNA were quantified by using a NanoDrop 1000 Spectrophotometer (Thermo Scientific, Waltham, MA) at 260 nm. The quality of RNA was evaluated by formaldehyde-agarose gel electrophoresis by visualizing the 28S and 18S rRNA bands under UV light and Agilent 2100 Bioanalyzer by Novogene. The RNA sequencing was performed by Novogene (Sacramento, CA) using 5 biological replicates.

FASTQ files with paired-end sequence reads were mapped to the mouse genome (UCSC mm10) using HISAT2 (Hierarchical Indexing for Spliced Alignment of Transcripts) ^50^. The resulting SAM (sequence alignment/map) files were converted to their binary form and sorted using SAMtools (version 1.2) ^51^. Transcript abundances were estimated with featureCounts (part of the Subread package, version 1.5.3) ^52^ using the Gencode mouse version 25 (vM25) gene transfer format (GTF). Differential expression was performed using DESeq2 ^53^ with false discovery rate (FDR)-adjusted p-value < 0.05. Gene ontology enrichment was conducted using the R package TopGO package (version 2.52.0) ^54^. Count data were normalized to transcript per million (TPM). Principal components analysis (PCA) was performed on the filtered TPM normalized counts (above 1^st^ quantile of standard deviation) using the princomp function in R. Differentially expressed genes involved in learning, memory, or cognition, as well as inflammation were retrieved from the Gene Ontology database ^55^ and were correlated with differentially abundant taxa at week 9 using cor.test function (Spearman correlation, FDR-adjusted p-value < 0.1) in R. Data were plotted using ComplexHeatmap (version 2.16.0) ^48,49^ and ggplot2 (version 3.4.4) ^56^. Raw sequencing files and processed counts data can be found in Gene Expression Omnibus (GEO) with accession GSE293538.

### RT-qPCR quantification of intestinal tissue

Total RNA isolated from intestinal tissues (duodenum, jejunum, ileum, and colon) was reverse transcribed into cDNA using a high-capacity cDNA Reverse Transcription Kit (Life Technologies, CA). The resulting cDNA products were then amplified by qPCR, using the Sso Advanced Universal SYBR Green Supermix in a Bio-Rad CFX384 Real-Time PCR detection system (Bio-Rad, Hercules, CA). Primer sequencing is shown in the Supplementary Table. Data were normalized to the housekeeping gene using the ΔΔCq method and were expressed as % of the housekeeping gene.

### Serum cytokine levels quantification

Serum cytokine levels were quantified using the MILLIPLEX^®^ Mouse High Sensitivity T Cell Magnetic Bead Panel assay (Millipore Sigma, cat. # MHSTCMAG-70KPMX) following the manufacturer’s instructions. Briefly, quality controls, standards, and serum samples were prepared in triplicate and added into a 96-well plate pre-coated with magnetic beads conjugated with capture antibodies specific to target cytokines. After an overnight incubation at 4 °C, plates were washed by using a wash buffer to remove unbound components. Detection antibodies were added into each well and incubated for one hour at room temperature, followed by the addition of streptavidin-phycoerythrin (PE) and a half-hour. After 3 times of gently wash, beads were resuspended in assay buffer and analyzed using the Belysa^®^ Immunoassay Curve Fitting Software (MilliporeSigma).

### Short-chain fatty acids quantification

Briefly, 50 mg of each tissue sample was homogenized with 20 μL hexanoic acid-6,6,6-d_3_ (internal standard; 200 µM in H_2_O), 20 μL sodium hydroxide solution (NaOH, 0.5 M in water), and 480 μL methanol (MeOH). Afterwards, 400 μL MeOH was added to adjust the pH to 10. Upon storage at -20°C for 20 min and centrifugation at 21,694 g for 10 min, 800 μL of supernatant were collected. Samples were then evaporated to dryness, reconstituted in 40 μL of methoxyamine hydrochloride in pyridine (20 mg/mL), and stored at 60°C for 90 min. Afterward, 60 μL of N-Methyl-N-tert-butyldimethylsilyltrifluoroacetamide was added and stored at 60°C for 30 min. Each sample was then vortexed for 30 s and centrifuged at 21,694 g for 10 min. Finally, 70 μL of supernatant was collected from each sample for GC-MS analysis. For serum/intestinal content samples, 20 μL of each sample was mixed with 30 μL aqueous NaOH (0.1M in water), 20 μL IS (hexanoic acid- 6,6,6-d_3_; 200 µM) and 430 μL MeOH in a 1.5 mL Eppendorf tube. The pH value for the mixture was 9. Samples were then vortexed for 10 s and stored under -20 °C for 20 min. After centrifugation at 14,000 RPM for 10 min at 4 °C, 450 μL supernatant was removed into a new Eppendorf tube. The samples were dried under vacuum at 37 °C for 120 min using a CentriVap Concentrator (Labconco, Fort Scott, KS). Each sample was first derivatized with 40 µL of methoxyamine hydrochloride solution in pyridine (MeOX, 20 mg/mL) under 60 ℃ for 90 min. Next, 60 µL of MTBSTFA was added, and the mixture was incubated under 60 ℃ for 30 min. Then the sample was vortexed for 30 s, followed by centrifugation at 14,000 rpm for 10 min. Finally, 70 µL supernatant was collected into a new glass vial for GC-MS analysis. GC-MS experiments were performed using an Agilent 7820A gas chromatography system coupled to an Agilent 5977B mass spectrometer (Agilent Technologies, Santa Clara, CA). Chemical derivatives in the samples were separated using an HP-5 ms capillary column coated with 5% phenyl-95% methylpolysiloxane (30 m×250 µm i.d., 0.25 µm film thickness, Agilent Technologies). 1 µL of each sample was injected, and the solvent delay time was set to 5 min. The initial oven temperature was held at 60 ℃ for 1 min, ramped up to 325 ℃ at a rate of 10 ℃/min, and finally held at 325 ℃ for 10 min. Helium was used as the carrier gas at a constant flow rate of 20 mL/min through the column. The temperatures of the front inlet, transfer line, and electron impact (EI) ion source were set at 250 ℃, 290 ℃, and 230 ℃, respectively. The electron energy was -70 eV, and the mass spectral data were collected in the full scan mode (*m/z* 30-600). Agilent MassHunter Workstation Software Quantitative Analysis (B.09.00) was used to process the GC-MS data for compound identification, peak picking, and quantification. The signal-to-noise ratio (S/N) was set to S/N=3. The retention time and quantification mass for each SCFA were determined using its chemical standard. The concentrations of SCFAs in biological samples were calculated using the calibration curves constructed from the corresponding SCFAs standards.

### Bile acid quantification

Bile acids were quantified in various compartments using a method that we described previously ^57,58^. Generally, 1 mg/mL stock solutions of individual BAs (for standard curve) and internal standards (ISs) were prepared in methanol and water (1:1). The 19 individual BA stock solutions were further diluted in 50% methanol to obtain 10 working standard solutions (0.05-10000 ng/mL). The 5 ISs were mixed to obtain a working IS solution. For serum BA extraction, 50 μL of serum samples were mixed with 10 ul of internal standard (IS) solution and vortexed for 5-10 minutes. 500 μL of ice-cold methanol was added to the serum samples and vortexed again to prepare a homogenous mixture. The sample mixture was centrifuged at 21000 g for 10 minutes at 4 °C. The supernatant was collected in a new tube and again 500 μL of ice-cold methanol was added to the pellet. The pellet was dissolved in methanol and again centrifuged following the same previous criteria. Each supernatant was combined and evaporated under vacuum (low temperature) for 2.5 hours. The dried samples were then reconstituted using 100ul 50% methanol. Before injecting the samples were again centrifuged for 10 minutes at 21,000 g and 50 ul of filtered samples were loaded in the autosampler vials for MS-analysis. Agilent 1290 UPLC (ultra-high pressure liquid chromatography) system combined with an Agilent 6460 triple quadrupole mass spectrometer via an electrospray ionization interface, was used for analysis. The chromatographic separation was performed using a ZORBAX Eclipse Plus C18 analytical column (2.1X100mm; id: 1.8µm). Samples were eluted using mobile phase A, consisting of 20% acetonitrile and 10 mM ammonium acetate in water, and mobile phase B, which consisted of 80% acetonitrile and 10 mM ammonium acetate in water, at a flow rate of 0.4 mL/min. The injection volume of the samples was 5 µL. The column temperature was set at 45 °C, and the sample tray temperature was maintained at 9 °C. MS/MS spectra were produced using the negative ionization mode.

### Statistical analysis

Statistical analyses were performed using GraphPad Prism software (GraphPad Software Inc, La Jolla, CA). The Student’s two-tailed t-test (α = 0.05) was used to analyze all results of behavioral tests, short-chain fatty acids analysis, Bile acid analysis, and RT-qPCR. Taxa that were differentially abundant between control and Cd exposure group in at least 2 time points were correlated with the novel object location test results using the cor.test function (Spearman correlation, Benjamini & Hochberg adjusted p < 0.2) in R. All data were expressed as mean SEM. n.s. not significant; *p < 0.05; **p < 0.01; ***p < 0.001.

## Results

### 1. Cd exposure impaired hippocampus-dependent learning and memory in mice

To investigate the effects of Cd exposure on hippocampus-dependent learning and memory, we exposed 8-week-old male C57BL/6 mice to 3 mg/L Cd through drinking water for 9 weeks (Fig.1A). The Cd exposure level was chosen based on our previous studies ^10,13^. We monitored body weight over nine weeks and observed no differences between the control and Cd-exposure groups (Fig. S1), aligning previous findings ^13^. In addition, our previous study demonstrated that the same exposure level of Cd had no effects on locomotor activity or anxiety in mice^13^. In the present study, our open-field test results (Fig. S2) further supported these findings.

To assess the onset of hippocampus-dependent learning and memory deficits, we conducted the 1-hour NOL test before initiating Cd exposure and then weekly from the second week of the exposure onward. Throughout all NOL tests, both groups spent similar amount of time exploring each object/location during the training session, indicating no inherent preference for either object or location (Fig. S3A). In the testing session (Fig. 2B), before Cd exposure, as expected, both groups spent more time exploring the object in the novel location (C) than the object in the original location (A), demonstrating their ability to remember the original object location from training. During the first three weeks of Cd exposure, both groups retained intact spatial memory. However, after four weeks of Cd exposure, while the control group continued to distinguish between the novel and original locations, the Cd-exposed group started to exhibit memory deficits, as indicated by equal amount of time spent exploring objects in both locations. This impairment in the Cd-exposed mice persisted for the remainder of the exposure period.

We conducted the contextual fear conditioning test to examine the effects of Cd exposure on contextual fear memory, another form of hippocampus-dependent learning and memory. During the training phase, both groups exhibited a similar increase in freezing behavior following each foot shock, indicating the success of contextual fear training (Fig. S3B). The testing session was conducted 24 hours after the training, where Cd-exposed mice displayed reduced freezing behavior compared to the control group, suggesting impaired contextual fear memory. However, no differences in freezing behavior were observed between the control and Cd-exposed groups during the cued test or novel context test (Fig. 2C). Together, these findings indicate that Cd exposure impaired hippocampus-dependent contextual fear memory in mice.

### 2. Cd exposure induced altered the expression of genes associated with learning, memory, and cognition in the hippocampus

RNA sequencing was conducted to characterize transcriptional alterations in the hippocampus of mice following nine weeks of Cd exposure. Principal component analysis revealed distinct clustering based on Cd exposure status (Fig. S4A), indicating that exposure is the primary factor influencing the transcriptomic signatures.

A volcano plot (Fig. S4B) and a hierarchical clustering dendrogram revealed 1133 differentially expressed genes, including 513 genes upregulated (Cluster 1) and 620 down-regulated genes (Cluster 2) by Cd exposure (Fig. S4C). To link these genes to biological functions, we conducted Gene Ontology (GO) enrichment analysis. Notably, Cd exposure down-regulated genes associated with learning and memory, cognition, muscle system process, and locomotory behavior (Fig. 3A).

**Figure 3.**
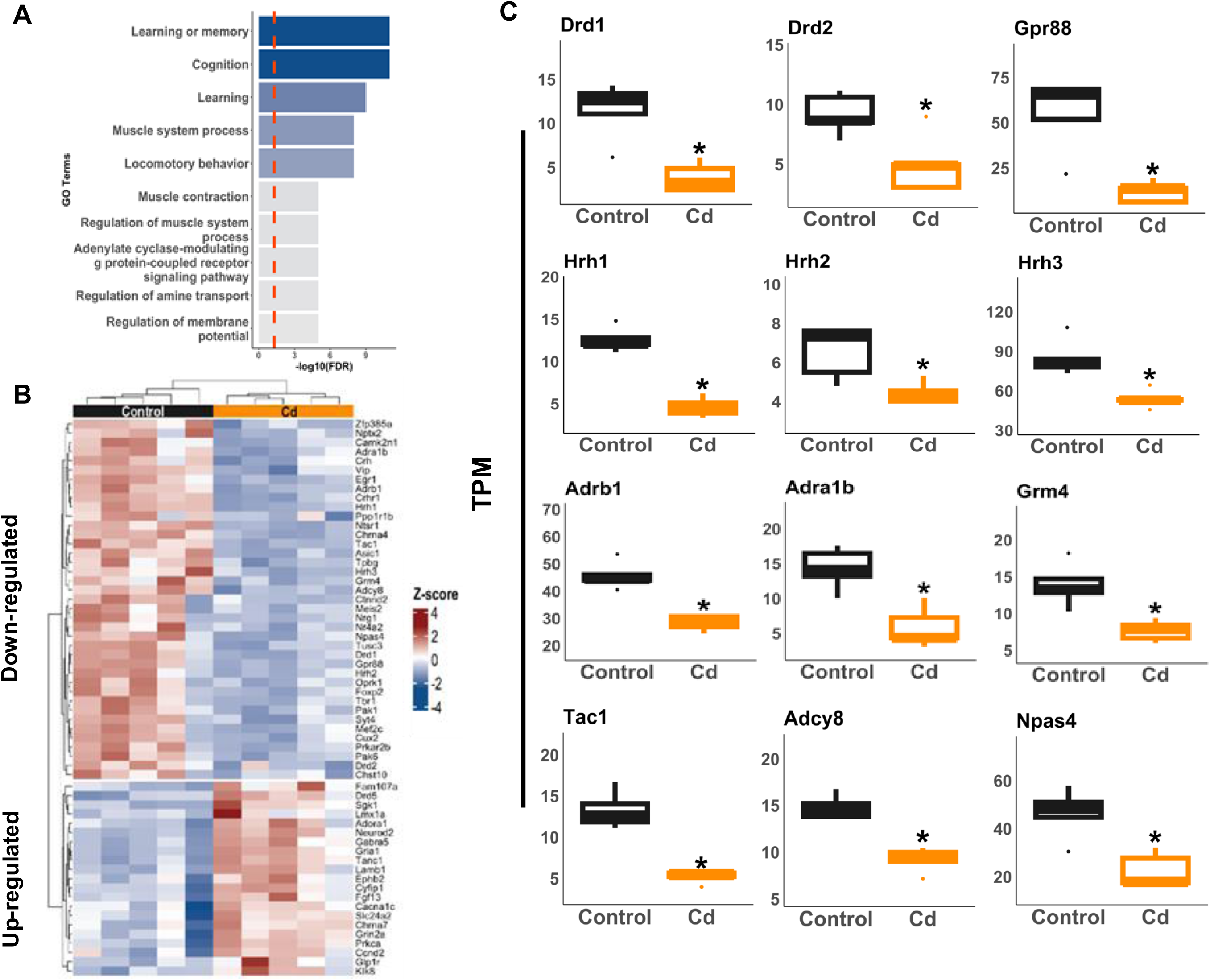
Dysregulated expression signatures of learning, memory, and cognition-related genes in the hippocampus after Cd exposure. **A.** Top 10 Gene Ontology enrichment of downregulated genes in the hippocampus following Cd exposure. **B.** Learning, memory, and cognition-related genes that are differentially expressed after 9 weeks of Cd exposure in the hippocampus are shown in a heatmap. The colors of the heatmap represent the log_2_-fold change of hippocampus genes of the Cd-exposed mice as compared with the control group. Red indicates upregulated, and Blue indicates downregulated from Cd-exposed mice (Bonferroni-adjusted p-value < 0.05). **C.** Examples of differentially regulated genes associated with learning, memory, and cognition. Asterisks represent statistically significant differences between Cd and Control groups. n = 5 in each group. *. p < 0.05.

We further examined the expression of genes involved in learning and memory, as well as cognition in the hippocampus of control and Cd-exposed mice (Fig. 3B). More cognition-related genes (39) were downregulated than upregulated (21) following Cd exposure. Specifically, Cd exposure dysregulated the expression of G protein-coupled receptors (GPCRs) in the hippocampus. For example, Cd downregulated dopamine receptors D1 *(Drd1)*/D2 (*Drd2*), histamine receptor H1 (*Hrh1*)/H2 (*Hrh2*)/H3 (*Hrh3*), adrenergic receptor alpha-1b (*Adra1b*)/beta-1 (*Adrb1*), metabotropic glutamate receptor 4 (*Grm4*), and probable G-protein coupled receptor 88 (*Gpr88*) (Fig. 3C). To note, GPCRs, the largest family of membrane proteins, play a critical role in enabling the central nervous system to respond accurately to external and internal stimuli, thereby modulating learning and memory ^59^. Additionally, Cd reduced the mRNA of tachykinin 1 (*Tac1*), which encodes tachykinin peptide hormone family that function as neurotransmitters ^60^, and adenylate cyclase type 8 (*Adcy8*), which catalyzes the cyclic adenosine monophosphate (cAMP) formation to active cAMP signaling, influencing synaptic plasticity ^61^. Additionally, Cd treatment downregulated the Neuronal PAS Domain Protein 4 (Npas4), an important regulator of synaptic plasticity ^62,63^ (Fig. 3C). These findings on the hippocampal transcriptome align with behavioral test results, which further demonstrated deficits in hippocampus-dependent learning and memory in Cd-exposed mice.

### 3. Cd exposure produced gut dysbiosis before the onset of learning and memory deficits in mice

To investigate whether Cd exposure alters the gut microbiome and the timing of the alternation relative to the onset of cognitive deficits, we performed MGS on fecal pellets collected at baseline as well as at weeks 1, 3, 5, 7, and 9 of Cd exposure. As shown in (Fig. S5 – S9), the overall gut microbiome composition remained relatively stable despite Cd exposure, as indicated by the lack of significant differences in alpha and beta diversity between the control and Cd-exposed groups. However, Cd exposure selectively altered the relative abundance of specific bacterial taxa, suggesting targeted disruption within the microbial community rather than broad shifts in overall diversity. Interestingly, Cd differentially regulated distinct microbes at early time points (weeks 1 and 3), before the observed onset of learning and memory deficits.

### 4. Persistently dysregulated microbes are highly correlated with changes in cognitive functions

To further investigate the association between the gut microbiome alterations and Cd-induced cognitive deficits, we identified microbes with their relative abundance altered in at least two time points between the control and Cd-exposed groups (Fig. S10 – S12). We then examined the relationship between these microbes and animals’ performance on the NOL test. As shown in Fig. 4A, 21 persistently dysregulated microbial taxa were significantly associated with the performance of mice in the NOL test (adjusted p-value < 0.2). Among these, eight bacteria exhibited particularly strong correlations than the others (adjusted p-value < 0.1), including UBA660, *Flavonifractor*, *Erysipelotrichaceae*, *Flintia butyricum*, *Faecalibacterium prausnitzii*, *Proteobacteria*, *Enterobacteriaceae*, and *Escherichia* (Fig. 4A). Among these 8 taxa, Cd exposure persistently decreased the abundance *UBA660* and microbe within the *Erysipelotrichaceae,* which are predominantly found in the gut microbiota of humans ^64^. Conversely, the abundance of *Flintia butyricum*,*Faecalibacterium prausnitzii*, *Flavonifractor* and *Escherichia*, *Enterobacteriaceae*, and *Proteobacteria* were persistently increased in Cd-exposed mice (Fig. 4B). Furthermore, the relative abundance of several bacteria previously found to be associated with learning and memory were persistently altered by Cd exposure, including *Akkermansia*, *Angelakisella*, and *Clostridiales* ^65–71^. As shown in Fig. 4B, *Akkermansia* were reduced in the Cd-exposed group, whereas *Angelakisella* and *Clostridiales* were increased in Cd-exposed mice. However, their correlations (adjusted p-value < 0.2 ^72–76^) with the performance of animals in the behavioral test are not as strong as those of the previous eight taxa (adjusted p-value < 0.1).

**Figure 4.**
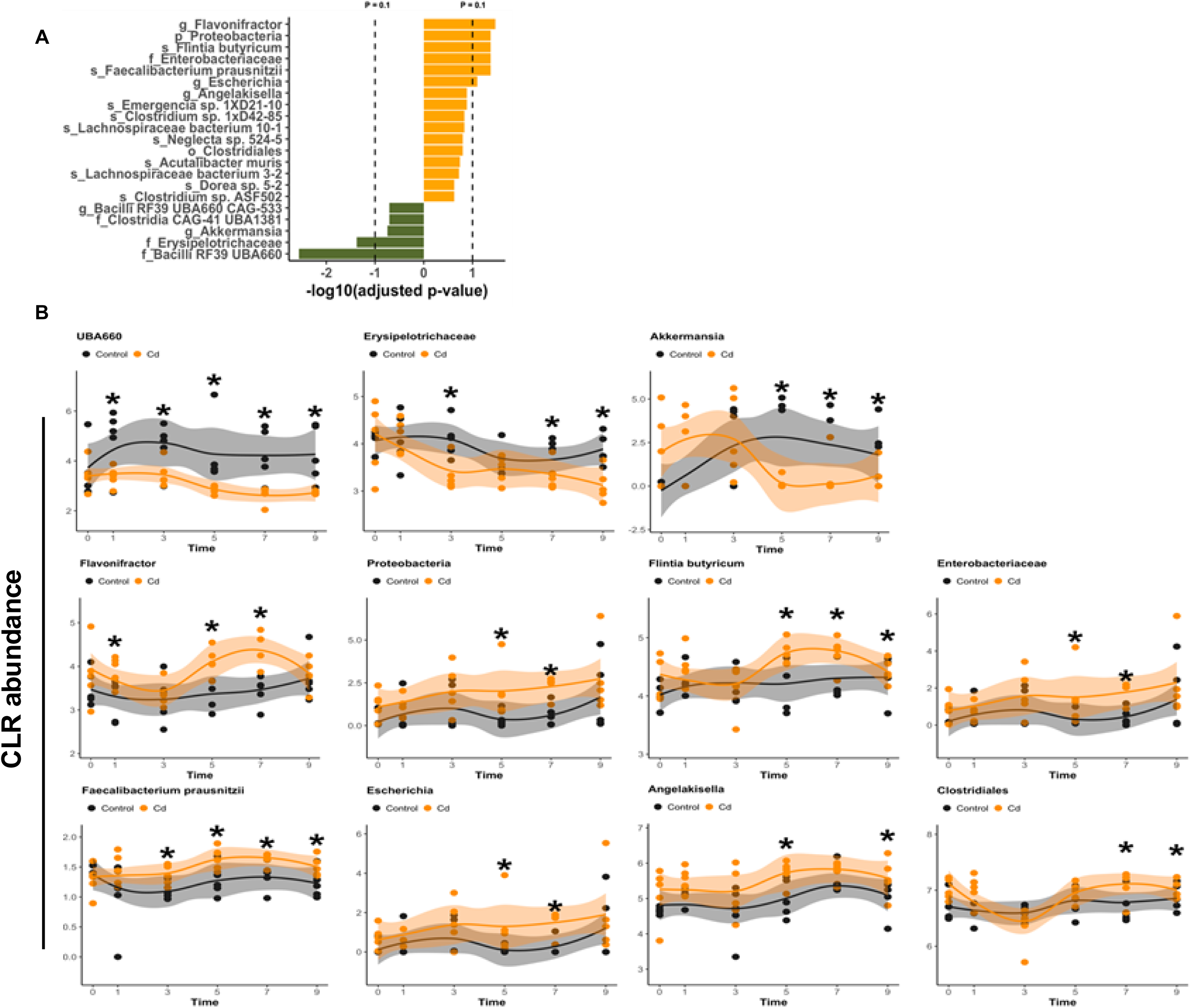
Persistent dysregulated gut microbiomes are correlated with Cd-induced learning and memory deficits. **A.** Significant correlation between the persistently dysregulated microbiome (≥ two different time points) and the performance of mice in the NOL test (adjusted p-value < 0.2). **B.** Examples of the persistently dysregulated microbiome that are significantly correlated with the performance of mice in the NOL test (adjusted p-value < 0.1). n = 5 in each group.

### 5. Cd exposure reduced intestinal SCFAs, tight junction transcripts, and SCFA receptors in mice, while increasing pro-inflammatory cytokines in serum

SCFAs are well-known to have anti-inflammatory properties and protect the intestinal barrier integrity ^77^ ^78^. Disruption of the intestinal barrier, coupled with gut microbiome dysbiosis, may increase the systemic and CNS bioavailability of harmful microbial products and pro-inflammatory cytokines, which may subsequently contribute to neurotoxicity ^79,80^,. Given these connections, we examined how Cd exposure influences endogenous microbial metabolites in the blood, intestine content, and brain, and quantified the mRNA expression of tight junction proteins, as well as SCFAs transporters and receptors in the intestine.

In the intestine, Cd exposure decreased acetic and butyric acids levels in both small and large intestinal contents(Fig. 5A). In addition, Cd exposure decreased the mRNA expression of SCFA receptors – free fatty acid receptor 2/3 (Ffar2/3) in colon (Fig. 5B). Cd exposure also downregulated the mRNAs of multiple tight junction proteins across three different sections of the intestine (Fig. 6). In the duodenum, the expression level of *Cldn7* was significantly decreased in the Cd-treated group. In the ileum, Cd exposure reduced the levels of *Tjp1 and Tjp2, Jam1 and Jam2,* and *Cldn7*, while in the colon, it decreased the expression levels of *Cldn1, Cldn2, and Cldn7 and Jam1.* Cd also decreased the mRNA of the xenobiotic efflux transporter ATP-binding cassette sub-family G member 2 (*Abcg2*), also known as the breast cancer resistance protein (*Bcrp*), which is an ATP-dependent efflux pump located on the apical side of intestine epithelial cells ^81^. In addition, BCRP is known for the efflux of the SCFA butyrate in intestinal cells ^82,83^, and its downregulation by Cd aligns with the Cd-mediated decrease in butyrate in the intestinal content (Fig. 5A) as well as increased serum SCFAs including butyric acid (Fig. 7A). However, other conventional SCFA transporters remained unaffected by Cd exposure (Fig. S13). Together, these findings suggest that Cd exposure reduces SCFA levels and downregulates the expression of various tight junction proteins, as well as SCFAs transporters and receptors, in the intestine.

**Figure 5.**
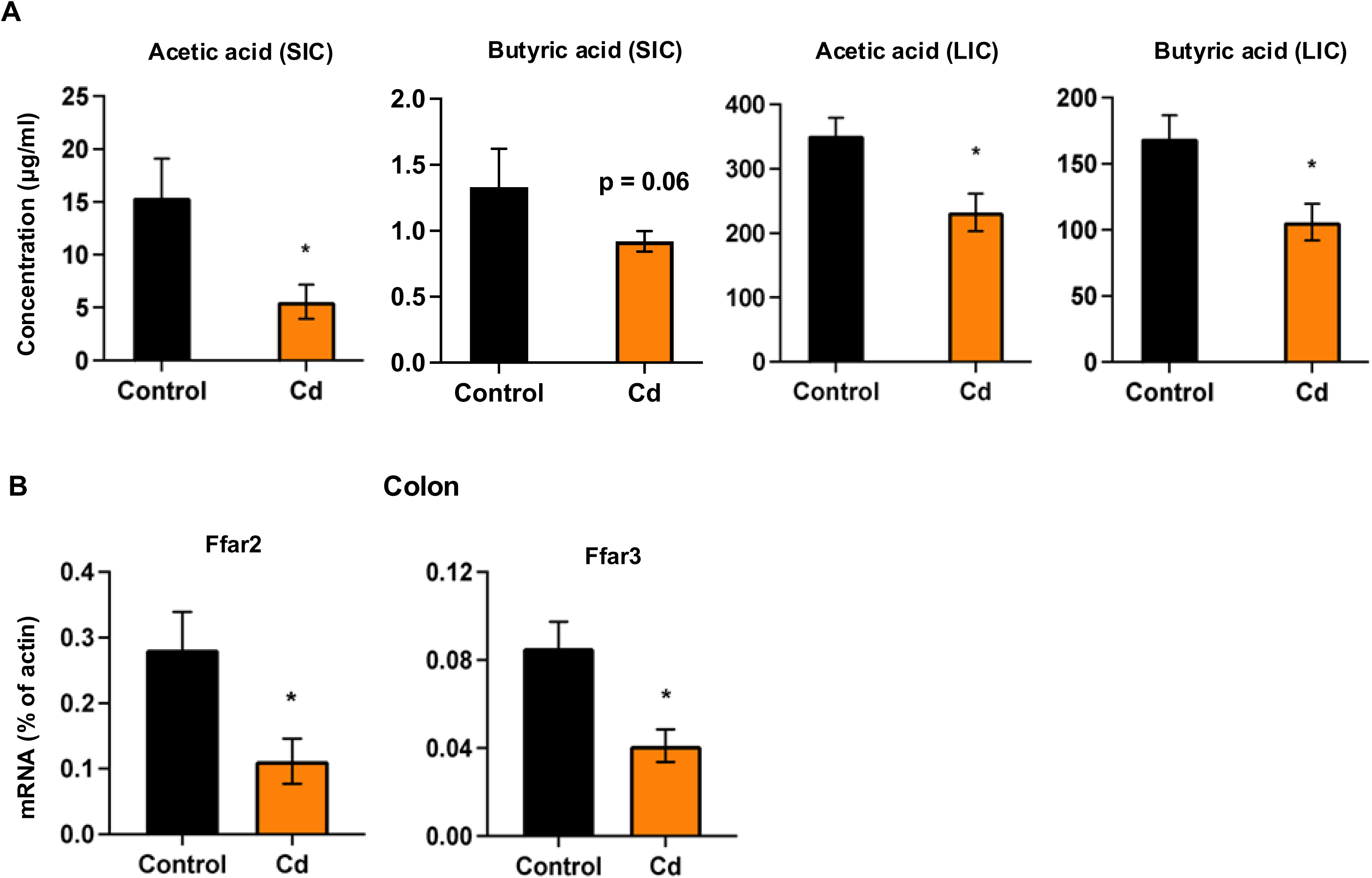
Cd exposure changes the concentrations of microbial metabolites and short-chain fatty acids-related receptors in mice intestines. **A.** The levels of short-chain fatty acids in the intestinal contents. **B.** Cd exposure changes Ffar2, and Ffar3 expression levels in the colon. n=7-8 in each group. * p < 0.05; ** p< 0.01; *** p < 0.001.

**Figure 6.**
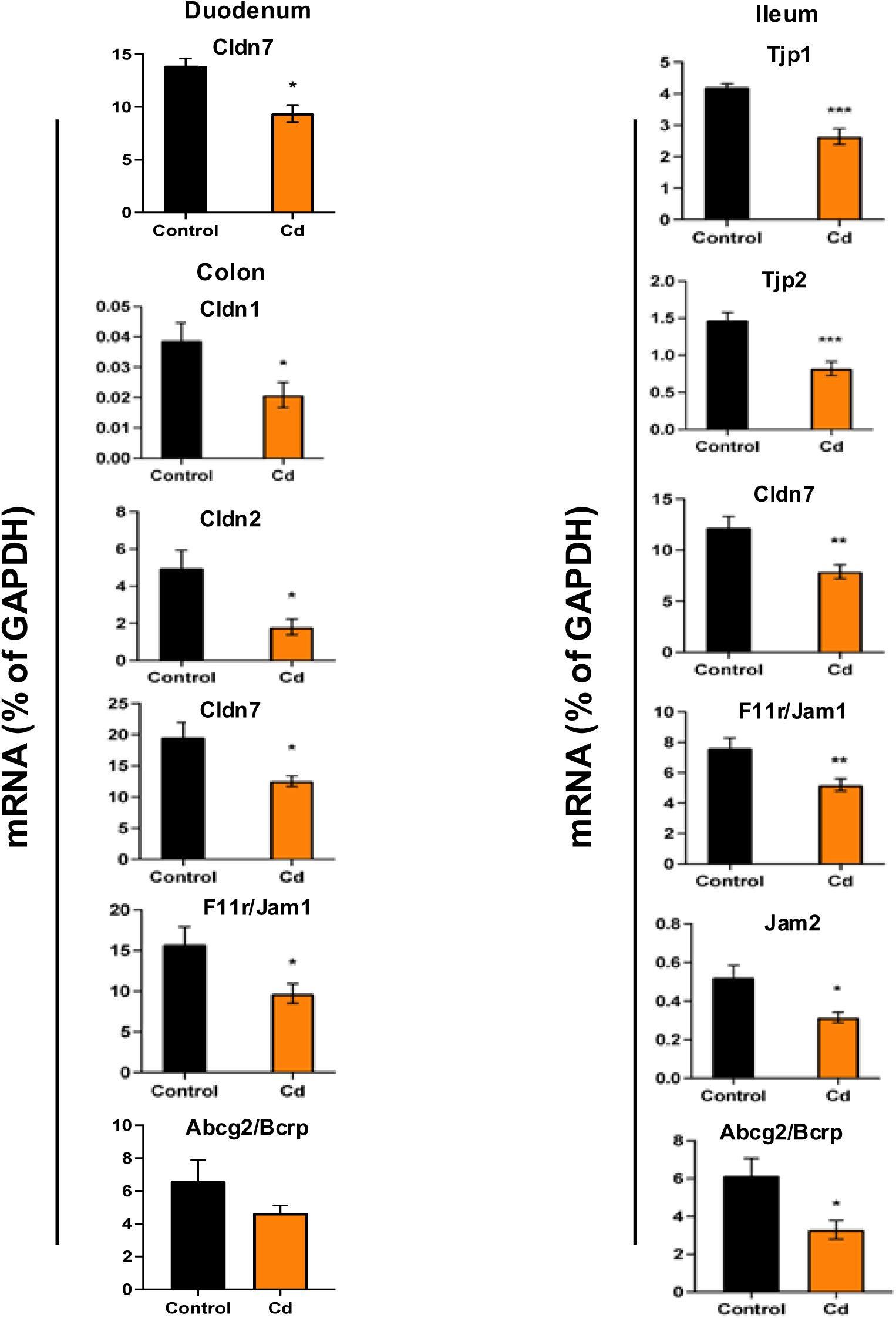
Cd exposure changes the expression of tight junction proteins in the intestine of mice. n=7-8 in each group. * p < 0.05; ** p< 0.01; *** p < 0.001.

**Figure 7.**
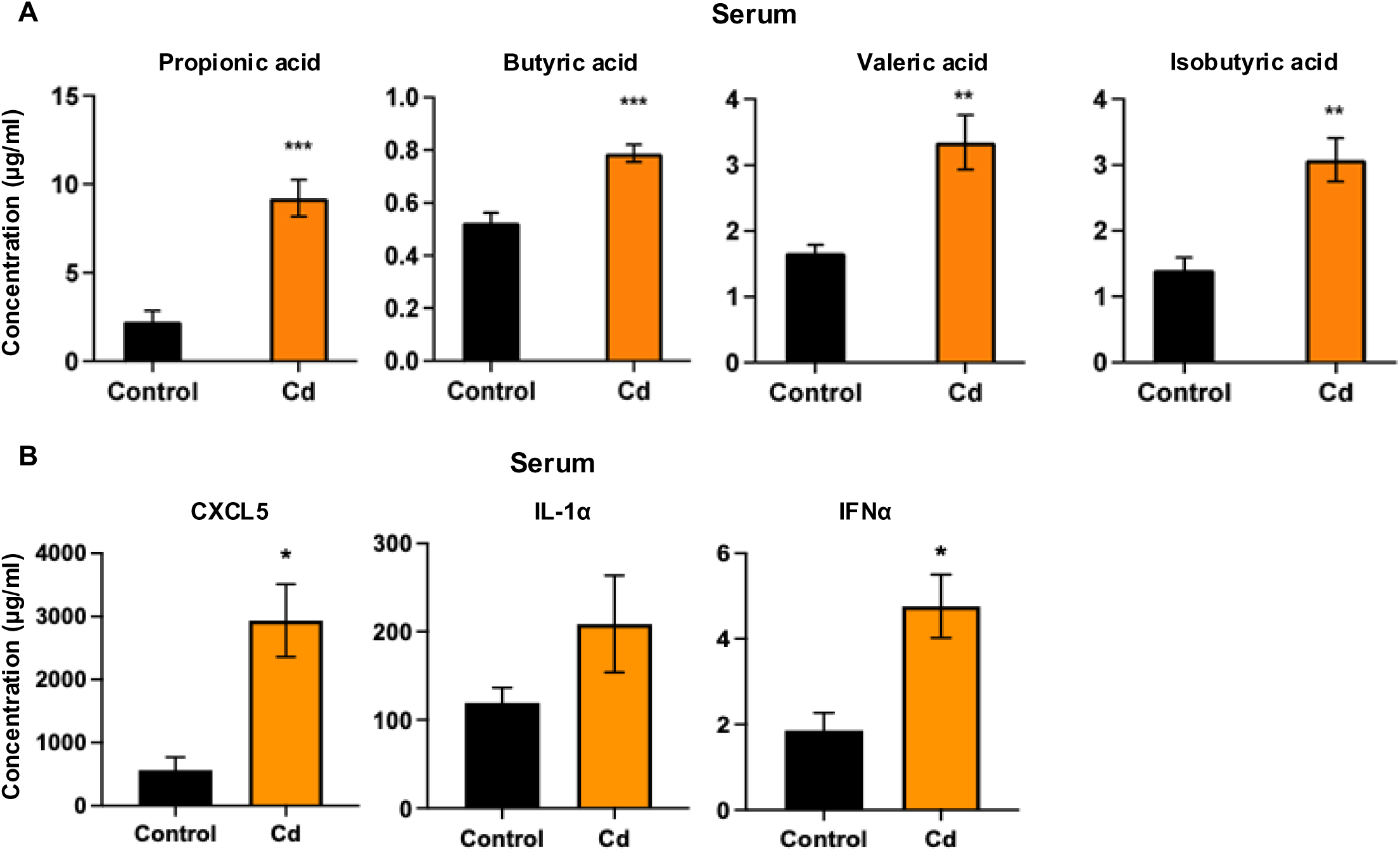
Cd exposure changes the levels of microbial metabolites and inflammatory cytokine levels in mice serum and brain. **A.** The levels of short-chain fatty acids in the serum. **B.** Cd exposure changes the inflammatory cytokines levels in the serum. n=7-8 in each group. * p < 0.05; ** p< 0.01; *** p < 0.001.

Associated with decreased expression of genes involved in gut barrier integrity and BCRP, which are hallmarks of intestinal inflammation ^84,85^, Cd exposure increased the levels of multiple pro-inflammatory cytokines in serum (Fig. 7B), including C-X-C motif Chemokine Ligand 5 (CXCL5), which is known to promote neuroinflammation and blood-brain barrier injury ^86^, as well as Interleukin-1 Alpha (IL-1α), and Interferon Alpha (IFN-α), which are critical components of leukocyte transition/recruitments to promote neuroinflammation ^87,88^.

### 6. Cd exposure induced significant changes in the expression of inflammation-related genes in the hippocampus

Neuroinflammation is a known mechanistic contributor to learning and memory deficits, and can be a result of intestinal inflammation, gut dysbiosis, and gut barrier disruption ^89^. Aligning with the reduced intestinal SCFAs and tight junction transcripts (Fig. 5 and Fig. 6), as well as increased serum cytokines (Fig. 7B), Cd exposure dysregulated 27 inflammation-related genes in the hippocampus (Fig. 8A). For example, Cd up-regulated multiple pro-inflammation-related genes (Fig. 8B): Interleukin-1 Receptor Type 1 (*Il1r1)* is a critical receptor in the interleukin-1 (IL-1) signaling pathway, and Interleukin-1 Receptor Accessory Protein (*il1rap)* serves as an important co-receptor that forms part of the IL-1 receptor complex. IL1R1 binds to the cytokine IL-1α/β, key mediators of inflammation, and upon ligand binding, it forms a complex with IL1RAP to initiate intracellular signaling cascades, including the activation of NF-κB and MAPKs, which lead to the production of cytokines, chemokines, and other inflammatory mediators ^90,91^. Similarly, IL-16 is a pro-inflammatory cytokine that activates immune T cells ^92^ and macrophages ^93^. By recruiting immune cells to inflammation sites and inducing the production of other pro-inflammatory cytokines, IL-16 plays an important role in the inflammatory response ^94^. Lipopolysaccharide Binding Protein (*Lbp*), a soluble acute-phase protein, can bind to bacterial lipopolysaccharide (LPS) and induce immune responses ^95^. Elevated levels of LBP have been associated with neurodegenerative diseases, including AD, PD, and Multiple Sclerosis (MS) ^96–98^. In addition, Cd exposure upregulated the expression levels of complement component 3 (*C3*), a central component of the complement system involved in host defense and inflammation, and C-X-C motif chemokine 5 (*Cxcl5*), a small cytokine from the CXC chemokine family that plays an important role in regulating neuroinflammation ^99–101^.

**Figure 8.**
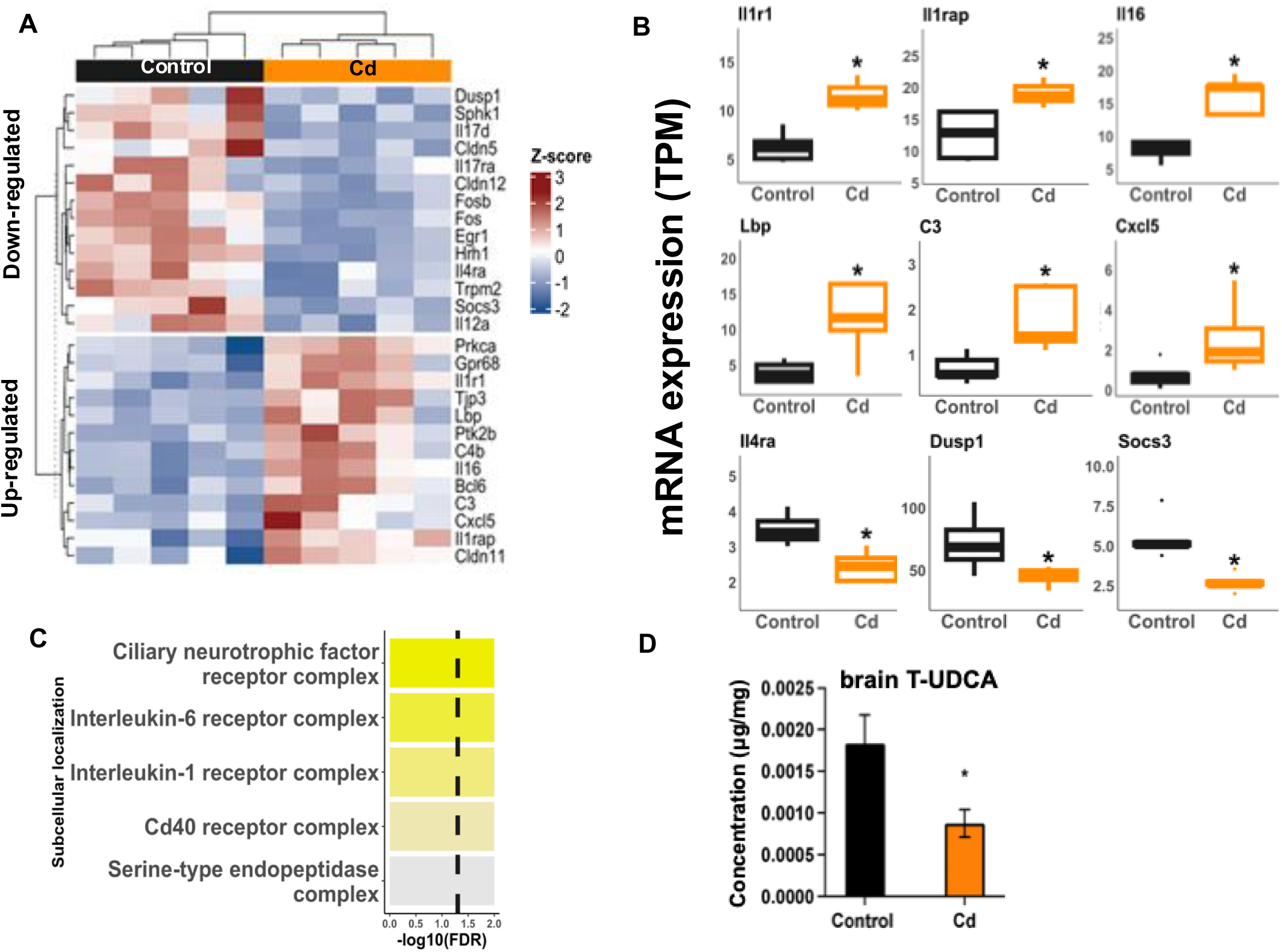
Dysregulated expression signatures of inflammation-related genes in the hippocampus. **A.** Inflammation-related genes that are differentially expressed after 9 weeks of Cd exposure in the hippocampus are shown in a heatmap. The colors of the heatmap represent the log_2_-fold change of hippocampus genes of the Cd-exposed mice as compared with the control group. Red indicates upregulated, and Blue indicates downregulated from Cd-exposed mice (Bonferroni-adjusted p-value < 0.05). **B.** Examples of differentially regulated genes associated with inflammation. Asterisks represent statistically significant differences between Cd and Control groups. n = 5 in each group. * *p* < 0.05. **C.** Pathway analysis of Cd-regulated inflammation related genes in hippocampus. **D.** Cd exposure decreased T-UDCA in the brain. n=7-8 in each group. * *p* < 0.05.

Cd exposure also downregulated multiple anti-inflammation related genes in the hippocampus (Fig. 8B). For example, Cd decreased the mRNA of interleukin-4 receptor Alpha *(Il4rα*), a receptor for Il-4 and Il-13. These cytokines exert anti-inflammatory effects and are known to contribute to cognitive function through their interaction with IL14RA ^102–106^. In addition, Cd exposure downregulated the levels of Dual specificity phosphatase-1 (*Dusp1*), also known as mitogen-activated protein kinase phosphatase-1 (MKP-1). DUSP1 is a key component of the anti-inflammatory response ^106,107^. Cd exposure also affected the expression of Suppressor of Cytokine Signaling 3 (*Socs3*), a member of the SOCS family that plays a pivotal role in regulating JAK-STAT pathways. SOCS3 acts as a negative feedback regulator of cytokine signaling, maintaining immune homeostasis and protecting against excessive inflammation in the CNS ^108^. Pathway analysis confirmed an upregulation of pro-inflammatory signaling in the hippocampus, including pathways associated with the IL-1 and IL-6 receptor complexes as well as the CD40 receptor complex (Fig. 8C). The neuroinflammation signature is associated with a decrease in the neuroprotective metabolite tauroursodeoxycholic acid (T-UDCA) in the brain (Fig. 8D). T-UDCA is a bile acid metabolite produced jointly by the liver and the gut microbiota. To note, T-UDCA is known to inhibit cognitive decline and improves memory with potential mechanisms include reducing inflammation, neuronal apoptosis, and amyloid beta markers ^109^. However, other microbially derived bile acids were not affected by Cd exposure in blood or brain (Fig. S14).

Together, these findings indicate that Cd exposure disrupts the balance of inflammatory and anti-inflammatory gene expression in the hippocampus, associated with leaky gut and pro-inflammatory signatures within the gut-brain axis. This dysregulation may contribute to the Cd-induced neurotoxicity, particularly in hippocampus-dependent learning and memory processes.

### 7. Cd exposure reduced expression levels of brain neuropeptides in mice

Neuropeptides are important mediators within the gut-brain communication^110^. Our findings revealed that Cd exposure reduced the mRNA expression levels of six neuropeptides in the hippocampus of mice, including corticotropin-releasing hormone (*Crh*), natriuretic peptide C (*Nppc*), neuropeptide Y (*Npy*), proenkephalin (*Penk*), thyrotropin-releasing hormone (*Trh*), and vasoactive intestinal peptide (*Vip*) (Fig. 9). These results suggest that Cd exposure can disrupt neuropeptide signaling in the hippocampus, potentially contributing to neurotoxicity.

**Figure 9.**
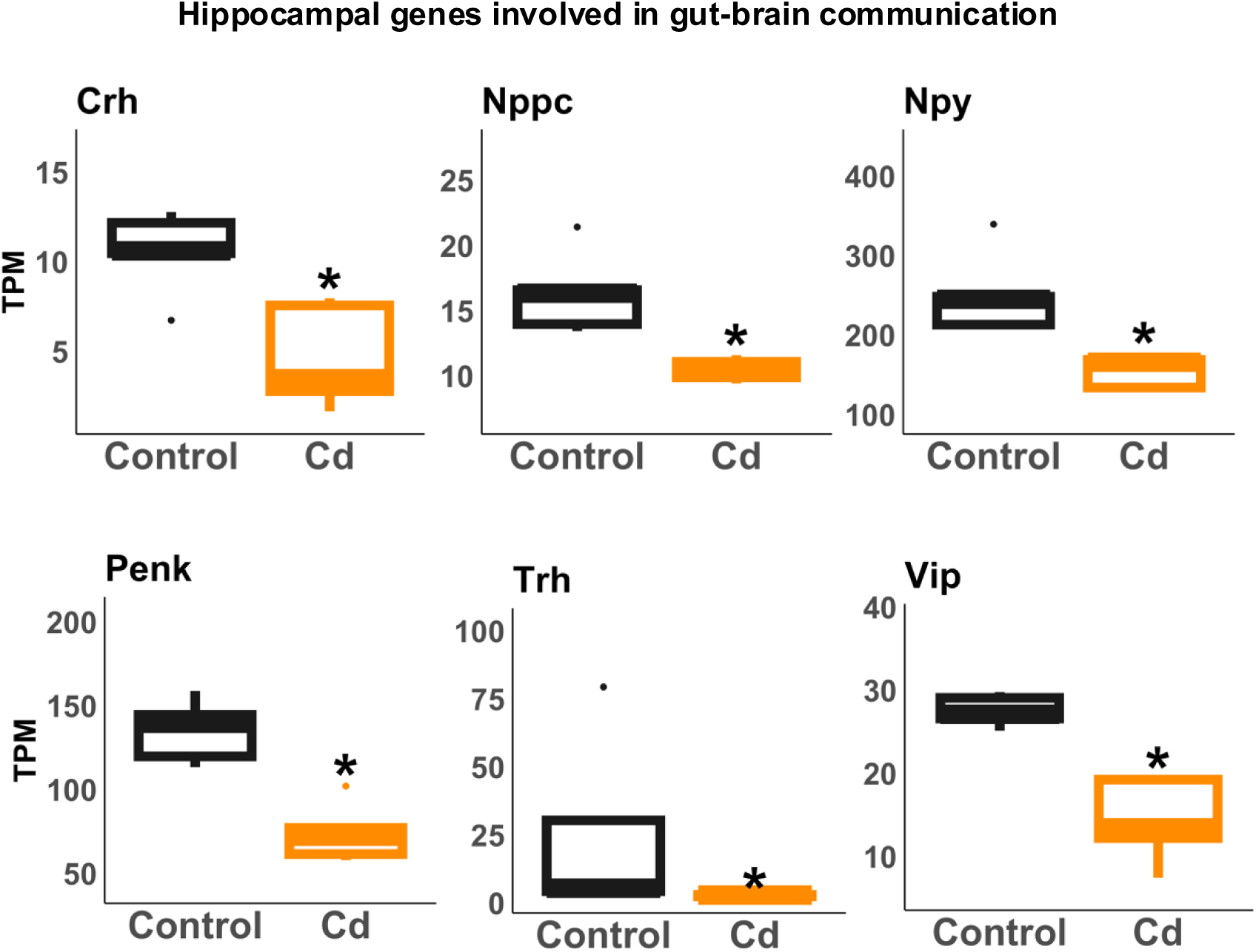
Cd exposure downregulates the expression of neuropeptides in the hippocampus. Examples of differentially regulated genes associated with neuropeptide. Asterisks represent statistically significant differences between Cd and Control groups. n = 5 in each group. *. p < 0.05.

### 8. Cd-induced dysregulation of the gut microbiome are strongly associated with the expression levels of cognitive- and inflammation-related genes in the hippocampus

To further investigate the association between the gut microbiome and Cd-mediated learning and memory deficits, we explored the potential relationship between Cd-regulated microbes and Cd-regulated genes involved in cognition and inflammation, using Spearman’s correlation (Fig. 10).

**Figure 10.**
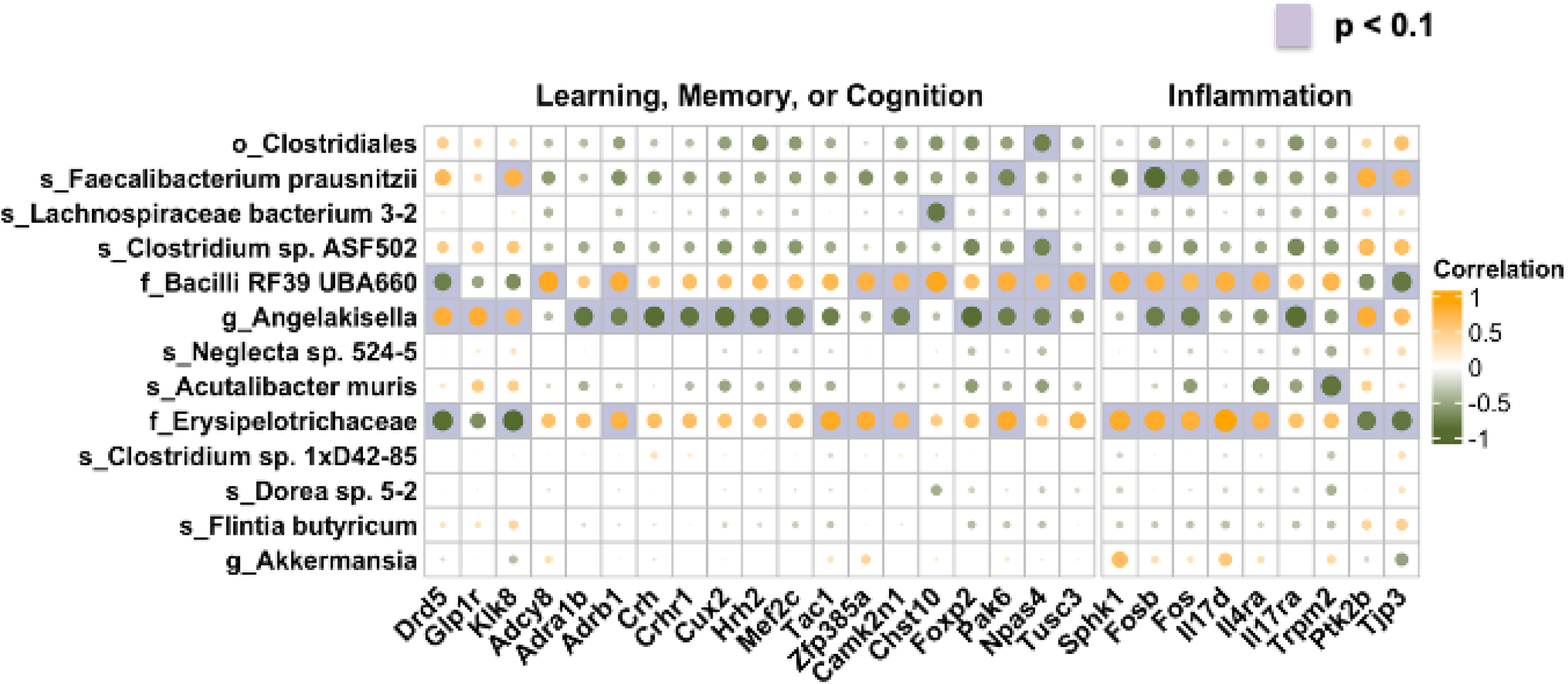
The correlation between the persistently dysregulated gut microbiome and the learning, memory, or cognition-related genes, as well as inflammation-related genes. Yellow and green colors show a positive or negative correlation between specific bacteria and the related gene. The purple square indicates the significant correlation (adjusted p-value < 0.1). n = 5 in each group.

For the cognition-related genes (Fig. 10 left panel), *Clostridiales* was negatively associated with *Npas4. Faecalibacterium prausnitzii* was positively associated with *Glp1r* and negatively associated with *Pak6. Lachnosporaceae bacterium 3-2* was negatively associated with *Chst10. Clostrodium sp. ASF502* was also negatively associated with *Npas4.* UBA660 was positively associated with *Adcy8, Adrb1, Zfp358a, Camk2n1, Chst10, Pak6, Npas4*, and *Tusc3*, and negatively associated with *Drd5*. *Angelakisella* was positively associated with *Drd5, Glp1r, Klk8,* and negatively associated with *Adra1b, Adrb1, Crh, Crhr1, Cux2, Hrh2, Mef2c, Camk2n1, Foxp2, Pak6,* and *Npas4.* In addition, *Erysipelotrichaceae* was positively associated with *Adrb1, Tac1, Zfp385a, Camk2n1,* and *Pak6*, and negatively associated with *Drd5* and *Klk8*.

For the inflammation-related genes (Fig. 10, right panel), *Faecalibacterium prausnitzii* was positively associated with *Ptk2b* and *Tjp3*, and negatively associated with *Fosb and Fos.* UBA660 was positively associated with *Sphk1, Fosb, Fos, Il17d,* and *Il4ra*; and negatively associated with *Tjp3*. *Angelakisella* was positively associated with *Ptk2b,* and negatively associated with *Fosb, Fos, and Il17ra.* Additionally, *Acutalibacter muris* was negatively associated with *Trpm2;* while *Erysipelotrichaceae* was positively associated with *Sphk1, Fosb, Fos, Il17d,* and *Il4ra*, and negatively associated with *Ptk2b* and *Tjp3*.

Together, these findings suggest that the gut microbiome is associated with alterations in the expression of cognition- and inflammation-related genes in the hippocampus. These correlations unveiled distinct microbial biomarkers for Cd-induced neurotoxicity, suggesting the potential influence of Cd-altered gut microbiota in mediating hippocampal dysfunction. The exact molecular mechanisms and causal relationship require further investigations.

## Discussion

Our study is among the first to systematically investigate the effects of Cd on the gut microbiome throughout the entire exposure period, as well as its association with the Cd neurotoxicity on cognitive function. We found that nine weeks of Cd exposure impaired hippocampus-dependent learning and memory and altered the expression of cognition- and inflammation-related genes in the hippocampus of young adult male mice. These changes were associated with Cd-induced gut dysbiosis, characterized by changes in gut microbiome composition, disruptions in SCFA metabolism, as well as reduced expression of genes involved in gut barrier integrity and xenobiotic efflux (*Bcrp*) in the intestine. Furthermore, we observed that gut microbiome alterations preceded the onset of Cd-induced cognitive deficits, with persistently dysregulated bacterial taxa showing strong correlations with cognitive impairments and cognition-associated gene expression in the hippocampus. Importantly, our study is the first to demonstrate that the persistent gut microbiome changes are strongly associated with Cd-induced learning and memory deficits in mice, highlighting a potential gut-brain axis mechanism underlying Cd neurotoxicity.

### Cd exposure induces deficits in hippocampus-dependent learning and memory and significantly alters cognition and neuroinflammation-related genes in the hippocampus

Our previous studies demonstrated that six weeks of exposure to 3mg/L Cd through drinking water can induce hippocampus-dependent learning and memory deficits ^13^. However, the precise time point at which this exposure level causes this deficit remains unclear. We previously showed that the NOL test is a reliable, noninvasive, and sensitive behavioral assay for detecting hippocampus-dependent spatial memory changes at multiple time points ^13,44,111,112^, with memory established typically lasting less than 24 hours and the results of the NOL tests remaining consistent when conducted weekly ^11,12,113^. In this study, we assessed spatial memory weekly using the NOL test, starting from the second week of the exposure period, to determine when deficits emerge. We found that the Cd-exposed group exhibited impaired memory starting at 4 weeks into exposure, and the deficits persisted throughout the remaining exposure period. Cognitive deficits were further confirmed in the contextual fear memory test at week nine into exposure.

Notably, our previous work showed that exposure to 3 mg/L Cd for 5-13 weeks resulted in peak blood Cd levels (2.125-2.25 μg/L), comparable to those observed in smokers in the U.S. (men: 0.58–0.94 μg/L; women: 0.69–1.17 μg/L) and below the standard trigger level of Cd (5 μg/L) for medical surveillance in Occupational Safety & Health Administration (OSHA) regulation. These findings indicate that environmentally relevant Cd exposure (3 mg/L) through drinking water impairs hippocampus-dependent learning and memory in mice, with deficits emerging as early as four weeks into exposure.

To further explore the mechanisms underlying these impairments, we analyzed transcriptional alterations in the hippocampus of mice following nine weeks of Cd exposure. Our results showed that Cd exposure downregulated pathways associated with learning, memory, and cognition, aligning with our behavioral findings. Notably, Cd exposure significantly reduced the expression of the G protein-coupled receptors (GPCRs), including dopamine receptor D1/2, histamine receptor H1/2/3, adrenergic receptor, metabotropic glutamate receptor 4, and probable G-protein coupled receptor 88.

GPCRs are important for synaptic plasticity, and their dysregulation contribute to hippocampus-dependent learning and memory deficits ^114^. For example, dopamine (DA), a key neurotransmitter, has been extensively studied for its important role in regulating learning and memory ^115–117^. DA neurotransmission is mediated by two main classes of DA receptors: the D1 class and the D2 class. The D1 family, which includes high-abundance D1 receptors and low-abundance D5 receptors, positively regulates adenylyl cyclase activity, resulting in increased intracellular cyclic AMP (cAMP) levels. The D2 family, which includes D2/3/4 receptors, inhibits cAMP production ^118^. Studies ^119–121^ have shown that D1 receptor activation can improve spatial working memory in rats, while presynaptic D2 receptor deletion in the hippocampus impairs spatial memory in mice ^122^. Interestingly, we observed an increased expression of D5 receptors in Cd-exposed mice, likely as a compensatory response to decreased expression of D1 receptors. However, given the lower abundance of D5 compared to D1 ^123^, this increase may be insufficient to counteract D1-related impairments.

Histamine receptors are also important for synapse and cognitive functions ^124^. Studies using knockout mice revealed that the absence of H1 and H2 receptor led to hippocampus-dependent learning and memory deficits ^125,126^. In addition, reduced histamine receptor levels and binding capacity have been observed in AD patients ^127^. These findings suggest that Cd exposure disrupts cognitive function by affecting histamine receptor signaling.

In addition to GPCR genes, Cd exposure affected the expression of other cognition-related genes involved in synaptic plasticity. For example, *Tac1* gene encodes the precursor to several neuropeptides in the tachykinin family, including substance P (SP) and neurokinin A (NKA) ^128^. Studies have demonstrated that the administration of SP can improve learning and memory in animal models^129,130^. *Tac1* expression is downregulated in the hippocampus of AD patients ^131^ and AD mouse model ^132^, consistent with our findings. Similarly, Type 8 adenylyl cyclase (ADCY 8), responsible for Ca^2+^-stimulated cAMP production and activity-dependent synaptic modification, is essential for synaptic plasticity, with *Adcy8* knockout mice exhibiting cognitive deficits ^61,133^. *Npas4*, a key regulator of learning-induced plasticity in CA3, is crucial for the formation of contextual fear memory ^62,134^, and its reduced expression has been linked to memory impairments in aged mice ^135^. These findings suggest that Cd-induced transcriptional changes in synaptic plasticity-related genes may contribute to its neurotoxicity on cognition.

Neuroinflammation is recognized as a potential mediator of cognitive deficits ^136^ ^137^, it can impair learning and memory through various mechanisms, including impairing the BBB, inducing oxidative stress, disrupting adult neurogenesis, and inhibiting long-term potential (LTP) ^138,139^. Neuroinflammation is also involved in neurodegenerative diseases, including AD, PD, and MS ^140–142^. Our study showed that Cd exposure increased the blood levels of inflammatory cytokines (CXCL5, IL-1α, and IFNα) and changed the hippocampal expression of inflammation-related genes. For example, Cd exposure upregulated the expression of pro-inflammation genes, such as *Il16* and *Lbp,* both linked to AD risk. Interleukin 16 (*Il16*), a T cell-derived cytokine that activates immune T cells ^92^ and macrophages ^93^, is elevated in AD patients ^143,144^. Lipopolysaccharide

Binding Protein (LBP), a soluble acute-phase protein that binds to bacterial Lipopolysaccharide (LPS) to trigger an immune response ^95^, is associated with a 30% increased likelihood of developing AD ^145^. In addition, increased LBP levels have been observed in patients with other neurodegenerative diseases, such as PD and MS ^96,98^. Complement component 3 (*c3*), a key component of the complement system, is increased in the brains of AD mouse models and patients ^99–101^, PD patients ^146^, and MS mouse models ^147^. In addition, C-X-C motif chemokine 5 (CXCL5), a cytokine involved in astrocyte-mediated neuroinflammation and BBB injury ^86^, has been linked to white matter damage and cognitive decline ^148^. Elevated *Cxcl5* expression in the hippocampus pf elderly individuals suggests its role in brain aging^149^. In the current study, we observed increased *Cxcl5* expression in the hippocampus, aligning with elevated CXCL5 blood levels and cognitive deficits in Cd-exposed mice.

Conversely, Cd exposure downregulated the expression levels of certain anti-inflammation genes, such as *Dusp1* and *Socs3*. Dual specificity phosphatase-1 (*Dusp1*), also known as mitogen-activated protein kinase phosphatase-1 (MKP-1), is a nuclear enzyme that dephosphorylates and inactivates MAP kinases. DUSP1 modulates immune responses ^150^, and its deficiency can increase inflammatory cytokines production ^151^ ^152^. Accumulating evidence suggests a link between DUSP1 and neurodegenerative diseases. For example, reduced *DUSP1* levels have been observed in the hippocampus of AD patients and AD model mice. Additionally, DUSP1 can mitigate oxidative stress and inflammatory injury, ultimately improving spatial learning and memory in epilepsy rats^153^. The suppressor of Cytokine Signaling 3 (*Socs3*), a key anti-inflammatory regulator of JAK-STAT pathways ^108^, modulates anti-inflammatory responses in microglial and macrophage. Increasing evidence suggests that *Socs3* is involved in inflammation-related neurodegenerative diseases ^154^. Studies have reported reduced *Socs3* mRNA levels in the hippocampus of APP/PS1 mice ^155^, while in animal models of MS, SOCS3 was found to exert a protective role ^156^. Together, these findings, along with our results, suggest that neuroinflammation contribute to Cd-induced cognitive deficits in mice.

### Cd-induced gut dysbiosis is associated with learning and memory deficits

Recent studies have increasingly highlighted the causal relationship between the gut microbiome and hippocampus-dependent learning and memory. Animal studies using antibiotics, probiotics, prebiotics, FMT, and other microbiome-modulating techniques have shown that gut microbiota can influence hippocampal structure, synaptic plasticity, and cognition-related behaviors ^157,158^. In humans, prebiotics supplementation has been found to enhance cognitive functions, while antibiotics-induced gut dysbiosis has been linked to the dysfunctions of hippocampus ^23–25,159,160^. Additionally, significant alteration in the gut microbiome have been observed in AD patients and AD mouse models ^16,17,161,162^. Notably, transplanting gut microbiome collected from AD patients into healthy mice induces severe cognitive deficits, which can be rescued by FMT from healthy human donors ^163^. Furthermore, two recent human case reports suggest that FMT can improve memory and cognitive performance in AD patients ^20,21^.

Our research is among the first to examine the effects of Cd on the gut microbiome throughout the entire exposure period. This unique study design provides a comprehensive understanding of gut microbiome changes and the identification of specific bacteria consistently affected by Cd exposure. In this study, we found that Cd exposure induces significant gut microbiome dysregulation in mice, consistent with previous findings that Cd alters the gut microbiome in rodents ^41,43,164–167^. Studies by He et al. ^43^, Yang et al. ^167^, and Li et al.^166^ reported that Cd exposure significantly reduced gut microbial richness (α-diversity) and/or alters microbial community structure (β-diversity). However, we did not observe significant differences in microbial richness or community structure between control and Cd-exposed mice. This discrepancy may be attributed to differences in Cd dosage, exposure duration, or the pre-antibiotic treatment used in their studies. Our exposure level (3 mg/L for 9 weeks) was much lower than in the other studies, suggesting that the exposure levels are critical for the effects of Cd on gut microbiota.

Although Cd exposure did not affect gut microbiome richness or community structure, it induced compositional changes throughout the entire exposure period. Notably, we are the first to report Cd-induced gut dysbiosis before the onset of cognitive deficits, further suggesting that the gut microbiome may contribute to Cd-induced cognitive deficits. Additionally, Spearman correlation analysis identified 21 persistently dysregulated microbiomes associated with the performance of mice in the NOL test (BH adjusted p-value < 0.2). Among these, eight bacterial taxa showed stronger associations (BH adjusted p-value < 0.1), including *UBA660*, *Flavonifractor*, *Erysipelotrichaceae*, *Flintia butyricum*, *Faecalibacterium prausnitzii*, *Proteobacteria*, *Enterobacteriaceae*, and *Escherichia*. In this study, we applied BH adjusted p-value threshold of <0.2 and <0.1, commonly used in exploratory microbiome research to retain potential associations for further validation^72–76,168–170^. This approach allows us to identify microbial taxa with robust correlations to cognitive deficits, providing candidate targets for future mechanistic investigations. Regarding the eight bacterial taxa, Cd exposure persistently downregulated the relative abundance of the family *Erysipelotrichaceae* and the taxon *Bacillales bacterium UBA660*, with these changes occurring before the onset of cognitive deficits. This aligns with previous research showing reduced relative abundances of genera within the same family following Cd exposure^166^. *Erysipelotrichaceae* is an important producer of short-chain fatty acids (SCFAs) such as acetate, butyrate, and propionate ^171^. While its association with cognition is inconsistent, studies have reported lower levels in AD model mice ^172,173^ and mice with cognitive deficitis^174^. In addition, salidroside treatment, which increases *Erysipelotrichaceae* levels, can mitigate memory impairment in rats ^175^. *UBA660*, an unclassified species within the order *Bacillales*, showed the strongest association with mice performance in the NOL test. Though little is known about UBA 660, *Bacillales* is for its protective role against dementia and AD ^176,177, 178^.

Cd exposure upregulated the relative abundance of the species *Faecalibacterium prausnitzii* and *Flintia butyricum,* the genera *Flavonifractor* and *Escherichia*, the family *Enterobacteriaceae*, and the phylum *Proteobacteria.* While the knowledge about *F. butyricum* is very limited, *F. prausnitzii* is one of the most abundant butyrate-producing bacteria in the human gut ^179^. Butyrate has pleiotropic effects on the intestinal cell life cycle and offers numerous beneficial effects, including protecting the intestinal barrier integrity and improving the cognitive function ^180^. Given that Cd exposure significantly reduced intestinal butyrate levels in mice, the increased abundance of *F. prausnitzii* may serve as a compensatory response. However, its relationship with cognition remains unclear. Some studies have reported higher *Faecalibacterium* levels in senior women with subjective memory complaints ^181^ and patients with mild cognitive impairment ^182^. In contrast, other research has found reduced levels in patients with cognitive deficits and AD ^179,181,183^.

*Flavonifractor* is a genus of anaerobic bacteria known for metabolizing flavonoids^184^, a class of polyphenolic secondary metabolites with antioxidant and anti-inflammatory properties^185^. *Flavonifractor* levels are negatively correlated with SCFAs, which play an important role in neuroprotection and are associated with different neurodegenerative diseases ^186^. Its prevalence is significantly higher in patients with AD and bipolar disorder (BD) ^184,187,188^. A recent epidemiological study also identified an association between elevated *Flavonifractor* levels and poorer executive function in senior adults ^183^. These findings align with our results, which show an increased abundance of *Flavonifractor* in Cd-exposed mice with cognitive deficits.

*Proteobacteria, Enterobacteriaceae, and Escherichia* all belong to the phylum *Proteobacteria.* In this study, their relative abundance increased in the Cd-exposed group, exhibiting a similar upregulation trend. *Proteobacteria* is one of the most dominant phyla in the human gut microbiota and is increasingly recognized as a potential microbial biomarker for various diseases, including neurodegenerative disorders ^189^. An epidemiological study found that AD patients with amnestic mild cognitive impairment had enriched levels of *Proteobacteria* in China ^190^. Similarly, higher abundances of *Proteobacteria* have been observed in senior adults with mild cognitive impairments (MCI) ^191^ and individuals with higher subjective memory complaint (SMCs) scores^192^. Animal studies further support these findings, showing an elevated abundance of *Proteobacteria* in AD mouse models ^193,194^ *Enterobacteriaceae*, a large family within the *Proteobacteria* phylum, includes many pathogens, such as *Enterobacter*, *Citrobacter*, *Escherichia coli*, and *Shigella*. This bacterial family is a major cause of various infections in humans ^195^. In the same Chinese cohort ^190^, AD patients with higher SMC scores exhibited increased levels of *Enterobacteriaceae*. Additionally, a meta-analysis study revealed an enrichment of *Enterobacteriaceae* in the gut microbiota of individuals with MCI and AD ^196^. *Escherichia* is considered a conditional pathogen associated with systemic inflammation ^197^. Numerous studies have reported an increased abundance of *Escherichia/shigella* in individuals with learning and memory deficits ^198–200^. Among these, *Escherichia Coli* is particularly notable for its role in disrupting gut integrity, promoting systemic and neuroinflammation, and contributing to cognitive deficits ^201–204^. However, in this study, we did not observe significant changes in the relative abundance of *E. coli* in the Cd group, suggesting that other unidentified species within the genus *Escherichia* may be involved in Cd neurotoxicity.

Furthermore, we observed a reduction in the genus *Akkermansia* and an increase in the genus *Angelakisella* and the order *Clostridiales* in Cd-exposed mice. Previous studies have linked these three bacterial taxa to cognitive function. In this study, their relative abundance also significantly correlated with behavioral performance of the mice in cognitive tests (BH adjusted p-value < 0.2).

*Akkermansia* is a mucin-degrading bacteria found in the human gut microbiota. Studies have found an association between *Akkermansia* and various neurodegenerative diseases. For example, its abundance decreases in ALS mouse model ^205^. Animal studies have also reported a negative correlation between *Akkermansia* and pathogenic Aβ42 levels in the brain ^206^. Moreover, *Akkermansia* is positively associated with cognitive function in adults in the U.S. ^69^. Dietary interventions, such as ketogenic or Mediterranean ketogenic diet, which increase *Akkermansia* abundance, have been shown to help prevent cognitive decline ^67,70,191^. In our study, Cd exposure significantly reduced *Akkermansia*, potentially contributing to Cd-induced cognitive deficits. Interestingly, *Akkermansia (A) muciniphila,* the most well-studied species within the *Akkermansia* genus with cognitive benefits ^207^, was not affected by Cd exposure, suggesting that other *Akkermansia* species may also support cognitive function. *Angelakisella*, a genus within the family *Eggerthellaceae*, is also associated with cognitive function. A higher abundance of *Angelakisella* has been observed in type 1 diabetic mice with cognitive deficits compared to healthy controls ^71^. Additionally, *Angelakisella* has been associated with poor cognitive performance in aging people with insomnia ^66^. The order *Clostridiales*, a large and diverse group within the gut microbiota, plays an important role in the gut-brain axis. Several studies have reported a positive association between *Clostridiales* abundance and cognitive performance^190,208–210^. However, other studies reported that higher *Clostridiales* is associated with autism ^65,211^.

Furthermore, an elevated proportion of *Clostridiales* in the gut microbiome is associated with poor cognitive function in mice fed a high-fat/high-sucrose diet ^68^, which aligns with our findings that Cd exposure led to increased *Clostridiales* levels in mice with cognitive deficits.

### Cd-induced gut microbiome alterations are associated with the expression levels of cognition related genes in the hippocampus

Our analysis revealed significant associations between the gut microbiome and differentially regulated genes related to learning, memory, and cognition after 9 weeks of Cd exposure. Notably, many of these genes are involved in synaptic plasticity modulation. For example, *F. prausnitzii* was positively associated with *Klk8* and negatively with *Pak6*, while *Angelakisella* positively associated with *Klk8* and negatively with *Adra1b/b1*, *Camk2n1, Pak6, and Npas4.* Additionally, *UBA660* and *Erysipelotrichaceae* were associated with *Klk8, Pak6, Tac1, Camk2n1,* and *Npas4*.

Among these genes, Kallikrein 8 (*Klk8*), a serine protease of the kallikrein family, is an important modulator of synaptic plasticity^212^. Elevated *Klk8* levels have been associated with cognitive deficits in both AD mouse models and patients^213^ ^214,215^. PAK6, a member of the p21 activated kinase (PAK) family of Ser/Thr kinases involved in cytoskeletal organization, is critical for the regulation of neural plasticity ^216,217^. Previous study has found that *Pak5* /*Pak6* double-knockout mice exhibit spatial learning and memory deficits ^218^. In addition, *Camk2n1* encodes proteins that function as inhibitors of calcium/calmodulin-dependent kinase II (CaMKII), a key enzyme for long-term potential (LTP) and memory formation ^219,220^. By inhibiting CaMKII activation, *Camk2n1* regulates the synaptic plasticity and contribute to LTP induction and consolidation ^221^.

As previously discussed, *Adra1b/b1, Npas4,* and *Tac1* are also key genes involved in synaptic plasticity. Collectively, these findings suggest the gut microbiome may affect learning and memory by impairing synaptic plasticity in mice.

Zinc finger protein 385A (*Zfp385a*) is a transcription factor involved in neuronal development, synaptic plasticity, and cognitive functions. *Lijima et al.* ^222^ first showed that *Zfp385a* protein can regulate dendritic localization and BDNF-induced translation of type 1 inositol 1,4,5-trisphosphate receptor (IP3RI) mRNA in neurons, indicating its important role in regulating the synaptic plasticity. In addition, an epidemiological study found that higher ZNF385A expression is associated with slower cognitive decline in humans ^223^, highlighting a potential protective role of this gene in cognition. In this study, we found that Cd exposure downregulated *Zfp385a* expression, with bacterial taxa *UBA60* and *Erysipelotrichaceae* showing significantly positive correlations with its expression.

In addition, Carbohydrate Sulfotransferase 10 (*Chst10*), also known as Human natural killer-1 sulfotransferase (HNK-1ST), is an enzyme that catalyzes the sulfation of specific carbohydrates in glycoproteins and glycolipids, playing an important role in cognitive function. *Senn et al.* ^224^ first reported that *HNK-1ST* knockout mice exhibited spatial learning and memory deficits. Similarly, another study found mice lacking HNK-1 in the brain exhibited impaired LTP and spatial learning and memory ^225^. Beyond its role in cognition, *Chst10* is also involved in neurogenesis. A study using siRNA knockdown demonstrated that silencing CHST10 reduces neural stem cell proliferation ^226^, suggesting its importance in neural development. In the present study, we found *UBA660* is positively correlated with Chst10, while *Lachnospiraceae bacterium 3-2* showed a negatively correlation.

Furthermore, *Angelakisella* was negatively associated with neurogenesis-related genes, including *Cux2, Mef2c*, *Foxp2,* and *Fosb, while UBA660 and Erysipelotrichaceae* were positively associated with *Fosb*. These genes encode transcription factors that regulate the proliferation, differentiation, and maturation of the neural progenitor cells in the brain ^227–230^. Adult neurogenesis, a process of generating functional new neurons from the adult neural progenitor cells, plays a critical role in hippocampus-dependent learning and memory. Our previous studies have shown that Cd exposure can impair learning and memory by disrupting the normal process of adult neurogenesis ^12,113,231^. In addition, accumulating evidence suggests that the gut microbiome can influence adult neurogenesis ^232–234^. Our findings align with previous research, indicating that gut microbiome alterations may contribute to Cd-induced cognitive deficits by impairing adult neurogenesis.

### Cd-induced gut microbiome alterations are associated with the expression levels of neuroinflammation-related genes in the hippocampus

Neuroinflammation is another important mechanism contributing to Cd neurotoxicity ^235–237^. Our study identified significant correlations between the specific bacterial taxa and inflammation-related genes in the hippocampus. Specifically, *UBA660 and Erysipelotrichaceae* were positively correlated with *Sphk1, Il4ra*, and *Il17d*, while *Erysipelotrichaceae* was negatively associated *with Ptk2b.* In addition, *F. prausnitzii* and *Angelakisella* were positively correlated with *Ptk2b*.

Sphingosine kinases 1 *(Sphk1)* plays an important role in mediating neuroinflammation by converting neuroprotective sphingosine to sphingosine-1-phosphate (S1P) ^238,239^. Reduced SPHK1 activity and protein expression has been observed in the hippocampus of AD patients, suggesting its involvement in learning and memory deficits ^238^. *Ptk2b*, which encodes the non-receptor tyrosine kinase (PYK2), regulates neuroinflammation ^240,241^ and has been identified as a genetic risk factor of late-onset AD (LOAD) ^242^. Inhibition of PYK2 can enhance microglial activity and improve phagocytosis, aiding in the removal of Aβ oligomers both *in vitro* and *in vivo* ^243–245^. *Il14ra* and *Il17d* are also implicated in cognitive impairments. Spatial memory deficits have been observed in mice deficient in IL-4, IL-13, or IL-4 receptor α (*Il4ra*) ^102,246,247^. Deletion of IL-4Rα in GABAergic neurons induces contextual fear memory deficits ^104^. Furthermore, IL-17D (*Il17d*), a member of the IL-17 family, is a potent pro-inflammatory cytokine strongly associated with neuroinflammation ^248^. IL-17 has been associated with neurogenesis, glial function, and BBB integrity, all of which contribute to learning and memory ^249^. *Ribeiro et al*. reported that IL-17 deficiency can impair spatial short-term memory, while γδ T cells, a major source of IL-17, enhance short-term memory by increasing glutamatergic synaptic plasticity in hippocampal neurons ^104,250^.

Consistent with our findings, IL-17 is suggested to mediate inflammatory signals via the microbiome-gut-brain axis. Various studies have reported associations between the gut microbiome and IL-17, IL-17 receptor, as well as Th17 cells ^249^. Our observation that Cd exposure increased the levels of inflammatory cytokines provides further evidence supporting this mechanism. Overall, our findings suggest that the gut microbiome alterations may contribute to Cd-induced cognitive deficits by disturbing the balance of neuroinflammation in mice.

### SCFAs may be a key factor in modulating Cd neurotoxicity through the gut-brain axis

SCFAs are neuroactive microbial metabolites that play a crucial role in regulating cognitive functions ^26^. In this study, we found that Cd exposure significantly reduced intestinal levels of acetic and butyric acids, while increasing serum levels of propionic acid, butyric acid, valeric acid, and isobutyric acid. We also observed a persistent decrease in the relative abundance of many acetate- and butyrate-producing bacteria throughout the exposure period, including *Erysipelotrichaceae*, *Flavonifractor plautii*, *Akkermanisa*, *Ruminococcus*, *Streptococcus*, and *Peptococcaceae* ^251–255^, which explains the reduction in intestinal acetic and butyric acid levels in Cd-exposed mice. Interestingly, we also observed an increase in *F. prausnitzii*, one of the most abundant butyrate-producing bacteria in the gastrointestinal tract ^179^. Given that Cd exposure significantly reduced the intestinal butyric acid levels, the increased abundance of *F. prausnitzii* may represent a compensatory response.

Acetic and butyric acids, the two major SCFAs produced by the gut microbiome, play beneficial roles in modulating learning and memory via the gut-brain axis by activating enteric and central nervous systems, mediating enteroendocrine signaling, as well as regulating epigenetic signaling, neuroinflammation, and the integrity of BBB ^180^. Furthermore, reduced acetic and butyric acids levels have been associated with neurodegenerative diseases. For example, reduced acetic acid levels have been observed in AD disease model in Drosophila ^256^, while decreased butyric acid levels have been reported in AD mouse models ^18^. Similarly, reductions in these SCFAs have been observed in the feces of patients with MCI, AD, and PD ^191,257,258^. These findings are consistent with our results, suggesting that Cd exposure impairs learning and memory by disrupting the balance of SCFA-producing bacteria.

The intestinal barrier plays a critical role in maintaining gut homeostasis and preventing the translocation of pathogens and their products to other organs, including the brain ^30^. SCFAs, particularly butyrate and acetate, are the most important components for maintaining intestinal barrier integrity ^77^. Butyric acid supports the intestinal epithelium by upregulating genes encoding tight junction components, stabilizing transcription factors that coordinate barrier protection, and exerting anti-inflammatory effects on lymphoid and CD4^+^ T cells ^259–262^. Acetic and butyric acids also can stimulate mucin secretion, which protects the intestinal epithelial cells ^263^. Consistent with the reduced levels of acetic and butyric acids in intestine contents following Cd exposure, we observed a significant decrease in the mRNA expression of tight junction and junctional adhesion proteins (*Cldn1/2/7, Tjp1/2, and JAM1/2*) across three different sections of the intestine (Fig.5).

A disrupted intestinal barrier allows pathogenic microorganisms and their toxic products to enter circulation and reach the brain, triggering inflammatory and neurogenic responses that contribute to neurological disorders^30^. Numerous studies have linked the disruption of intestinal barrier to neurodegenerative diseases in humans, including AD ^37^, PD ^264^, and MS ^265^. Animal studies further demonstrated that alleviating intestinal barrier impairment using probiotics can delay neuropathological changes and improve cognitive function ^266–268^. These findings, along with our results, suggest that intestinal barrier disruption may contributes to Cd-induced cognitive deficits, with SCFAs alterations being a key contributing factor.

Moreover, Cd-induced intestinal barrier impairment may explain the divergent SCFAs trends observed between serum and intestinal content. While SCFAs levels in intestinal contents declined (e.g., acetic and butyric acids), a leaky gut may facilitate the direct translocation of these SCFAs into the bloodstream, bypassing normal intestinal epithelial absorption and leading to elevated circulating SCFAs levels. The unchanged expression levels of SCFA transporters (Fig. S13) further suggests that a leaky gut may contribute to the increased serum SCFA levels.

Recent studies indicate that the SCFA butyrate is a substrate for ABCG2/BCRP-mediated efflux in intestinal cells ^82,83^. Because BCRP exports butyrate into the intestinal lumen, its downregulation may also contribute to increased serum butyric acid levels. The Cd-induced reduction of *Bcrp* may result in decreased butyric acid secretion into intestinal lumen, which exacerbates inflammation and leaky gut. This detrimental effect may outweigh the protective effects of elevated serum SCFAs (Fig. 7A), ultimately associated with in CNS toxicity rather than protection.

SCFAs can affect learning and memory via the gut-brain axis by modulating the secretion of gut hormones, such as neuropeptides and neurotransmitters ^27^. Neuropeptides are small, protein-like molecules that function as chemical messengers in the nervous system and other tissues, including the GI tract. These molecules can be produced by central, peripheral, and enteric neurons, as well as by enteroendocrine cells in the GI tract., SCFAs exert their effects by biding to FFAR2 (GPR41) and FFAR3 (GPR43), thereby inducing the secretion of neuropeptides, such as Peptide YY (PYY) and glucagon-like peptide 1/2 (GLP1/2), from enteroendocrine L cells in the colon ^269–271^. These neuropeptides are then transported to the brain via vagal afferents or circulating blood, where they can influence neuroplasticity, memory, and cognitive function ^180^. In addition, SCFAs utilize the neuropeptides released from enteroendocrine cells as secondary endocrine messengers to regulate gastric motility, glucose homeostasis, behavior, and the gut microbiome, ultimately affecting the secretion of cerebral neuropeptides, including neuropeptide Y (NPY), corticotropin-releasing hormone (CRH), and vasoactive intestinal peptide (VIP) ^272^.

NPY is one of the most abundant neuropeptides in the brain ^270^, and plasma NPY can cross the BBB, indicating that brain NPY may also originate from peripheral sources^273^. Studies have shown that NPY and its receptors are important for learning and memory ^274^. For example, central NPY treatment can prevent Aβ-induced spatial memory deficits ^275^, and reduced hippocampal NPY expression has been linked to impaired neurogenesis and cognitive deficits in aged mice^276^. Similarly, decreased NPY levels have been observed in hippocampus, plasma, and cerebrospinal fluid (CSF) in AD patients ^277^. CRH, another important neuropeptide for learning and memory, is beneficial for hippocampus-dependent learning and memory by improving the synaptic plasticity ^278,279^. VIP, known for its neurotrophic and neuroprotective properties, is widely expressed in both the peripheral and central nervous systems and plays an important role in cognitive functions^128^. Notably, VIP has been implicated in the pathophysiology of neurodegenerative diseases, including AD and PD ^280^. Due to its anti-inflammatory and neuroprotective properties, VIP has been proposed as a promising therapeutic target for AD treatment ^281^. Consistent with these findings, our study found significantly reduced intestinal levels of acetic and butyric acids, as along with the expression of their receptors (FFAR2/3) in the colon. More importantly, the expression of various neuropeptides, including NPY, CRH, NPPC, PENK, TRH, and VIP, was reduced in the hippocampus of Cd-exposed group. These results together suggest that the SCFAs-modulated neuropeptide alterations may also contribute to Cd-induced learning and memory deficits.

Neuroinflammation is another potential mechanism by which SCFAs contribute to Cd-induced cognitive deficits. SCFAs modulate the differentiation, maturation, and activation of different immune cells, including neutrophils, dendritic cells, macrophages, T cells and monocytes^282^. Studies have shown that acetic and butyric acids can inhibit the maturation and activation of immune cells and reduce pro-inflammatory cytokine expression^282–284^. In addition, butyric acid can enhance intestinal barrier integrity and suppress LPS-induced inflammation in rat primary microglia, hippocampal slice culture, and co-culture of rat granule neurons, astrocytes and microglia cells ^285^. More importantly, epidemiological, genetic, and descriptive research using brain tissue from patients and animal models strongly suggest that neuroinflammation plays a critical role in the neurodegenerative process and cognitive decline ^286^. In our study, Cd exposure significantly reduced intestinal acetic and butyric acid levels while increasing serum propionic acid – a SCFA associated with neuroinflammation in animal brains and elevated in AD animal models and patients. In addition, the downregulation of tight junction proteins and the *Abcg2* in the intestine, both strongly indicative of intestinal inflammation ^287^, along with significant alterations in hippocampal inflammatory genes and elevated serum inflammatory cytokines, further suggest that SCFAs and their influence on inflammation play a significant role in modulating Cd-induced cognitive deficits.

### Alteration in T-UDCA may also contribute to Cd-induced learning and memory impairments

In the current study, we found that Cd exposure significantly reduced the T-UDCA levels in the brain. Numerous studies have suggested that T-UDCA, a taurine conjugate of ursodeoxycholic acid (UDCA), exerts neuroprotective effects in various neurodegenerative disorders models, including AD, PD, and Huntington’s disease, both *in vitro* and *in vivo* ^288^. T-UDCA is known to protect neurons by inhibiting apoptosis, preventing oxidative stress, and reducing inflammation^289^. These findings, together with our results, suggest that the reduced levels of T-UDCA may contribute to Cd-induced learning and memory deficits in mice.

However, the mechanisms leading to reduced T-UDCA levels in the brains of Cd-treated mice remain unclear. Notably, there is no difference in blood T-UDCA levels between the control and Cd-exposed group (Fig. S14). One potential explanation for the decreased T-UDCA levels in the brain is a reduction in the expression of these T-UDCA transporters at the BBB. In addition, previous studies suggest that BA levels in the brain are approximately 10-fold higher than circulating levels under normal conditions, indicating that BAs can also be synthesized in the brain^290^. This raises the possibility that Cd exposure may inhibit the synthesis of T-UDCA in the brain, leading to its lower concentration in this region. Future studies should investigate the expression levels of T-UDCA transporters at the BBB and measure the concentrations of brain-synthesized T-UDCA to clarify these mechanisms.

### Limitation of the study and future direction

Our study has a few limitations. First, although our correlation analysis identified significant correlations between gut microbiome and cognitive deficits, as well as cognition- and inflammation-related genes expression in the hippocampus, the causal relationship between Cd-induced gut dysbiosis and learning and memory deficits remains unclear. Cd is known to exert neurotoxic effects through multiple direct mechanisms, including oxidative stress, neuronal cell death, neuroinflammation, neurotransmitter disruption, and BBB disruption ^14^. However, the extent to which gut microbiome disruption contributes to Cd-induced neurotoxicity on learning and memory is still not fully understood. In addition, while we observed that Cd exposure impaired intestinal barrier integrity and altered SCFAs and secondary BAs levels in both the intestine and brain, it remains unclear whether these effects directly from Cd toxicity or are secondary to Cd-induced gut microbiome alterations. To address these uncertainties, future studies using germ-free mice or antibiotics-treated mice to Cd will be essential to determine the causal relationship between the gut microbiome and Cd-induced cognitive impairments.

Furthermore, our research suggests that SCFAs and secondary BAs (T-UDCA) may contribute to the effects of Cd on learning and memory. To confirm their significance, supplementary studies, both *in vitro* and *in vivo*, are needed in the future to investigate the potential rescue or protective effects of SCFAs and secondary BAs supplementation in Cd-exposed neuronal cells and mice, as well as to elucidate the underlying molecular and cellular mechanisms.

Another limitation of our current study is the exclusive use of male mice, which prevents us from assessing potential sex-specific differences in how the gut microbiome may modulate Cd-induced neurotoxicity on learning and memory. Increasing evidence highlights sex as a critical biological variable in toxicological responses, including those to Cd. For example, several epidemiological studies have reported stronger association between Cd exposure and neurological impairments, such as neurobehavioral development deficits ^291^and mental health problem ^292^ in females. Animal studies have also demonstrated sex-dependent effects of Cd neurotoxicity. Lamtai et al. ^293^ found that chronic Cd exposure induced significant anxiety-like behavior and memory deficits in rats, with males appearing more susceptible than females. Similarly, our previous study showed that male *ApoE3/E4* mice exhibited greater vulnerability to Cd-induced cognitive deficits than females^11^.

Furthermore, sex-dependent effects of Cd exposure have also been observed in the gut microbiome. One study reported that early-life Cd exposure induced significant sex-dependent effects on gut microbiome in mice. Additionally, in ApoE4-KI mice, Cd-induced gut dysbiosis was more pronounced in males than females^294^.

Together, these findings underscore the importance of considering sex as an independent variable in future research. Evaluating both male and female animals will help clarify whether sex-dependent mechanisms influence the role of the gut-brain axis in modulating Cd neurotoxicity.

## Conclusion

In summary, our study is among the first to provide evidence that Cd can produce persistent alterations in the gut microbiome composition and microbial metabolites (Fig. 11). Notably, we identified significant gut dysbiosis occurring prior to the onset of hippocampus-dependent learning and memory deficits and discovered significant correlations between Cd-induced microbiome changes and cognitive deficits. Furthermore, we observed a significant reduction in the levels of acetate and butyrate in the intestine and TUDCA in the brain, suggesting that these microbial metabolites may play important roles in modulating Cd neurotoxicity on learning and memory. Although our findings are observational, they provide new insights into the mechanisms underlying Cd neurotoxicity and lay the groundwork for future research.

**Figure 11.**
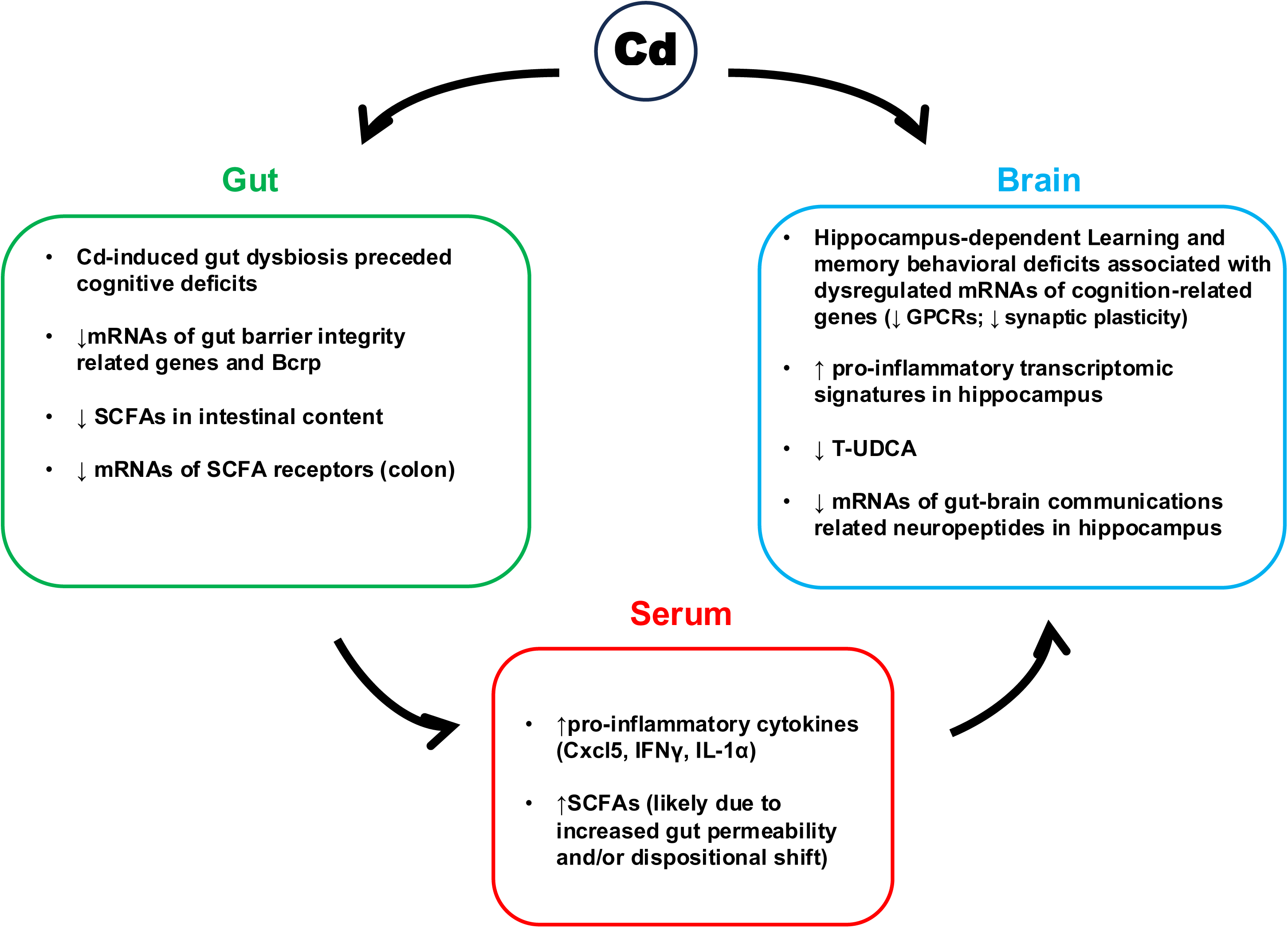
Summary of the key findings. Summary and working hypothesis regarding how gut-brain axis may contribute to the Cd-induced hippocampus-dependent learning and memory deficits in mice.

## Acknowledgments

This research was supported by the National Institute of Environmental Health Sciences [R00ES034068 (to Dr. Wang); University of Michigan Lifestage Environmental Exposures and Disease Center (NIH P30ES017885); National Institute of Diabetes and Digestive and Kidney Disease 1F31DK139707 (to Lim); R01ES025708, R01ES 030197, R01ES031098 (to Dr. Cui)]; University of Washington Environmental Health and Microbiome Research Center (EHMBRACE), as well as University of Washington Center for Exposures, Diseases, Genomics, and Environment (NIH P30ES0007033). The authors would also like to thank the members of Dr. Wang’s and Dr. Cui’s laboratories for their help in tissue collection and manuscript revision.

## Reference

1 Wang, M. et al. A review on Cadmium Exposure in the Population and Intervention Strategies Against Cadmium Toxicity. Bull Environ Contam Toxicol 106, 65–74, doi:10.1007/s00128-020-03088-1 (2021).

2 Genchi, G., Sinicropi, M. S., Lauria, G., Carocci, A. & Catalano, A. The Effects of Cadmium Toxicity. Int J Environ Res Public Health 17, doi:10.3390/ijerph17113782 (2020).

3 Charkiewicz, A. E., Omeljaniuk, W. J., Nowak, K., Garley, M. & Niklinski, J. Cadmium Toxicity and Health Effects-A Brief Summary. Molecules 28, doi:10.3390/molecules28186620 (2023).

4 Wang, B. & Du, Y. Cadmium and its neurotoxic effects. Oxid Med Cell Longev 2013, 898034, doi:10.1155/2013/898034 (2013).

5 Branca, J. J. V., Morucci, G. & Pacini, A. Cadmium-induced neurotoxicity: still much ado. Neural Regen Res 13, 1879–1882, doi:10.4103/1673-5374.239434 (2018).

6 Peng, Q., Bakulski, K. M., Nan, B. & Park, S. K. Cadmium and Alzheimer’s disease mortality in U.S. adults: Updated evidence with a urinary biomarker and extended follow-up time. Environ Res 157, 44–51, doi:10.1016/j.envres.2017.05.011 (2017).

7 Min, J. Y. & Min, K. B. Blood cadmium levels and Alzheimer’s disease mortality risk in older US adults. Environ Health 15, 69, doi:10.1186/s12940-016-0155-7 (2016).

8 Okuda, B., Iwamoto, Y., Tachibana, H. & Sugita, M. Parkinsonism after acute cadmium poisoning. Clin Neurol Neurosurg 99, 263–265, doi:10.1016/s0303-8467(97)00090-5 (1997).

9 Voss, J. L., Bridge, D. J., Cohen, N. J. & Walker, J. A. A Closer Look at the Hippocampus and Memory. Trends Cogn Sci 21, 577–588, doi:10.1016/j.tics.2017.05.008 (2017).

10 Wang, H., Abel, G. M., Storm, D. R. & Xia, Z. Adolescent cadmium exposure impairs cognition and hippocampal neurogenesis in C57BL/6 mice. Environ Toxicol 37, 335–348, doi:10.1002/tox.23402 (2022).

11 Zhang, L., Wang, H., Abel, G. M., Storm, D. R. & Xia, Z. The Effects of Gene-Environment Interactions Between Cadmium Exposure and Apolipoprotein E4 on Memory in a Mouse Model of Alzheimer’s Disease. Toxicol Sci 173, 189–201, doi:10.1093/toxsci/kfz218 (2020).

12 Wang, H. et al. Inducible and Conditional Stimulation of Adult Hippocampal Neurogenesis Rescues Cadmium-Induced Impairments of Adult Hippocampal Neurogenesis and Hippocampus-Dependent Memory in Mice. Toxicol Sci 177, 263–280, doi:10.1093/toxsci/kfaa104 (2020).

13 Wang, H., Zhang, L., Abel, G. M., Storm, D. R. & Xia, Z. Cadmium Exposure Impairs Cognition and Olfactory Memory in Male C57BL/6 Mice. Toxicol Sci 161, 87–102, doi:10.1093/toxsci/kfx202 (2018).

14 Arruebarrena, M. A., Hawe, C. T., Lee, Y. M. & Branco, R. C. Mechanisms of Cadmium Neurotoxicity. Int J Mol Sci 24, doi:10.3390/ijms242316558 (2023).

15 Dempsey, J. L., Little, M. & Cui, J. Y. Gut microbiome: An intermediary to neurotoxicity. Neurotoxicology 75, 41–69, doi:10.1016/j.neuro.2019.08.005 (2019).

16 Zhuang, Z. Q. et al. Gut Microbiota is Altered in Patients with Alzheimer’s Disease. J Alzheimers Dis 63, 1337–1346, doi:10.3233/JAD-180176 (2018).

17 Vogt, N. M. et al. Gut microbiome alterations in Alzheimer’s disease. Sci Rep 7, 13537, doi:10.1038/s41598-017-13601-y (2017).

18 Zhang, L. et al. Altered Gut Microbiota in a Mouse Model of Alzheimer’s Disease. J Alzheimers Dis 60, 1241–1257, doi:10.3233/JAD-170020 (2017).

19 Harach, T. et al. Reduction of Abeta amyloid pathology in APPPS1 transgenic mice in the absence of gut microbiota. Sci Rep 7, 41802, doi:10.1038/srep41802 (2017).

20 Hazan, S. Rapid improvement in Alzheimer’s disease symptoms following fecal microbiota transplantation: a case report. J Int Med Res 48, 300060520925930, doi:10.1177/0300060520925930 (2020).

21 Park, S. H. et al. Cognitive function improvement after fecal microbiota transplantation in Alzheimer’s dementia patient: a case report. Curr Med Res Opin 37, 1739–1744, doi:10.1080/03007995.2021.1957807 (2021).

22 Savignac, H. M., Kiely, B., Dinan, T. G. & Cryan, J. F. Bifidobacteria exert strain-specific effects on stress-related behavior and physiology in BALB/c mice. Neurogastroenterol Motil 26, 1615–1627, doi:10.1111/nmo.12427 (2014).

23 Lew, L. C. et al. Probiotic Lactobacillus plantarum P8 alleviated stress and anxiety while enhancing memory and cognition in stressed adults: A randomised, double-blind, placebo-controlled study. Clin Nutr 38, 2053–2064, doi:10.1016/j.clnu.2018.09.010 (2019).

24 Allen, A. P. et al. Bifidobacterium longum 1714 as a translational psychobiotic: modulation of stress, electrophysiology and neurocognition in healthy volunteers. Transl Psychiatry 6, e939, doi:10.1038/tp.2016.191 (2016).

25 Smith, A. P., Sutherland, D. & Hewlett, P. An Investigation of the Acute Effects of Oligofructose-Enriched Inulin on Subjective Wellbeing, Mood and Cognitive Performance. Nutrients 7, 8887–8896, doi:10.3390/nu7115441 (2015).

26 Mayer, E. A., Nance, K. & Chen, S. The Gut-Brain Axis. Annu Rev Med 73, 439–453, doi:10.1146/annurev-med-042320-014032 (2022).

27 Dalile, B., Van Oudenhove, L., Vervliet, B. & Verbeke, K. The role of short-chain fatty acids in microbiota-gut-brain communication. Nat Rev Gastroenterol Hepatol 16, 461–478, doi:10.1038/s41575-019-0157-3 (2019).

28 MahmoudianDehkordi, S. et al. Altered bile acid profile associates with cognitive impairment in Alzheimer’s disease-An emerging role for gut microbiome. Alzheimers Dement 15, 76–92, doi:10.1016/j.jalz.2018.07.217 (2019).

29 Marksteiner, J., Blasko, I., Kemmler, G., Koal, T. & Humpel, C. Bile acid quantification of 20 plasma metabolites identifies lithocholic acid as a putative biomarker in Alzheimer’s disease. Metabolomics 14, 1, doi:10.1007/s11306-017-1297-5 (2018).

30 Pellegrini, C. et al. The intestinal barrier in disorders of the central nervous system. Lancet Gastroenterol Hepatol 8, 66–80, doi:10.1016/S2468-1253(22)00241-2 (2023).

31 Bercik, P., Collins, S. M. & Verdu, E. F. Microbes and the gut-brain axis. Neurogastroenterol Motil 24, 405–413, doi:10.1111/j.1365-2982.2012.01906.x (2012).

32 Rothhammer, V. et al. Type I interferons and microbial metabolites of tryptophan modulate astrocyte activity and central nervous system inflammation via the aryl hydrocarbon receptor. Nat Med 22, 586–597, doi:10.1038/nm.4106 (2016).

33 Forsyth, C. B. et al. Increased intestinal permeability correlates with sigmoid mucosa alpha-synuclein staining and endotoxin exposure markers in early Parkinson’s disease. PLoS One 6, e28032, doi:10.1371/journal.pone.0028032 (2011).

34 Esnafoglu, E. et al. Increased Serum Zonulin Levels as an Intestinal Permeability Marker in Autistic Subjects. J Pediatr 188, 240–244, doi:10.1016/j.jpeds.2017.04.004 (2017).

35 Solas, M., Milagro, F. I., Ramirez, M. J. & Martinez, J. A. Inflammation and gut-brain axis link obesity to cognitive dysfunction: plausible pharmacological interventions. Curr Opin Pharmacol 37, 87–92, doi:10.1016/j.coph.2017.10.005 (2017).

36 Stevens, B. R. et al. Increased human intestinal barrier permeability plasma biomarkers zonulin and FABP2 correlated with plasma LPS and altered gut microbiome in anxiety or depression. Gut 67, 1555–1557, doi:10.1136/gutjnl-2017-314759 (2018).

37 Wang, X., Liu, G. J., Gao, Q., Li, N. & Wang, R. T. C-type lectin-like receptor 2 and zonulin are associated with mild cognitive impairment and Alzheimer’s disease. Acta Neurol Scand 141, 250–255, doi:10.1111/ane.13196 (2020).

38 Tinkov, A. A. et al. Gut as a target for cadmium toxicity. Environmental Pollution 235, 429–434, doi:10.1016/j.envpol.2017.12.114 (2018).

39 Shao, M. & Zhu, Y. Long-term metal exposure changes gut microbiota of residents surrounding a mining and smelting area. Sci Rep 10, 4453, doi:10.1038/s41598-020-61143-7 (2020).

40 Zhai, Q. et al. Oral Administration of Probiotics Inhibits Absorption of the Heavy Metal Cadmium by Protecting the Intestinal Barrier. Appl Environ Microbiol 82, 4429–4440, doi:10.1128/AEM.00695-16 (2016).

41 Liu, Y., Li, Y., Liu, K. & Shen, J. Exposing to cadmium stress cause profound toxic effect on microbiota of the mice intestinal tract. PLoS One 9, e85323, doi:10.1371/journal.pone.0085323 (2014).

42 Liu, Y. et al. The Dysbiosis of Gut Microbiota Caused by Low-Dose Cadmium Aggravate the Injury of Mice Liver through Increasing Intestinal Permeability. Microorganisms 8, doi:10.3390/microorganisms8020211 (2020).

43 He, X. et al. Structural and functional alterations of gut microbiome in mice induced by chronic cadmium exposure. Chemosphere 246, 125747, doi:10.1016/j.chemosphere.2019.125747 (2020).

44 Engstrom, A. K., Snyder, J. M., Maeda, N. & Xia, Z. Gene-environment interaction between lead and Apolipoprotein E4 causes cognitive behavior deficits in mice. Mol Neurodegener 12, 14, doi:10.1186/s13024-017-0155-2 (2017).

45 Mallick, H. et al. Multivariable association discovery in population-scale meta-omics studies. PLoS Comput Biol 17, e1009442, doi:10.1371/journal.pcbi.1009442 (2021).

46 Caporaso, J. G. et al. QIIME allows analysis of high-throughput community sequencing data. Nat Methods 7, 335–336, doi:10.1038/nmeth.f.303 (2010).

47 Quinn, T. P. et al. A field guide for the compositional analysis of any-omics data. Gigascience 8, doi:10.1093/gigascience/giz107 (2019).

48 Gu, Z., Eils, R. & Schlesner, M. Complex heatmaps reveal patterns and correlations in multidimensional genomic data. Bioinformatics 32, 2847–2849, doi:10.1093/bioinformatics/btw313 (2016).

49 Gu, Z. & Hubschmann, D. Make Interactive Complex Heatmaps in R. Bioinformatics 38, 1460–1462, doi:10.1093/bioinformatics/btab806 (2022).

50 Kim, D., Langmead, B. & Salzberg, S. L. HISAT: a fast spliced aligner with low memory requirements. Nat Methods 12, 357–360, doi:10.1038/nmeth.3317 (2015).

51 Li, H. et al. The Sequence Alignment/Map format and SAMtools. Bioinformatics 25, 2078–2079, doi:10.1093/bioinformatics/btp352 (2009).

52 Liao, Y., Smyth, G. K. & Shi, W. The Subread aligner: fast, accurate and scalable read mapping by seed-and-vote. Nucleic Acids Res 41, e108, doi:10.1093/nar/gkt214 (2013).

53 Love, M. I., Huber, W. & Anders, S. Moderated estimation of fold change and dispersion for RNA-seq data with DESeq2. Genome Biol 15, 550, doi:10.1186/s13059-014-0550-8 (2014).

54 Alexa, A., Rahnenfuhrer, J. & Lengauer, T. Improved scoring of functional groups from gene expression data by decorrelating GO graph structure. Bioinformatics 22, 1600–1607, doi:10.1093/bioinformatics/btl140 (2006).

55 Gene Ontology, C., et al. The Gene Ontology knowledgebase in 2023. Genetics 224, doi:10.1093/genetics/iyad031 (2023).

56 Wickham, H. in ggplot2: Elegant Graphics for Data Analysis 241–253 (Springer International Publishing, 2016).

57 Dutta, M. et al. Chronic exposure to ambient traffic-related air pollution (TRAP) alters gut microbial abundance and bile acid metabolism in a transgenic rat model of Alzheimer’s disease. Toxicol Rep 9, 432–444, doi:10.1016/j.toxrep.2022.03.003 (2022).

58 Little, M. et al. Understanding the physiological functions of the host xenobiotic-sensing nuclear receptors PXR and CAR on the gut microbiome using genetically modified mice. Acta Pharm Sin B 12, 801–820, doi:10.1016/j.apsb.2021.07.022 (2022).

59 Wong, T. S. et al. G protein-coupled receptors in neurodegenerative diseases and psychiatric disorders. Signal Transduct Target Ther 8, 177, doi:10.1038/s41392-023-01427-2 (2023).

60 Bilkei-Gorzo, A., Racz, I., Michel, K. & Zimmer, A. Diminished anxiety- and depression-related behaviors in mice with selective deletion of the Tac1 gene. J Neurosci 22, 10046–10052, doi:10.1523/JNEUROSCI.22-22-10046.2002 (2002).

61 Zhang, M. et al. Ca-stimulated type 8 adenylyl cyclase is required for rapid acquisition of novel spatial information and for working/episodic-like memory. J Neurosci 28, 4736–4744, doi:10.1523/JNEUROSCI.1177-08.2008 (2008).

62 Weng, F. J. et al. Npas4 Is a Critical Regulator of Learning-Induced Plasticity at Mossy Fiber-CA3 Synapses during Contextual Memory Formation. Neuron 97, 1137–1152 e1135, doi:10.1016/j.neuron.2018.01.026 (2018).

63 Maya-Vetencourt, J. F. Activity-dependent NPAS4 expression and the regulation of gene programs underlying plasticity in the central nervous system. Neural Plast 2013, 683909, doi:10.1155/2013/683909 (2013).

64 Kaakoush, N. O. Insights into the Role of Erysipelotrichaceae in the Human Host. Front Cell Infect Microbiol 5, 84, doi:10.3389/fcimb.2015.00084 (2015).

65 de Theije, C. G. et al. Altered gut microbiota and activity in a murine model of autism spectrum disorders. Brain Behav Immun 37, 197–206, doi:10.1016/j.bbi.2013.12.005 (2014).

66 Haimov, I. et al. Variation in Gut Microbiota Composition is Associated with Sleep Quality and Cognitive Performance in Older Adults with Insomnia. Nat Sci Sleep 14, 1753–1767, doi:10.2147/NSS.S377114 (2022).

67 Ma, D. et al. Ketogenic diet enhances neurovascular function with altered gut microbiome in young healthy mice. Sci Rep 8, 6670, doi:10.1038/s41598-018-25190-5 (2018).

68 Magnusson, K. R. et al. Relationships between diet-related changes in the gut microbiome and cognitive flexibility. Neuroscience 300, 128–140, doi:10.1016/j.neuroscience.2015.05.016 (2015).

69 Meyer, K. et al. Association of the Gut Microbiota With Cognitive Function in Midlife. JAMA Netw Open 5, e2143941, doi:10.1001/jamanetworkopen.2021.43941 (2022).

70 Neth, B. J. et al. Modified ketogenic diet is associated with improved cerebrospinal fluid biomarker profile, cerebral perfusion, and cerebral ketone body uptake in older adults at risk for Alzheimer’s disease: a pilot study. Neurobiol Aging 86, 54–63, doi:10.1016/j.neurobiolaging.2019.09.015 (2020).

71 Zhao, Q. et al. Microbiota from healthy mice alleviates cognitive decline via reshaping the gut-brain metabolic axis in diabetic mice. Chem Biol Interact 382, 110638, doi:10.1016/j.cbi.2023.110638 (2023).

72 Zhou, X. et al. Longitudinal profiling of the microbiome at four body sites reveals core stability and individualized dynamics during health and disease. Cell Host Microbe 32, 506–526 e509, doi:10.1016/j.chom.2024.02.012 (2024).

73 Vadaq, N. et al. Untargeted Plasma Metabolomics and Gut Microbiome Profiling Provide Novel Insights into the Regulation of Platelet Reactivity in Healthy Individuals. Thromb Haemost 122, 529–539, doi:10.1055/a-1541-3706 (2022).

74 Pu, Y. et al. Gut microbial features and circulating metabolomic signatures of frailty in older adults. Nat Aging 4, 1249–1262, doi:10.1038/s43587-024-00678-0 (2024).

75 Lim, M. Y. et al. Gut Microbiome Structure and Association with Host Factors in a Korean Population. mSystems 6, e0017921, doi:10.1128/mSystems.00179-21 (2021).

76 Asnicar, F. et al. Microbiome connections with host metabolism and habitual diet from 1,098 deeply phenotyped individuals. Nat Med 27, 321–332, doi:10.1038/s41591-020-01183-8 (2021).

77 Liu, P. et al. The role of short-chain fatty acids in intestinal barrier function, inflammation, oxidative stress, and colonic carcinogenesis. Pharmacol Res 165, 105420, doi:10.1016/j.phrs.2021.105420 (2021).

78 Cryan, J. F. et al. The Microbiota-Gut-Brain Axis. Physiol Rev 99, 1877–2013, doi:10.1152/physrev.00018.2018 (2019).

79 Stolfi, C., Maresca, C., Monteleone, G. & Laudisi, F. Implication of Intestinal Barrier Dysfunction in Gut Dysbiosis and Diseases. Biomedicines 10, doi:10.3390/biomedicines10020289 (2022).

80 Kinashi, Y. & Hase, K. Partners in Leaky Gut Syndrome: Intestinal Dysbiosis and Autoimmunity. Front Immunol 12, 673708, doi:10.3389/fimmu.2021.673708 (2021).

81 Sivaprakasam, S., Bhutia, Y. D., Yang, S. & Ganapathy, V. Short-Chain Fatty Acid Transporters: Role in Colonic Homeostasis. Compr Physiol 8, 299–314, doi:10.1002/cphy.c170014 (2017).

82 Goncalves, P., Gregorio, I. & Martel, F. The short-chain fatty acid butyrate is a substrate of breast cancer resistance protein. Am J Physiol Cell Physiol 301, C984–994, doi:10.1152/ajpcell.00146.2011 (2011).

83 Goncalves, P. & Martel, F. Butyrate and colorectal cancer: the role of butyrate transport. Curr Drug Metab 14, 994–1008, doi:10.2174/1389200211314090006 (2013).

84 Englund, G. et al. Efflux transporters in ulcerative colitis: decreased expression of BCRP (ABCG2) and Pgp (ABCB1). Inflamm Bowel Dis 13, 291–297, doi:10.1002/ibd.20030 (2007).

85 Takiishi, T., Fenero, C. I. M. & Camara, N. O. S. Intestinal barrier and gut microbiota: Shaping our immune responses throughout life. Tissue Barriers 5, e1373208, doi:10.1080/21688370.2017.1373208 (2017).

86 Wang, L. Y., Tu, Y. F., Lin, Y. C. & Huang, C. C. CXCL5 signaling is a shared pathway of neuroinflammation and blood-brain barrier injury contributing to white matter injury in the immature brain. J Neuroinflammation 13, 6, doi:10.1186/s12974-015-0474-6 (2016).

87 Amorim, A. et al. IFNgamma and GM-CSF control complementary differentiation programs in the monocyte-to-phagocyte transition during neuroinflammation. Nat Immunol 23, 217–228, doi:10.1038/s41590-021-01117-7 (2022).

88 Shaftel, S. S., Griffin, W. S. & O’Banion, M. K. The role of interleukin-1 in neuroinflammation and Alzheimer disease: an evolving perspective. J Neuroinflammation 5, 7, doi:10.1186/1742-2094-5-7 (2008).

89 Kearns, R. Gut-Brain Axis and Neuroinflammation: The Role of Gut Permeability and the Kynurenine Pathway in Neurological Disorders. Cell Mol Neurobiol 44, 64, doi:10.1007/s10571-024-01496-z (2024).

90 Boraschi, D., Italiani, P., Weil, S. & Martin, M. U. The family of the interleukin-1 receptors. Immunol Rev 281, 197–232, doi:10.1111/imr.12606 (2018).

91 Dinarello, C. A. Immunological and inflammatory functions of the interleukin-1 family. Annu Rev Immunol 27, 519–550, doi:10.1146/annurev.immunol.021908.132612 (2009).

92 Cruikshank, W. W., Kornfeld, H. & Center, D. M. Interleukin-16. J Leukoc Biol 67, 757–766, doi:10.1002/jlb.67.6.757 (2000).

93 Huang, Y. et al. IL-16 regulates macrophage polarization as a target gene of mir-145-3p. Mol Immunol 107, 1–9, doi:10.1016/j.molimm.2018.12.027 (2019).

94 Hridi, S. U. et al. Increased Levels of IL-16 in the Central Nervous System during Neuroinflammation Are Associated with Infiltrating Immune Cells and Resident Glial Cells. Biology (Basel) 10, doi:10.3390/biology10060472 (2021).

95 Schumann, R. R. Old and new findings on lipopolysaccharide-binding protein: a soluble pattern-recognition molecule. Biochem Soc Trans 39, 989–993, doi:10.1042/BST0390989 (2011).

96 Zhao, Y. et al. Lipopolysaccharide-binding protein and future Parkinson’s disease risk: a European prospective cohort. J Neuroinflammation 20, 170, doi:10.1186/s12974-023-02846-2 (2023).

97 Romo, E. Z. et al. Elevated lipopolysaccharide binding protein in Alzheimer’s disease patients with APOE3/E3 but not APOE3/E4 genotype. Front Neurol 15, 1408220, doi:10.3389/fneur.2024.1408220 (2024).

98 Escribano, B. M. et al. Lipopolysaccharide Binding Protein and Oxidative Stress in a Multiple Sclerosis Model. Neurotherapeutics 14, 199–211, doi:10.1007/s13311-016-0480-0 (2017).

99 Wu, T. et al. Complement C3 Is Activated in Human AD Brain and Is Required for Neurodegeneration in Mouse Models of Amyloidosis and Tauopathy. Cell Rep 28, 2111–2123 e2116, doi:10.1016/j.celrep.2019.07.060 (2019).

100 Shi, Q. et al. Complement C3 deficiency protects against neurodegeneration in aged plaque-rich APP/PS1 mice. Sci Transl Med 9, doi:10.1126/scitranslmed.aaf6295 (2017).

101 Hong, S. et al. Complement and microglia mediate early synapse loss in Alzheimer mouse models. Science 352, 712–716, doi:10.1126/science.aad8373 (2016).

102 Brombacher, T. M. et al. IL-13-Mediated Regulation of Learning and Memory. J Immunol 198, 2681–2688, doi:10.4049/jimmunol.1601546 (2017).

103 Hanuscheck, N. et al. Interleukin-4 receptor signaling modulates neuronal network activity. J Exp Med 219, doi:10.1084/jem.20211887 (2022).

104 Herz, J. et al. GABAergic neuronal IL-4R mediates T cell effect on memory. Neuron 109, 3609–3618 e3609, doi:10.1016/j.neuron.2021.10.022 (2021).

105 Vogelaar, C. F. et al. Fast direct neuronal signaling via the IL-4 receptor as therapeutic target in neuroinflammation. Sci Transl Med 10, doi:10.1126/scitranslmed.aao2304 (2018).

106 Yan, P. et al. The establishment of humanized IL-4/IL-4RA mouse model by gene editing and efficacy evaluation. Immunobiology 225, 151998, doi:10.1016/j.imbio.2020.151998 (2020).

107 Wang, X. et al. DUSP1 Promotes Microglial Polarization toward M2 Phenotype in the Medial Prefrontal Cortex of Neuropathic Pain Rats via Inhibition of MAPK Pathway. ACS Chem Neurosci 12, 966–978, doi:10.1021/acschemneuro.0c00567 (2021).

108 Baker, B. J., Akhtar, L. N. & Benveniste, E. N. SOCS1 and SOCS3 in the control of CNS immunity. Trends Immunol 30, 392–400, doi:10.1016/j.it.2009.07.001 (2009).

109 Song, H. et al. Tauroursodeoxycholic acid: a bile acid that may be used for the prevention and treatment of Alzheimer’s disease. Front Neurosci 18, 1348844, doi:10.3389/fnins.2024.1348844 (2024).

110 Holzer, P. & Farzi, A. Neuropeptides and the microbiota-gut-brain axis. Adv Exp Med Biol 817, 195–219, doi:10.1007/978-1-4939-0897-4_9 (2014).

111 Pan, Y. W., Storm, D. R. & Xia, Z. Role of adult neurogenesis in hippocampus-dependent memory, contextual fear extinction and remote contextual memory: new insights from ERK5 MAP kinase. Neurobiol Learn Mem 105, 81–92, doi:10.1016/j.nlm.2013.07.011 (2013).

112 Wang, W. B. et al. Genetic Activation of ERK5 MAP Kinase Enhances Adult Neurogenesis and Extends Hippocampus-Dependent Long-Term Memory. Journal of Neuroscience 34, 2130–2147 (2014).

113 Matsushita, M. T., Wang, H., Abel, G. M. & Xia, Z. Inducible and Conditional Activation of Adult Neurogenesis Rescues Cadmium-Induced Hippocampus-Dependent Memory Deficits in ApoE4-KI Mice. Int J Mol Sci 24, doi:10.3390/ijms24119118 (2023).

114 Jong, Y. I., Harmon, S. K. & O’Malley, K. L. Intracellular GPCRs Play Key Roles in Synaptic Plasticity. ACS Chem Neurosci 9, 2162–2172, doi:10.1021/acschemneuro.7b00516 (2018).

115 da Silva, W. C., Kohler, C. C., Radiske, A. & Cammarota, M. D1/D5 dopamine receptors modulate spatial memory formation. Neurobiol Learn Mem 97, 271–275, doi:10.1016/j.nlm.2012.01.005 (2012).

116 Schultz, W. Predictive reward signal of dopamine neurons. J Neurophysiol 80, 1–27, doi:10.1152/jn.1998.80.1.1 (1998).

117 Zhang, A., Neumeyer, J. L. & Baldessarini, R. J. Recent progress in development of dopamine receptor subtype-selective agents: potential therapeutics for neurological and psychiatric disorders. Chem Rev 107, 274–302, doi:10.1021/cr050263h (2007).

118 Civelli, O., Bunzow, J. R. & Grandy, D. K. Molecular diversity of the dopamine receptors. Annu Rev Pharmacol Toxicol 33, 281–307, doi:10.1146/annurev.pa.33.040193.001433 (1993).

119 Hotte, M. et al. D1 receptor modulation of memory retrieval performance is associated with changes in pCREB and pDARPP-32 in rat prefrontal cortex. Behav Brain Res 171, 127–133, doi:10.1016/j.bbr.2006.03.026 (2006).

120 Hotte, M., Naudon, L. & Jay, T. M. Modulation of recognition and temporal order memory retrieval by dopamine D1 receptor in rats. Neurobiol Learn Mem 84, 85–92, doi:10.1016/j.nlm.2005.04.002 (2005).

121 de Lima, M. N. et al. Modulatory influence of dopamine receptors on consolidation of object recognition memory. Neurobiol Learn Mem 95, 305–310, doi:10.1016/j.nlm.2010.12.007 (2011).

122 Rocchetti, J. et al. Presynaptic D2 dopamine receptors control long-term depression expression and memory processes in the temporal hippocampus. Biol Psychiatry 77, 513–525, doi:10.1016/j.biopsych.2014.03.013 (2015).

123 Dalley, J. W. & Everitt, B. J. Dopamine receptors in the learning, memory and drug reward circuitry. Semin Cell Dev Biol 20, 403–410, doi:10.1016/j.semcdb.2009.01.002 (2009).

124 Dere, E. et al. Neuronal histamine and the interplay of memory, reinforcement and emotions. Behav Brain Res 215, 209–220, doi:10.1016/j.bbr.2009.12.045 (2010).

125 Dai, H. et al. Selective cognitive dysfunction in mice lacking histamine H1 and H2 receptors. Neurosci Res 57, 306–313, doi:10.1016/j.neures.2006.10.020 (2007).

126 Ambree, O. et al. Impaired spatial learning and reduced adult hippocampal neurogenesis in histamine H1-receptor knockout mice. Eur Neuropsychopharmacol 24, 1394–1404, doi:10.1016/j.euroneuro.2014.04.006 (2014).

127 Zlomuzica, A. et al. Neuronal histamine and cognitive symptoms in Alzheimer’s disease. Neuropharmacology 106, 135–145, doi:10.1016/j.neuropharm.2015.05.007 (2016).

128 Borbely, E., Scheich, B. & Helyes, Z. Neuropeptides in learning and memory. Neuropeptides 47, 439–450, doi:10.1016/j.npep.2013.10.012 (2013).

129 Schlesinger, K. et al. Substance P enhancement of passive and active avoidance conditioning in mice. Pharmacol Biochem Behav 19, 655–661, doi:10.1016/0091-3057(83)90341-6 (1983).

130 Schlesinger, K. et al. Substance P facilitation of memory: effects in an appetitively motivated learning task. Behav Neural Biol 45, 230–239, doi:10.1016/s0163-1047(86)90805-8 (1986).

131 Li, W. X., et al. Integrated Analysis of Alzheimer’s Disease and Schizophrenia Dataset Revealed Different Expression Pattern in Learning and Memory. J Alzheimers Dis 51, 417–425, doi:10.3233/JAD-150807 (2016).

132 Liu, Y. J. et al. Identification of hub genes associated with cognition in the hippocampus of Alzheimer’s Disease. Bioengineered 12, 9598–9609, doi:10.1080/21655979.2021.1999549 (2021).

133 Zhang, M. & Wang, H. Ca(2+)-stimulated ADCY1 and ADCY8 regulate distinct aspects of synaptic and cognitive flexibility. Front Cell Neurosci 17, 1215255, doi:10.3389/fncel.2023.1215255 (2023).

134 Ramamoorthi, K. et al. Npas4 regulates a transcriptional program in CA3 required for contextual memory formation. Science 334, 1669–1675, doi:10.1126/science.1208049 (2011).

135 Qiu, J. et al. Decreased Npas4 and Arc mRNA Levels in the Hippocampus of Aged Memory-Impaired Wild-Type But Not Memory Preserved 11beta-HSD1 Deficient Mice. J Neuroendocrinol 28, n/a, doi:10.1111/jne.12339 (2016).

136 Kumar, A. Editorial: Neuroinflammation and Cognition. Front Aging Neurosci 10, 413, doi:10.3389/fnagi.2018.00413 (2018).

137 Knezevic, D. & Mizrahi, R. Molecular imaging of neuroinflammation in Alzheimer’s disease and mild cognitive impairment. Prog Neuropsychopharmacol Biol Psychiatry 80, 123–131, doi:10.1016/j.pnpbp.2017.05.007 (2018).

138 Barrientos, R. M., Kitt, M. M., Watkins, L. R. & Maier, S. F. Neuroinflammation in the normal aging hippocampus. Neuroscience 309, 84–99, doi:10.1016/j.neuroscience.2015.03.007 (2015).

139 Di Benedetto, S., Muller, L., Wenger, E., Duzel, S. & Pawelec, G. Contribution of neuroinflammation and immunity to brain aging and the mitigating effects of physical and cognitive interventions. Neurosci Biobehav Rev 75, 114–128, doi:10.1016/j.neubiorev.2017.01.044 (2017).

140 Ellwardt, E. & Zipp, F. Molecular mechanisms linking neuroinflammation and neurodegeneration in MS. Exp Neurol 262 Pt A, 8–17, doi:10.1016/j.expneurol.2014.02.006 (2014).

141 De Virgilio, A. et al. Parkinson’s disease: Autoimmunity and neuroinflammation. Autoimmun Rev 15, 1005–1011, doi:10.1016/j.autrev.2016.07.022 (2016).

142 Heneka, M. T. et al. Neuroinflammation in Alzheimer’s disease. Lancet Neurol 14, 388–405, doi:10.1016/S1474-4422(15)70016-5 (2015).

143 Di Rosa, M. et al. Chitotriosidase and inflammatory mediator levels in Alzheimer’s disease and cerebrovascular dementia. Eur J Neurosci 23, 2648–2656, doi:10.1111/j.1460-9568.2006.04780.x (2006).

144 Anvar, N. E. et al. Association between polymorphisms in Interleukin-16 gene and risk of late-onset Alzheimer’s disease. J Neurol Sci 358, 324–327, doi:10.1016/j.jns.2015.09.344 (2015).

145 Andre, P. et al. Lipopolysaccharide-Binding Protein, Soluble CD14, and the Long-Term Risk of Alzheimer’s Disease: A Nested Case-Control Pilot Study of Older Community Dwellers from the Three-City Cohort. J Alzheimers Dis 71, 751–761, doi:10.3233/JAD-190295 (2019).

146 Liddelow, S. A. et al. Neurotoxic reactive astrocytes are induced by activated microglia. Nature 541, 481–487, doi:10.1038/nature21029 (2017).

147 Tassoni, A. et al. The astrocyte transcriptome in EAE optic neuritis shows complement activation and reveals a sex difference in astrocytic C3 expression. Sci Rep 9, 10010, doi:10.1038/s41598-019-46232-6 (2019).

148 Cao, Q. et al. Astrocytic CXCL5 hinders microglial phagocytosis of myelin debris and aggravates white matter injury in chronic cerebral ischemia. J Neuroinflammation 20, 105, doi:10.1186/s12974-023-02780-3 (2023).

149 Cribbs, D. H. et al. Extensive innate immune gene activation accompanies brain aging, increasing vulnerability to cognitive decline and neurodegeneration: a microarray study. J Neuroinflammation 9, 179, doi:10.1186/1742-2094-9-179 (2012).

150 Hendriks, W. J. & Pulido, R. Protein tyrosine phosphatase variants in human hereditary disorders and disease susceptibilities. Biochim Biophys Acta 1832, 1673–1696, doi:10.1016/j.bbadis.2013.05.022 (2013).

151 Salojin, K. & Oravecz, T. Regulation of innate immunity by MAPK dual-specificity phosphatases: knockout models reveal new tricks of old genes. J Leukoc Biol 81, 860–869, doi:10.1189/jlb.1006639 (2007).

152 Wancket, L. M., Frazier, W. J. & Liu, Y. Mitogen-activated protein kinase phosphatase (MKP)-1 in immunology, physiology, and disease. Life Sci 90, 237–248, doi:10.1016/j.lfs.2011.11.017 (2012).

153 Shao, L. L., Gao, M. M., Gong, J. X. & Yang, L. Y. DUSP1 regulates hippocampal damage in epilepsy rats via ERK1/2 pathway. J Chem Neuroanat 118, 102032, doi:10.1016/j.jchemneu.2021.102032 (2021).

154 Cianciulli, A., Calvello, R., Porro, C., Trotta, T. & Panaro, M. A. Understanding the role of SOCS signaling in neurodegenerative diseases: Current and emerging concepts. Cytokine Growth Factor Rev 37, 67–79, doi:10.1016/j.cytogfr.2017.07.005 (2017).

155 Pedros, I. et al. Adipokine pathways are altered in hippocampus of an experimental mouse model of Alzheimer’s disease. J Nutr Health Aging 19, 403–412, doi:10.1007/s12603-014-0574-5 (2015).

156 Li, Y., Chu, N., Rostami, A. & Zhang, G. X. Dendritic cells transduced with SOCS-3 exhibit a tolerogenic/DC2 phenotype that directs type 2 Th cell differentiation in vitro and in vivo. J Immunol 177, 1679–1688, doi:10.4049/jimmunol.177.3.1679 (2006).

157 Tang, W. et al. Roles of Gut Microbiota in the Regulation of Hippocampal Plasticity, Inflammation, and Hippocampus-Dependent Behaviors. Front Cell Infect Microbiol 10, 611014, doi:10.3389/fcimb.2020.611014 (2020).

158 Kuijer, E. J. & Steenbergen, L. The microbiota-gut-brain axis in hippocampus-dependent learning and memory: current state and future challenges. Neurosci Biobehav Rev 152, 105296, doi:10.1016/j.neubiorev.2023.105296 (2023).

159 Serra, M. C., Nocera, J. R., Kelleher, J. L. & Addison, O. Prebiotic Intake in Older Adults: Effects on Brain Function and Behavior. Curr Nutr Rep 8, 66–73, doi:10.1007/s13668-019-0265-2 (2019).

160 Caliskan, G. et al. Antibiotic-induced gut dysbiosis leads to activation of microglia and impairment of cholinergic gamma oscillations in the hippocampus. Brain Behav Immun 99, 203–217, doi:10.1016/j.bbi.2021.10.007 (2022).

161 Zhang, T. et al. Longitudinal and Multi-Kingdom Gut Microbiome Alterations in a Mouse Model of Alzheimer’s Disease. Int J Mol Sci 25, doi:10.3390/ijms252111472 (2024).

162 Verhaar, B. J. H. et al. Gut Microbiota Composition Is Related to AD Pathology. Front Immunol 12, 794519, doi:10.3389/fimmu.2021.794519 (2021).

163 Shen, H. et al. New mechanism of neuroinflammation in Alzheimer’s disease: The activation of NLRP3 inflammasome mediated by gut microbiota. Prog Neuropsychopharmacol Biol Psychiatry 100, 109884, doi:10.1016/j.pnpbp.2020.109884 (2020).

164 Zhai, Q. et al. Effects of subchronic oral toxic metal exposure on the intestinal microbiota of mice. Sci Bull (Beijing) 62, 831–840, doi:10.1016/j.scib.2017.01.031 (2017).

165 Zhang, J. et al. Subchronic cadmium exposure upregulates the mRNA level of genes associated to hepatic lipid metabolism in adult female CD1 mice. J Appl Toxicol 38, 1026–1035, doi:10.1002/jat.3612 (2018).

166 Li, X. et al. Heavy metal exposure causes changes in the metabolic health-associated gut microbiome and metabolites. Environ Int 126, 454–467, doi:10.1016/j.envint.2019.02.048 (2019).

167 Yang, J., Chen, W., Sun, Y., Liu, J. & Zhang, W. Effects of cadmium on organ function, gut microbiota and its metabolomics profile in adolescent rats. Ecotoxicol Environ Saf 222, 112501, doi:10.1016/j.ecoenv.2021.112501 (2021).

168 Fromentin, S. et al. Microbiome and metabolome features of the cardiometabolic disease spectrum. Nat Med 28, 303–314, doi:10.1038/s41591-022-01688-4 (2022).

169 Guo, M. et al. Guild-Level Microbiome Signature Associated with COVID-19 Severity and Prognosis. mBio 14, e0351922, doi:10.1128/mbio.03519-22 (2023).

170 Zhou, W. et al. Skin microbiome attributes associate with biophysical skin ageing. Exp Dermatol 32, 1546–1556, doi:10.1111/exd.14863 (2023).

171 Waluga, M. in A Comprehensive Overview of Irritable Bowel Syndrome (ed Jakub Fichna) 107–127 (Academic Press, 2020).

172 Chen, Y. et al. Gut microbiota-driven metabolic alterations reveal gut-brain communication in Alzheimer’s disease model mice. Gut Microbes 16, 2302310, doi:10.1080/19490976.2024.2302310 (2024).

173 Peng, W. et al. Association of gut microbiota composition and function with a senescence-accelerated mouse model of Alzheimer’s Disease using 16S rRNA gene and metagenomic sequencing analysis. Aging (Albany NY) 10, 4054–4065, doi:10.18632/aging.101693 (2018).

174 Yang, X. et al. Alterations in gut microbiota contribute to cognitive deficits induced by chronic infection of Toxoplasma gondii. Brain Behav Immun 119, 394–407, doi:10.1016/j.bbi.2024.04.008 (2024).

175 Jiao, Y. et al. Salidroside ameliorates memory impairment following long-term ethanol intake in rats by modulating the altered intestinal microbiota content and hippocampal gene expression. Front Microbiol 14, 1172936, doi:10.3389/fmicb.2023.1172936 (2023).

176 Ren, Z. L. et al. Causal relationships between gut microbiota and dementia: A two-sample, bidirectional, Mendelian randomization study. World J Clin Cases 12, 2780–2788, doi:10.12998/wjcc.v12.i16.2780 (2024).

177 Fu, J., Qin, Y., Xiao, L. & Dai, X. Causal relationship between gut microflora and dementia: a Mendelian randomization study. Front Microbiol 14, 1306048, doi:10.3389/fmicb.2023.1306048 (2023).

178 Shi, L., Liu, X., Zhang, S. & Zhou, A. Association of gut microbiota with cerebral cortical thickness: A Mendelian randomization study. J Affect Disord 352, 312–320, doi:10.1016/j.jad.2024.02.063 (2024).

179 Haran, J. P. et al. Alzheimer’s Disease Microbiome Is Associated with Dysregulation of the Anti-Inflammatory P-Glycoprotein Pathway. mBio 10, 10.1128/mbio.00632-00619, doi:doi:10.1128/mbio.00632-19 (2019).

180 O’Riordan, K. J. et al. Short chain fatty acids: Microbial metabolites for gut-brain axis signalling. Mol Cell Endocrinol 546, 111572, doi:10.1016/j.mce.2022.111572 (2022).

181 Ueda, A. et al. Identification of Faecalibacterium prausnitzii strains for gut microbiome-based intervention in Alzheimer’s-type dementia. Cell Rep Med 2, 100398, doi:10.1016/j.xcrm.2021.100398 (2021).

182 Pei, Y., Lu, Y., Li, H., Jiang, C. & Wang, L. Gut microbiota and intestinal barrier function in subjects with cognitive impairments: a cross-sectional study. Front Aging Neurosci 15, 1174599, doi:10.3389/fnagi.2023.1174599 (2023).

183 Fan, K. C. et al. Altered gut microbiota in older adults with mild cognitive impairment: a case-control study. Front Aging Neurosci 15, 1162057, doi:10.3389/fnagi.2023.1162057 (2023).

184 Carlier, J. P., Bedora-Faure, M., K’Ouas, G., Alauzet, C. & Mory, F. Proposal to unify Clostridium orbiscindens Winter et al. 1991 and Eubacterium plautii (Seguin 1928) Hofstad and Aasjord 1982, with description of Flavonifractor plautii gen. nov., comb. nov., and reassignment of Bacteroides capillosus to Pseudoflavonifractor capillosus gen. nov., comb. nov. Int J Syst Evol Microbiol 60, 585–590, doi:10.1099/ijs.0.016725-0 (2010).

185 Boots, A. W., Haenen, G. R. & Bast, A. Health effects of quercetin: from antioxidant to nutraceutical. Eur J Pharmacol 585, 325–337, doi:10.1016/j.ejphar.2008.03.008 (2008).

186 Sittipo, P., Choi, J., Lee, S. & Lee, Y. K. The function of gut microbiota in immune-related neurological disorders: a review. J Neuroinflammation 19, 154, doi:10.1186/s12974-022-02510-1 (2022).

187 Coello, K. et al. Gut microbiota composition in patients with newly diagnosed bipolar disorder and their unaffected first-degree relatives. Brain Behav Immun 75, 112–118, doi:10.1016/j.bbi.2018.09.026 (2019).

188 Coello, K. et al. Affective disorders impact prevalence of Flavonifractor and abundance of Christensenellaceae in gut microbiota. Prog Neuropsychopharmacol Biol Psychiatry 110, 110300, doi:10.1016/j.pnpbp.2021.110300 (2021).

189 Rizzatti, G., Lopetuso, L. R., Gibiino, G., Binda, C. & Gasbarrini, A. Proteobacteria: A Common Factor in Human Diseases. Biomed Res Int 2017, 9351507, doi:10.1155/2017/9351507 (2017).

190 Liu, P. et al. Altered microbiomes distinguish Alzheimer’s disease from amnestic mild cognitive impairment and health in a Chinese cohort. Brain Behav Immun 80, 633–643, doi:10.1016/j.bbi.2019.05.008 (2019).

191 Nagpal, R., Neth, B. J., Wang, S., Craft, S. & Yadav, H. Modified Mediterranean-ketogenic diet modulates gut microbiome and short-chain fatty acids in association with Alzheimer’s disease markers in subjects with mild cognitive impairment. EBioMedicine 47, 529–542, doi:10.1016/j.ebiom.2019.08.032 (2019).

192 Wu, F. et al. Gut Microbiota and Subjective Memory Complaints in Older Women. J Alzheimers Dis 88, 251–262, doi:10.3233/JAD-220011 (2022).

193 Bauerl, C., Collado, M. C., Diaz Cuevas, A., Vina, J. & Perez Martinez, G. Shifts in gut microbiota composition in an APP/PSS1 transgenic mouse model of Alzheimer’s disease during lifespan. Lett Appl Microbiol 66, 464–471, doi:10.1111/lam.12882 (2018).

194 Wei, Z., Li, D. & Shi, J. Alterations of Spatial Memory and Gut Microbiota Composition in Alzheimer’s Disease Triple-Transgenic Mice at 3, 6, and 9 Months of Age. Am J Alzheimers Dis Other Demen 38, 15333175231174193, doi:10.1177/15333175231174193 (2023).

195 Goldman, E. & Green, L. H. Practical handbook of microbiology. Fourth edition. edn, (CRC Press, 2021).

196 Chen, G., Zhou, X., Zhu, Y., Shi, W. & Kong, L. Gut microbiome characteristics in subjective cognitive decline, mild cognitive impairment and Alzheimer’s disease: a systematic review and meta-analysis. Eur J Neurol 30, 3568–3580, doi:10.1111/ene.15961 (2023).

197 Escobar, Y. H., O’Piela, D., Wold, L. E. & Mackos, A. R. Influence of the Microbiota-Gut-Brain Axis on Cognition in Alzheimer’s Disease. J Alzheimers Dis 87, 17–31, doi:10.3233/JAD-215290 (2022).

198 Lu, S. et al. Gut Microbiota and Targeted Biomarkers Analysis in Patients With Cognitive Impairment. Front Neurol 13, 834403, doi:10.3389/fneur.2022.834403 (2022).

199 Qu, L. et al. Gut Microbiome Signatures Are Predictive of Cognitive Impairment in Hypertension Patients-A Cohort Study. Front Microbiol 13, 841614, doi:10.3389/fmicb.2022.841614 (2022).

200 Cattaneo, A. et al. Association of brain amyloidosis with pro-inflammatory gut bacterial taxa and peripheral inflammation markers in cognitively impaired elderly. Neurobiol Aging 49, 60–68, doi:10.1016/j.neurobiolaging.2016.08.019 (2017).

201 Hennessey, C. et al. Neonatal Enteropathogenic Escherichia coli Infection Disrupts Microbiota-Gut-Brain Axis Signaling. Infect Immun 89, e0005921, doi:10.1128/IAI.00059-21 (2021).

202 Bland, S. T. et al. Enduring consequences of early-life infection on glial and neural cell genesis within cognitive regions of the brain. Brain Behav Immun 24, 329–338, doi:10.1016/j.bbi.2009.09.012 (2010).

203 Schutze, S. et al. Intracerebral Infection with E. coli Impairs Spatial Learning and Induces Necrosis of Hippocampal Neurons in the Tg2576 Mouse Model of Alzheimer’s Disease. J Alzheimers Dis Rep 6, 101–114, doi:10.3233/ADR-210049 (2022).

204 Yang, R. C. et al. Meningitic Escherichia coli-induced upregulation of PDGF-B and ICAM-1 aggravates blood-brain barrier disruption and neuroinflammatory response. J Neuroinflammation 16, 101, doi:10.1186/s12974-019-1497-1 (2019).

205 Fang, P., Kazmi, S. A., Jameson, K. G. & Hsiao, E. Y. The Microbiome as a Modifier of Neurodegenerative Disease Risk. Cell Host Microbe 28, 201–222, doi:10.1016/j.chom.2020.06.008 (2020).

206 Li, Z., Zhu, H., Zhang, L. & Qin, C. The intestinal microbiome and Alzheimer’s disease: A review. Animal Model Exp Med 1, 180–188, doi:10.1002/ame2.12033 (2018).

207 Xu, R. et al. The role of the probiotic Akkermansia muciniphila in brain functions: insights underpinning therapeutic potential. Crit Rev Microbiol 49, 151–176, doi:10.1080/1040841X.2022.2044286 (2023).

208 Cao, W. et al. Causal relationship of gut microbiota and metabolites on cognitive performance: A mendelian randomization analysis. Neurobiol Dis 191, 106395, doi:10.1016/j.nbd.2023.106395 (2024).

209 Li, J. et al. Clostridiales are predominant microbes that mediate psychiatric disorders. J Psychiatr Res 130, 48–56, doi:10.1016/j.jpsychires.2020.07.018 (2020).

210 Zheng, M. et al. Probiotic Clostridium butyricum ameliorates cognitive impairment in obesity via the microbiota-gut-brain axis. Brain Behav Immun 115, 565–587, doi:10.1016/j.bbi.2023.11.016 (2024).

211 De Angelis, M. et al. Fecal microbiota and metabolome of children with autism and pervasive developmental disorder not otherwise specified. PLoS One 8, e76993, doi:10.1371/journal.pone.0076993 (2013).

212 Shiosaka, S. Kallikrein 8: A key sheddase to strengthen and stabilize neural plasticity. Neurosci Biobehav Rev 140, 104774, doi:10.1016/j.neubiorev.2022.104774 (2022).

213 Tamura, H. et al. Neuropsin is essential for early processes of memory acquisition and Schaffer collateral long-term potentiation in adult mouse hippocampus in vivo. J Physiol 570, 541–551, doi:10.1113/jphysiol.2005.098715 (2006).

214 Herring, A. et al. Kallikrein-8 inhibition attenuates Alzheimer’s disease pathology in mice. Alzheimers Dement 12, 1273–1287, doi:10.1016/j.jalz.2016.05.006 (2016).

215 Shimizu-Okabe, C. et al. Expression of the kallikrein gene family in normal and Alzheimer’s disease brain. Neuroreport 12, 2747–2751, doi:10.1097/00001756-200108280-00031 (2001).

216 Pensold, D. et al. The DNA Methyltransferase 1 (DNMT1) Controls the Shape and Dynamics of Migrating POA-Derived Interneurons Fated for the Murine Cerebral Cortex. Cereb Cortex 27, 5696–5714, doi:10.1093/cercor/bhw341 (2017).

217 Civiero, L. et al. Leucine-rich repeat kinase 2 interacts with p21-activated kinase 6 to control neurite complexity in mammalian brain. J Neurochem 135, 1242–1256, doi:10.1111/jnc.13369 (2015).

218 Nekrasova, T., Jobes, M. L., Ting, J. H., Wagner, G. C. & Minden, A. Targeted disruption of the Pak5 and Pak6 genes in mice leads to deficits in learning and locomotion. Dev Biol 322, 95–108, doi:10.1016/j.ydbio.2008.07.006 (2008).

219 Herring, B. E. & Nicoll, R. A. Long-Term Potentiation: From CaMKII to AMPA Receptor Trafficking. Annu Rev Physiol 78, 351–365, doi:10.1146/annurev-physiol-021014-071753 (2016).

220 Lisman, J., Yasuda, R. & Raghavachari, S. Mechanisms of CaMKII action in long-term potentiation. Nat Rev Neurosci 13, 169–182, doi:10.1038/nrn3192 (2012).

221 Astudillo, D. et al. CaMKII inhibitor 1 (CaMK2N1) mRNA is upregulated following LTP induction in hippocampal slices. Synapse 74, e22158, doi:10.1002/syn.22158 (2020).

222 Iijima, T. et al. Hzf protein regulates dendritic localization and BDNF-induced translation of type 1 inositol 1,4,5-trisphosphate receptor mRNA. Proc Natl Acad Sci U S A 102, 17190–17195, doi:10.1073/pnas.0504684102 (2005).

223 Yu, L. et al. Association Between Brain Gene Expression, DNA Methylation, and Alteration of Ex Vivo Magnetic Resonance Imaging Transverse Relaxation in Late-Life Cognitive Decline. JAMA Neurol 74, 1473–1480, doi:10.1001/jamaneurol.2017.2807 (2017).

224 Senn, C. et al. Mice deficient for the HNK-1 sulfotransferase show alterations in synaptic efficacy and spatial learning and memory. Mol Cell Neurosci 20, 712–729, doi:10.1006/mcne.2002.1142 (2002).

225 Yamamoto, S. et al. Mice deficient in nervous system-specific carbohydrate epitope HNK-1 exhibit impaired synaptic plasticity and spatial learning. J Biol Chem 277, 27227–27231, doi:10.1074/jbc.C200296200 (2002).

226 Yagi, H. et al. HNK-1 epitope-carrying tenascin-C spliced variant regulates the proliferation of mouse embryonic neural stem cells. J Biol Chem 285, 37293–37301, doi:10.1074/jbc.M110.157081 (2010).

227 Weiss, L. A. & Nieto, M. The crux of Cux genes in neuronal function and plasticity. Brain Res 1705, 32–42, doi:10.1016/j.brainres.2018.02.044 (2019).

228 Basu, S. et al. The Mef2c Gene Dose-Dependently Controls Hippocampal Neurogenesis and the Expression of Autism-Like Behaviors. J Neurosci 44, doi:10.1523/JNEUROSCI.1058-23.2023 (2024).

229 Chiu, Y. C. et al. Foxp2 regulates neuronal differentiation and neuronal subtype specification. Dev Neurobiol 74, 723–738, doi:10.1002/dneu.22166 (2014).

230 Manning, C. E. et al. Hippocampal Subgranular Zone FosB Expression Is Critical for Neurogenesis and Learning. Neuroscience 406, 225–233, doi:10.1016/j.neuroscience.2019.03.022 (2019).

231 Wang, H., Matsushita, M. T., Abel, G. M., Storm, D. R. & Xia, Z. Inducible and conditional activation of ERK5 MAP kinase rescues mice from cadmium-induced olfactory memory deficits. Neurotoxicology 81, 127–136, doi:10.1016/j.neuro.2020.09.038 (2020).

232 Ogbonnaya, E. S. et al. Adult Hippocampal Neurogenesis Is Regulated by the Microbiome. Biol Psychiatry 78, e7–9, doi:10.1016/j.biopsych.2014.12.023 (2015).

233 Ribeiro, M. F. et al. Diet-dependent gut microbiota impacts on adult neurogenesis through mitochondrial stress modulation. Brain Commun 2, fcaa165, doi:10.1093/braincomms/fcaa165 (2020).

234 Wei, G. Z. et al. Tryptophan-metabolizing gut microbes regulate adult neurogenesis via the aryl hydrocarbon receptor. Proc Natl Acad Sci U S A 118, doi:10.1073/pnas.2021091118 (2021).

235 Ali, T. et al. Cadmium, an Environmental Contaminant, Exacerbates Alzheimer’s Pathology in the Aged Mice’s Brain. Front Aging Neurosci 13, 650930, doi:10.3389/fnagi.2021.650930 (2021).

236 Hao, R. et al. Caffeic acid phenethyl ester reversed cadmium-induced cell death in hippocampus and cortex and subsequent cognitive disorders in mice: Involvements of AMPK/SIRT1 pathway and amyloid-tau-neuroinflammation axis. Food Chem Toxicol 144, 111636, doi:10.1016/j.fct.2020.111636 (2020).

237 Khan, A., Ikram, M., Muhammad, T., Park, J. & Kim, M. O. Caffeine Modulates Cadmium-Induced Oxidative Stress, Neuroinflammation, and Cognitive Impairments by Regulating Nrf-2/HO-1 In Vivo and In Vitro. J Clin Med 8, doi:10.3390/jcm8050680 (2019).

238 Su, D. et al. Sphk1 mediates neuroinflammation and neuronal injury via TRAF2/NF-kappaB pathways in activated microglia in cerebral ischemia reperfusion. J Neuroimmunol 305, 35–41, doi:10.1016/j.jneuroim.2017.01.015 (2017).

239 Zheng, S. et al. Sphingosine kinase 1 mediates neuroinflammation following cerebral ischemia. Exp Neurol 272, 160–169, doi:10.1016/j.expneurol.2015.03.012 (2015).

240 Hide, I. et al. P2Y(2) receptor mediates dying cell removal via inflammatory activated microglia. J Pharmacol Sci 153, 55–67, doi:10.1016/j.jphs.2023.06.004 (2023).

241 Xi, C. X., Xiong, F., Zhou, Z., Mei, L. & Xiong, W. C. PYK2 interacts with MyD88 and regulates MyD88-mediated NF-kappaB activation in macrophages. J Leukoc Biol 87, 415–423, doi:10.1189/jlb.0309125 (2010).

242 Kunkle, B. W. et al. Genetic meta-analysis of diagnosed Alzheimer’s disease identifies new risk loci and implicates Abeta, tau, immunity and lipid processing. Nat Genet 51, 414–430, doi:10.1038/s41588-019-0358-2 (2019).

243 Lee, J. W. et al. Enhanced phagocytosis associated with multinucleated microglia via Pyk2 inhibition in an acute beta-amyloid infusion model. J Neuroinflammation 21, 196, doi:10.1186/s12974-024-03192-7 (2024).

244 Salazar, S. V. et al. Alzheimer’s Disease Risk Factor Pyk2 Mediates Amyloid-beta-Induced Synaptic Dysfunction and Loss. J Neurosci 39, 758–772, doi:10.1523/JNEUROSCI.1873-18.2018 (2019).

245 Guo, Y., Sun, C. K., Tang, L. & Tan, M. S. Microglia PTK2B/Pyk2 in the Pathogenesis of Alzheimer’s Disease. Curr Alzheimer Res 20, 692–704, doi:10.2174/0115672050299004240129051655 (2023).

246 Brombacher, T. M. et al. IL-4R alpha deficiency influences hippocampal-BDNF signaling pathway to impair reference memory. Sci Rep 10, 16506, doi:10.1038/s41598-020-73574-3 (2020).

247 Brombacher, T. M. et al. IL-4Ralpha deletion disrupts psychomotor performance and reference memory in mice while sparing behavioural phenotype associated with spatial learning. Brain Behav Immun 92, 157–164, doi:10.1016/j.bbi.2020.12.003 (2021).

248 Liu, X., Sun, S. & Liu, D. IL-17D: A Less Studied Cytokine of IL-17 Family. Int Arch Allergy Immunol 181, 618–623, doi:10.1159/000508255 (2020).

249 Lu, Y., Zhang, P., Xu, F., Zheng, Y. & Zhao, H. Advances in the study of IL-17 in neurological diseases and mental disorders. Front Neurol 14, 1284304, doi:10.3389/fneur.2023.1284304 (2023).

250 Ribeiro, M. et al. Meningeal gammadelta T cell-derived IL-17 controls synaptic plasticity and short-term memory. Sci Immunol 4, doi:10.1126/sciimmunol.aay5199 (2019).

251 Li, J. et al. Mechanisms of regulation of glycolipid metabolism by natural compounds in plants: effects on short-chain fatty acids. Nutr Metab (Lond) 21, 49, doi:10.1186/s12986-024-00829-5 (2024).

252 Martin-Gallausiaux, C., Marinelli, L., Blottiere, H. M., Larraufie, P. & Lapaque, N. SCFA: mechanisms and functional importance in the gut. Proc Nutr Soc 80, 37–49, doi:10.1017/S0029665120006916 (2021).

253 Cui, J. et al. Butyrate-Producing Bacteria and Insulin Homeostasis: The Microbiome and Insulin Longitudinal Evaluation Study (MILES). Diabetes 71, 2438–2446, doi:10.2337/db22-0168 (2022).

254 Vital, M., Karch, A. & Pieper, D. H. Colonic Butyrate-Producing Communities in Humans: an Overview Using Omics Data. mSystems 2, doi:10.1128/mSystems.00130-17 (2017).

255 Holland, S. I., Ertan, H., Montgomery, K., Manefield, M. J. & Lee, M. Novel dichloromethane-fermenting bacteria in the Peptococcaceae family. ISME J 15, 1709–1721, doi:10.1038/s41396-020-00881-y (2021).

256 Kong, Y., Jiang, B. & Luo, X. Gut microbiota influences Alzheimer’s disease pathogenesis by regulating acetate in Drosophila model. Future Microbiol 13, 1117–1128, doi:10.2217/fmb-2018-0185 (2018).

257 Wu, L. et al. Altered Gut Microbial Metabolites in Amnestic Mild Cognitive Impairment and Alzheimer’s Disease: Signals in Host-Microbe Interplay. Nutrients 13, doi:10.3390/nu13010228 (2021).

258 Unger, M. M. et al. Short chain fatty acids and gut microbiota differ between patients with Parkinson’s disease and age-matched controls. Parkinsonism Relat Disord 32, 66–72, doi:10.1016/j.parkreldis.2016.08.019 (2016).

259 Peng, L., Li, Z. R., Green, R. S., Holzman, I. R. & Lin, J. Butyrate enhances the intestinal barrier by facilitating tight junction assembly via activation of AMP-activated protein kinase in Caco-2 cell monolayers. J Nutr 139, 1619–1625, doi:10.3945/jn.109.104638 (2009).

260 Kelly, C. J. et al. Crosstalk between Microbiota-Derived Short-Chain Fatty Acids and Intestinal Epithelial HIF Augments Tissue Barrier Function. Cell Host Microbe 17, 662–671, doi:10.1016/j.chom.2015.03.005 (2015).

261 Zheng, L. et al. Microbial-Derived Butyrate Promotes Epithelial Barrier Function through IL-10 Receptor-Dependent Repression of Claudin-2. J Immunol 199, 2976–2984, doi:10.4049/jimmunol.1700105 (2017).

262 Kim, C. H. Control of lymphocyte functions by gut microbiota-derived short-chain fatty acids. Cell Mol Immunol 18, 1161–1171, doi:10.1038/s41423-020-00625-0 (2021).

263 Barcelo, A. et al. Mucin secretion is modulated by luminal factors in the isolated vascularly perfused rat colon. Gut 46, 218–224, doi:10.1136/gut.46.2.218 (2000).

264 Schwiertz, A. et al. Fecal markers of intestinal inflammation and intestinal permeability are elevated in Parkinson’s disease. Parkinsonism Relat Disord 50, 104–107, doi:10.1016/j.parkreldis.2018.02.022 (2018).

265 Buscarinu, M. C. et al. Intestinal Permeability in Relapsing-Remitting Multiple Sclerosis. Neurotherapeutics 15, 68–74, doi:10.1007/s13311-017-0582-3 (2018).

266 Ou, Z. et al. Protective effects of Akkermansia muciniphila on cognitive deficits and amyloid pathology in a mouse model of Alzheimer’s disease. Nutr Diabetes 10, 12, doi:10.1038/s41387-020-0115-8 (2020).

267 Yang, X., Yu, D., Xue, L., Li, H. & Du, J. Probiotics modulate the microbiota-gut-brain axis and improve memory deficits in aged SAMP8 mice. Acta Pharm Sin B 10, 475–487, doi:10.1016/j.apsb.2019.07.001 (2020).

268 Fang, X. et al. Evaluation of the Anti-Aging Effects of a Probiotic Combination Isolated From Centenarians in a SAMP8 Mouse Model. Front Immunol 12, 792746, doi:10.3389/fimmu.2021.792746 (2021).

269 Samuel, B. S. et al. Effects of the gut microbiota on host adiposity are modulated by the short-chain fatty-acid binding G protein-coupled receptor, Gpr41. Proc Natl Acad Sci U S A 105, 16767–16772, doi:10.1073/pnas.0808567105 (2008).

270 Holzer, P., Reichmann, F. & Farzi, A. Neuropeptide Y, peptide YY and pancreatic polypeptide in the gut-brain axis. Neuropeptides 46, 261–274, doi:10.1016/j.npep.2012.08.005 (2012).

271 Holst, J. J. The physiology of glucagon-like peptide 1. Physiol Rev 87, 1409–1439, doi:10.1152/physrev.00034.2006 (2007).

272 Holzer, P. Neuropeptides, Microbiota, and Behavior. Int Rev Neurobiol 131, 67–89, doi:10.1016/bs.irn.2016.08.005 (2016).

273 Kastin, A. J. & Akerstrom, V. Nonsaturable entry of neuropeptide Y into brain. Am J Physiol 276, E479–482, doi:10.1152/ajpendo.1999.276.3.E479 (1999).

274 Lin, S., Boey, D. & Herzog, H. NPY and Y receptors: lessons from transgenic and knockout models. Neuropeptides 38, 189–200, doi:10.1016/j.npep.2004.05.005 (2004).

275 dos Santos, V. V., et al. Neuropeptide Y (NPY) prevents depressive-like behavior, spatial memory deficits and oxidative stress following amyloid-beta (Abeta(1-40)) administration in mice. Behav Brain Res 244, 107–115, doi:10.1016/j.bbr.2013.01.039 (2013).

276 Beck, B. & Pourie, G. Ghrelin, neuropeptide Y, and other feeding-regulatory peptides active in the hippocampus: role in learning and memory. Nutr Rev 71, 541–561, doi:10.1111/nure.12045 (2013).

277 Pain, S., Brot, S. & Gaillard, A. Neuroprotective Effects of Neuropeptide Y against Neurodegenerative Disease. Curr Neuropharmacol 20, 1717–1725, doi:10.2174/1570159X19666210906120302 (2022).

278 Regev, L. & Baram, T. Z. Corticotropin releasing factor in neuroplasticity. Front Neuroendocrinol 35, 171–179, doi:10.1016/j.yfrne.2013.10.001 (2014).

279 Blank, T., Nijholt, I., Eckart, K. & Spiess, J. Priming of long-term potentiation in mouse hippocampus by corticotropin-releasing factor and acute stress: implications for hippocampus-dependent learning. J Neurosci 22, 3788–3794, doi:10.1523/JNEUROSCI.22-09-03788.2002 (2002).

280 White, C. M., Ji, S., Cai, H., Maudsley, S. & Martin, B. Therapeutic potential of vasoactive intestinal peptide and its receptors in neurological disorders. CNS Neurol Disord Drug Targets 9, 661–666, doi:10.2174/187152710793361595 (2010).

281 Chapter, M. C. et al. Chemical modification of class II G protein-coupled receptor ligands: frontiers in the development of peptide analogs as neuroendocrine pharmacological therapies. Pharmacol Ther 125, 39–54, doi:10.1016/j.pharmthera.2009.07.006 (2010).

282 Correa-Oliveira, R., Fachi, J. L., Vieira, A., Sato, F. T. & Vinolo, M. A. Regulation of immune cell function by short-chain fatty acids. Clin Transl Immunology 5, e73, doi:10.1038/cti.2016.17 (2016).

283 Chang, P. V., Hao, L., Offermanns, S. & Medzhitov, R. The microbial metabolite butyrate regulates intestinal macrophage function via histone deacetylase inhibition. Proc Natl Acad Sci U S A 111, 2247–2252, doi:10.1073/pnas.1322269111 (2014).

284 Ang, Z. et al. Human and mouse monocytes display distinct signalling and cytokine profiles upon stimulation with FFAR2/FFAR3 short-chain fatty acid receptor agonists. Sci Rep 6, 34145, doi:10.1038/srep34145 (2016).

285 Huuskonen, J., Suuronen, T., Nuutinen, T., Kyrylenko, S. & Salminen, A. Regulation of microglial inflammatory response by sodium butyrate and short-chain fatty acids. Br J Pharmacol 141, 874–880, doi:10.1038/sj.bjp.0705682 (2004).

286 Ransohoff, R. M. How neuroinflammation contributes to neurodegeneration. Science 353, 777–783, doi:10.1126/science.aag2590 (2016).

287 Saib, S. & Delavenne, X. Inflammation Induces Changes in the Functional Expression of P-gp, BCRP, and MRP2: An Overview of Different Models and Consequences for Drug Disposition. Pharmaceutics 13, doi:10.3390/pharmaceutics13101544 (2021).

288 Zangerolamo, L., Vettorazzi, J. F., Rosa, L. R. O., Carneiro, E. M. & Barbosa, H. C. L. The bile acid TUDCA and neurodegenerative disorders: An overview. Life Sci 272, 119252, doi:10.1016/j.lfs.2021.119252 (2021).

289 Huang, F., Pariante, C. M. & Borsini, A. From dried bear bile to molecular investigation: A systematic review of the effect of bile acids on cell apoptosis, oxidative stress and inflammation in the brain, across pre-clinical models of neurological, neurodegenerative and neuropsychiatric disorders. Brain Behav Immun 99, 132–146, doi:10.1016/j.bbi.2021.09.021 (2022).

290 Mano, N. et al. Presence of protein-bound unconjugated bile acids in the cytoplasmic fraction of rat brain. J Lipid Res 45, 295–300, doi:10.1194/jlr.M300369-JLR200 (2004).

291 Kippler, M. et al. Early-life cadmium exposure and child development in 5-year-old girls and boys: a cohort study in rural Bangladesh. Environ Health Perspect 120, 1462–1468, doi:10.1289/ehp.1104431 (2012).

292 Gade, M., Comfort, N. & Re, D. B. Sex-specific neurotoxic effects of heavy metal pollutants: Epidemiological, experimental evidence and candidate mechanisms. Environ Res 201, 111558, doi:10.1016/j.envres.2021.111558 (2021).

293 Lamtai, M., Chaibat, J., Ouakki, S., Berkiks, I., Rifi, El-H., El Hessni, A., Mesfioui, A., Tadlaoui Hbibi, A., Ahyayauch, H., Essamri, A. and Ouichou, A. Effect of Chronic Administration of Cadmium on Anxiety-Like, Depression-Like and Memory Deficits in Male and Female Rats: Possible Involvement of Oxidative Stress Mechanism Journal of Behavioral and Brain Science 8, 28 (2018).

294 Zhang, A. et al. Cadmium exposure modulates the gut-liver axis in an Alzheimer’s disease mouse model. Commun Biol 4, 1398, doi:10.1038/s42003-021-02898-1 (2021).

