## Supplementary material for "Cadmium-induced gut dysbiosis precedes the onset of hippocampus-dependent learning and memory deficits in mice": manuscript_supplementary data_v11.pdf

**Figure.S1**

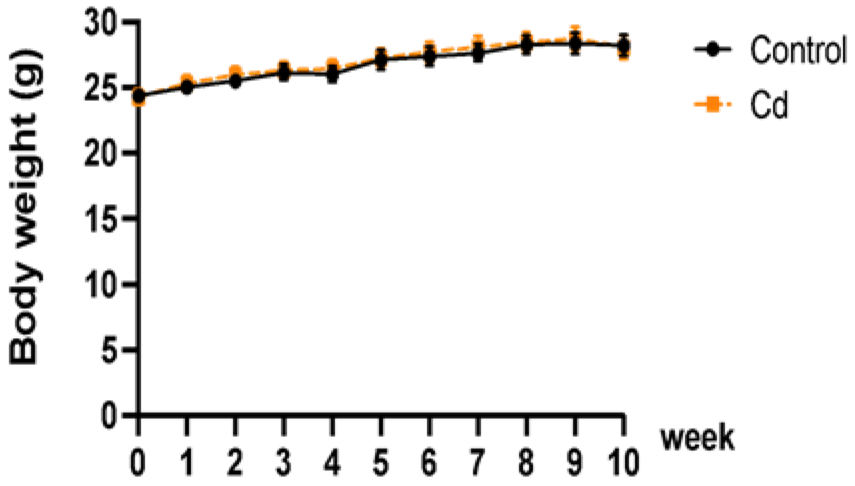

Figure.S2

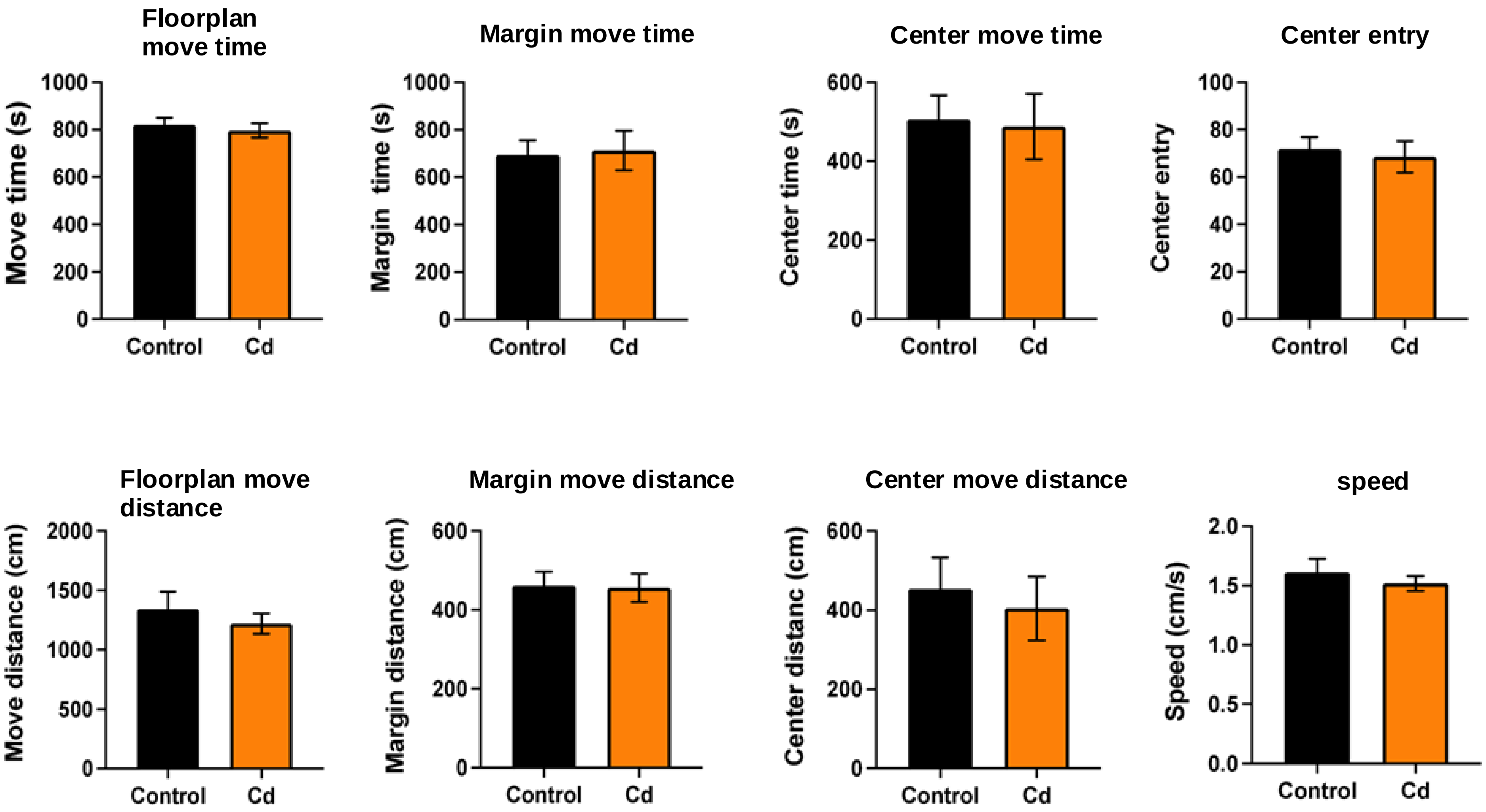

Floorplan move time
Move time (s)
1000
800
600
400
200
0
Control
Cd
Margin move time
Margin time (s)
1000
800
600
400
200
0
Control
Cd
Center move time
Center time (s)
600
400
200
0
Control
Cd
Center entry
Center entry
100
80
60
40
20
0
Control
Cd
Floorplan move distance
Move distance (cm)
2000
1500
1000
500
0
Control
Cd
Margin move distance
Margin distance (cm)
600
400
200
0
Control
Cd
Center move distance
Center distanc (cm)
600
400
200
0
Control
Cd
speed
Speed (cm/s)
2.0
1.5
1.0
0.5
0.0
Control
Cd

Figure.S3

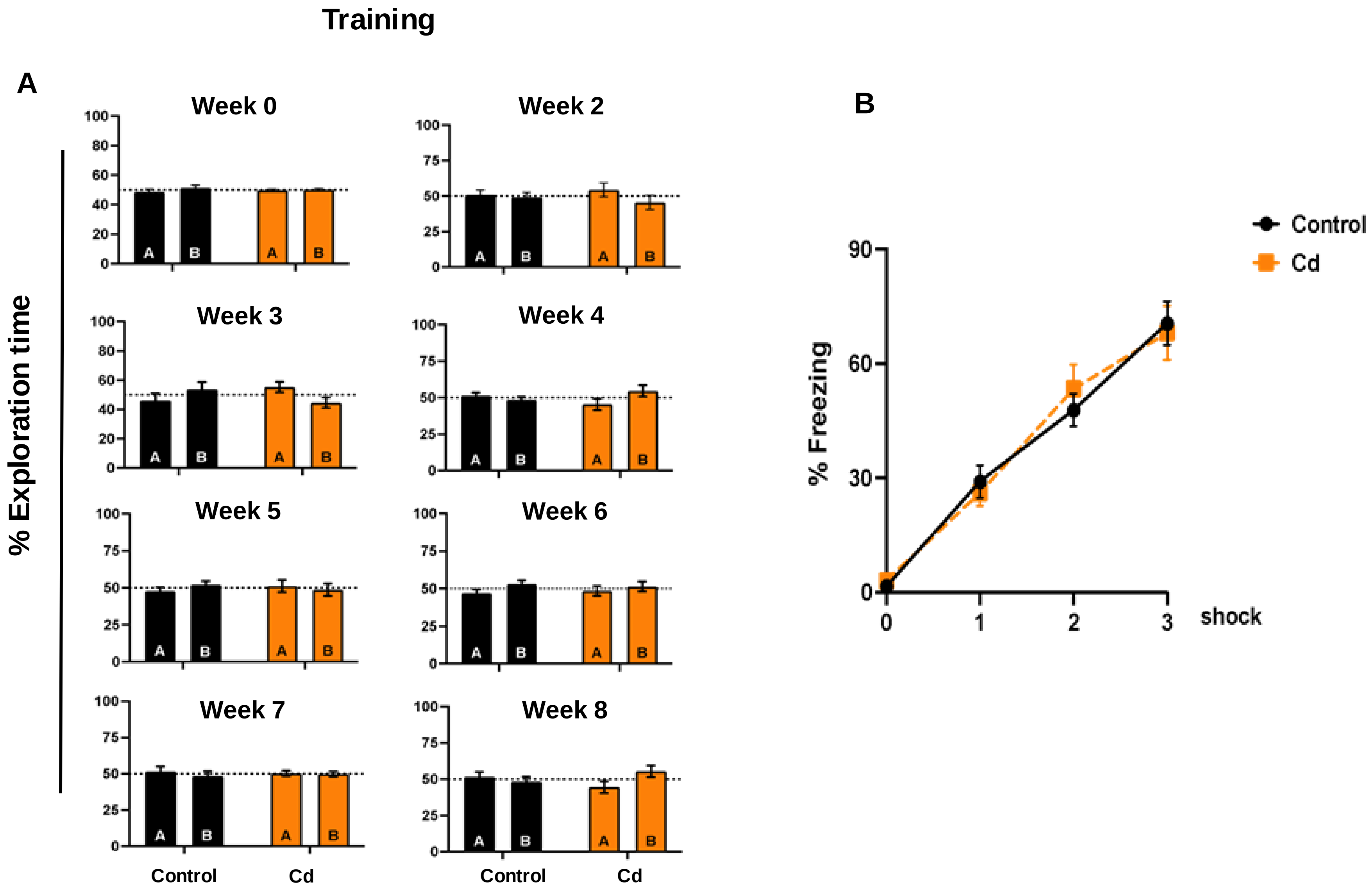

Training
A
Week 0
Week 2
Week 3
Week 4
Week 5
Week 6
Week 7
Week 8
% Exploration time
A
B
Control
Cd
B
% Freezing
shock
Control
Cd

**Figure S4**

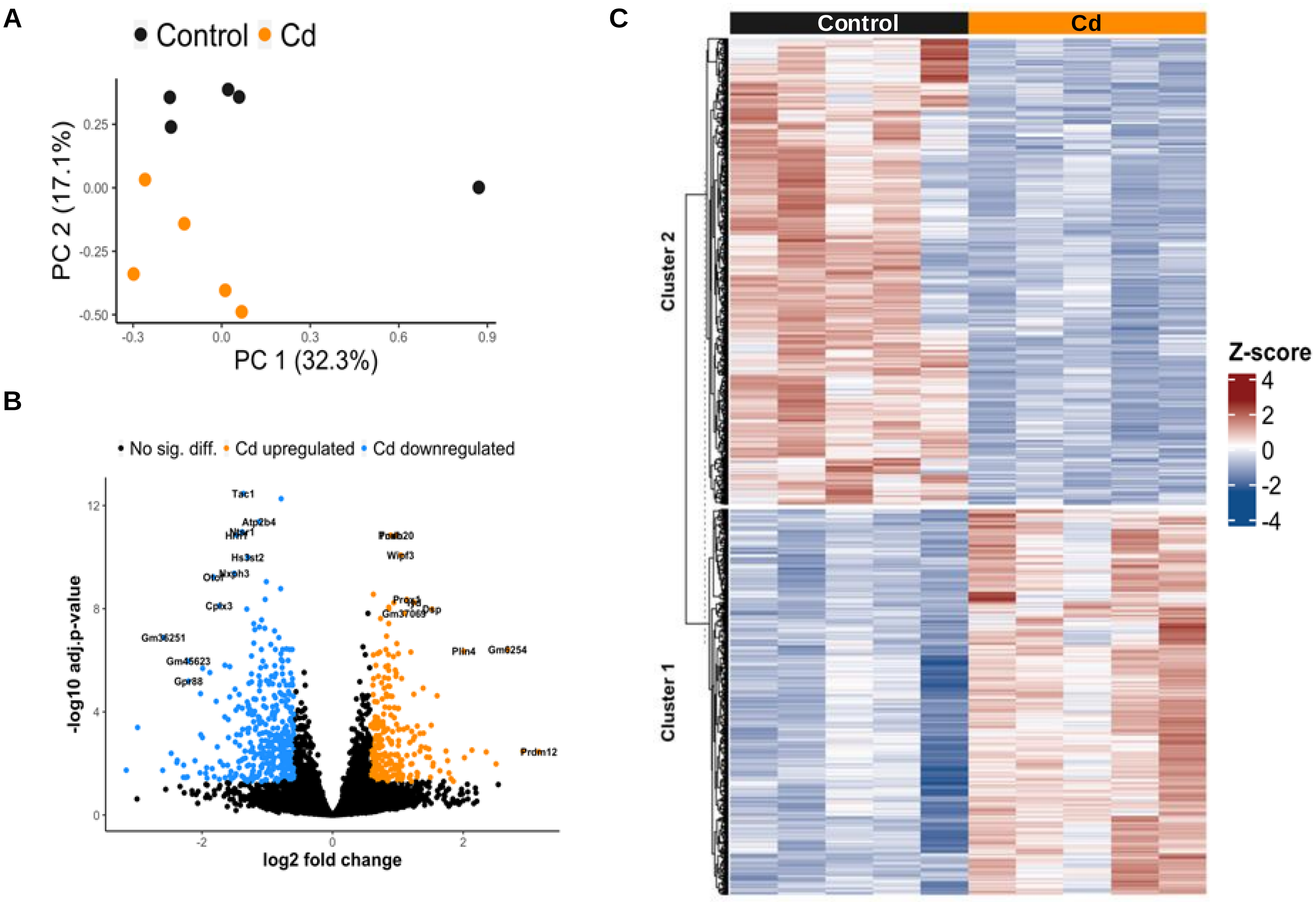
A
Control
Cd
PC 2 (17.1%)
PC 1 (32.3%)
0.25
0.00
-0.25
-0.50
-0.3
0.0
0.3
0.6
0.9
B
No sig. diff.
Cd upregulated
Cd downregulated
-log10 adj.p-value
log2 fold change
Tac1
Atp2b4
Hs3st2
Cplx3
Gm36251
Gr88
Wipf3
Plin4
Gm6254
Prdm12
Dsp
12
8
4
0
-2
0
2
C
Control
Cd
Cluster 2
Cluster 1
Z-score
4
2
0
-2
-4

Figure.S5

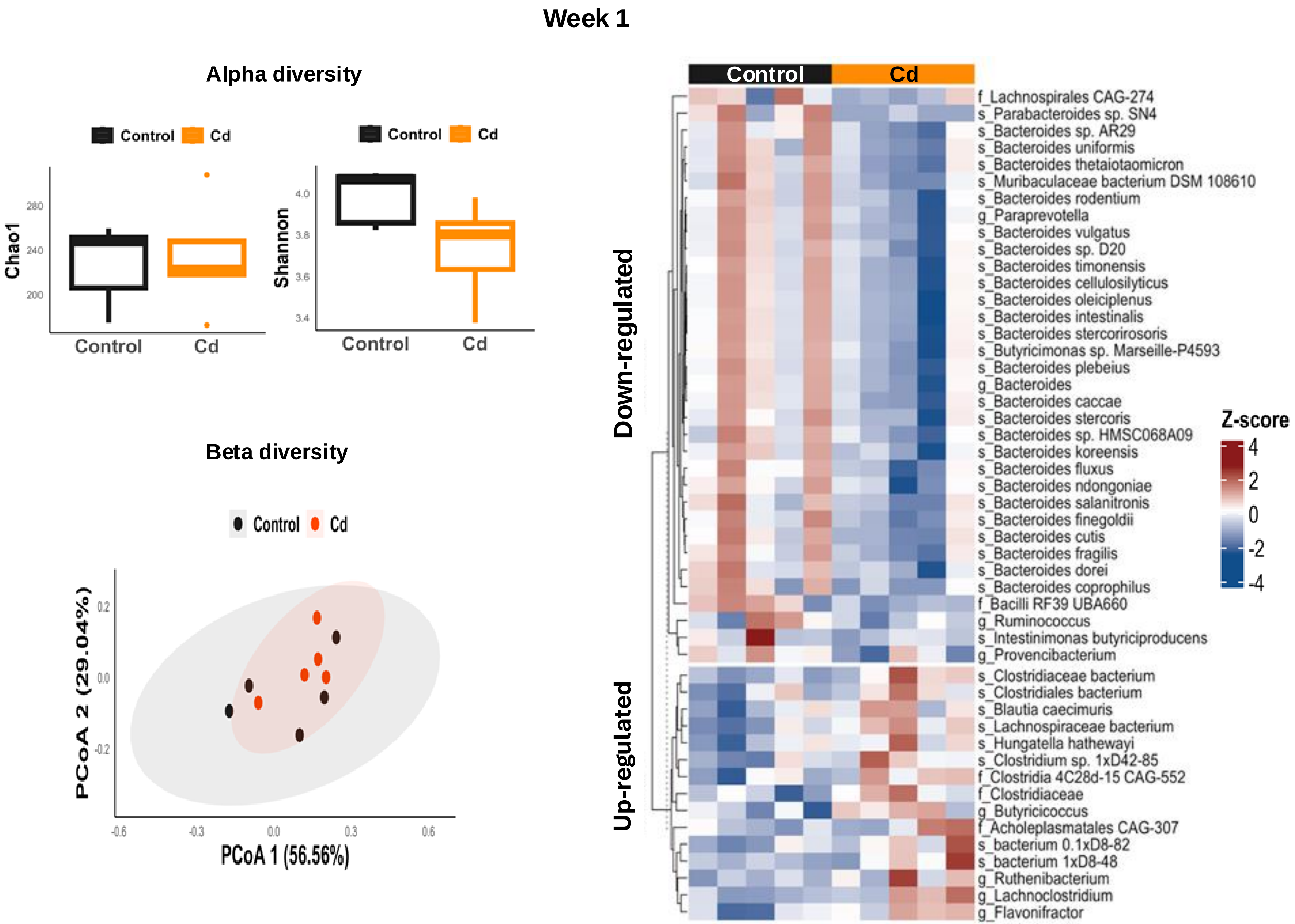

Week 1
Alpha diversity
Control Cd
Chao1
280
240
200
Control
Cd
Control Cd
Shannon
4.0
3.8
3.6
3.4
Control
Cd
Beta diversity
Control Cd
PCoA 2 (29.04%)
0.2
0.0
-0.2
-0.6
-0.3
0.0
0.3
0.6
PCoA 1 (56.56%)
Control
Cd
Down-regulated
Up-regulated
f_Lachnospirales CAG-274
s_Parabacteroides sp. SN4
s_Bacteroides sp. AR29
s_Bacteroides uniformis
s_Bacteroides thetaiotaomicron
s_Muribaculaceae bacterium DSM 108610
s_Bacteroides rodentium
g_Paraprevotella
s_Bacteroides vulgatus
s_Bacteroides sp. D20
s_Bacteroides timonensis
s_Bacteroides cellulosilyticus
s_Bacteroides oleiciplenus
s_Bacteroides intestinalis
s_Bacteroides stercorirosoris
s_Butyricimonas sp. Marseille-P4593
s_Bacteroides plebeius
g_Bacteroides
s_Bacteroides caccae
s_Bacteroides stercoris
s_Bacteroides sp. HMSC068A09
s_Bacteroides koreensis
s_Bacteroides fluxus
s_Bacteroides ndongoniae
s_Bacteroides salanitronis
s_Bacteroides finegoldii
s_Bacteroides cutis
s_Bacteroides fragilis
s_Bacteroides dorei
s_Bacteroides coprophilus
f_Bacilli RF39 UBA660
g_Ruminococcus
s_Intestinimonas butyriciproducens
g_Provencibacterium
s_Clostridiaceae bacterium
s_Clostridiales bacterium
s_Blautia caecimuris
s_Lachnospiraceae bacterium
s_Hungatella hathewayi
s_Clostridium sp. 1xD42-85
f_Clostridia 4C28d-15 CAG-552
f_Clostridiaceae
g_Butyricicoccus
f_Acholeplasmatales CAG-307
s_bacterium 0.1xD8-82
s_bacterium 1xD8-48
g_Ruthenibacterium
g_Lachnoclostridium
g_Flavonifractor
Z-score
4
2
0
-2
-4

**Figure.S6**

#### Week 3

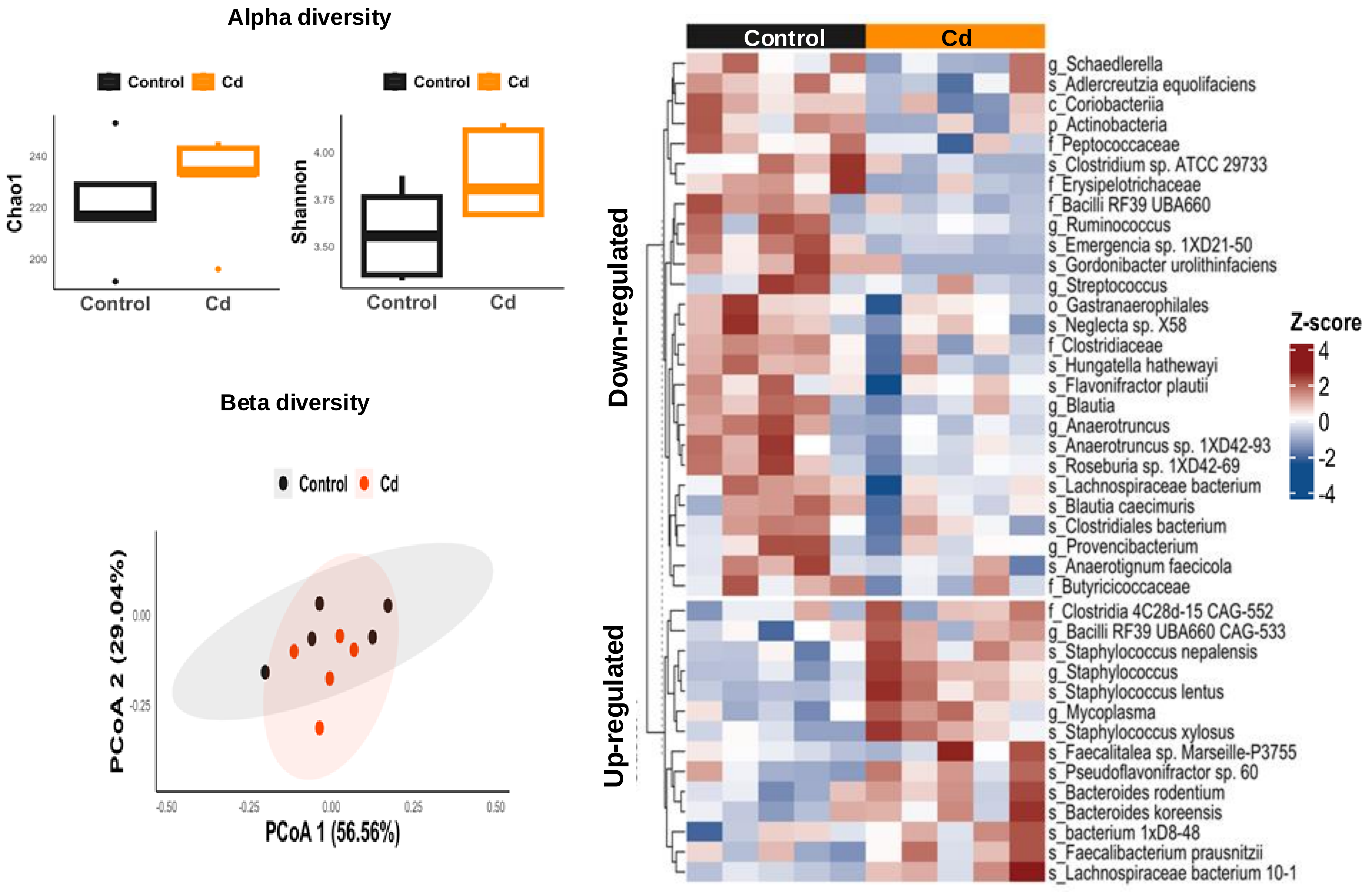

Figure.S7

### Week 5

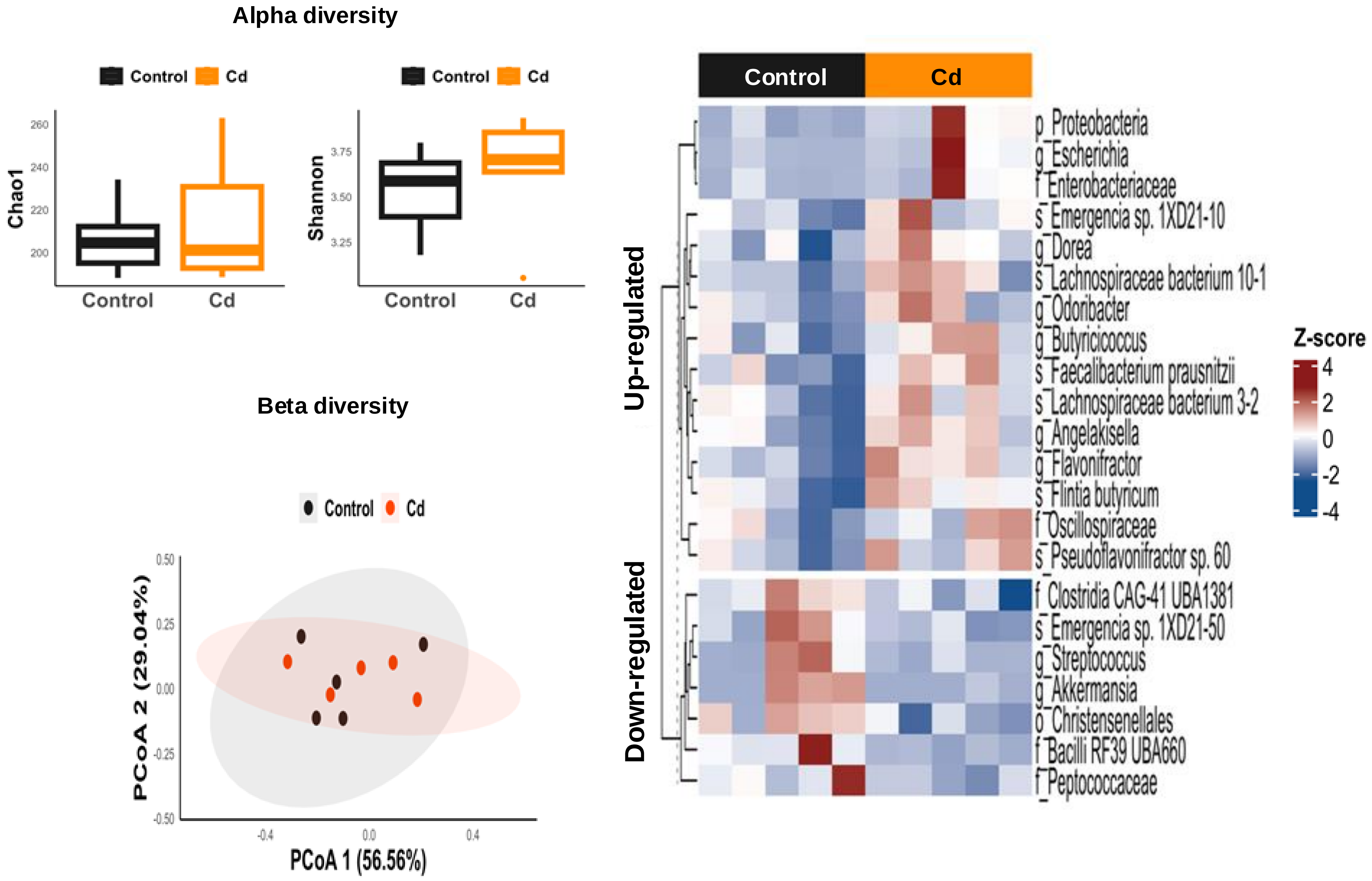

**Figure.S8**

#### Week 7

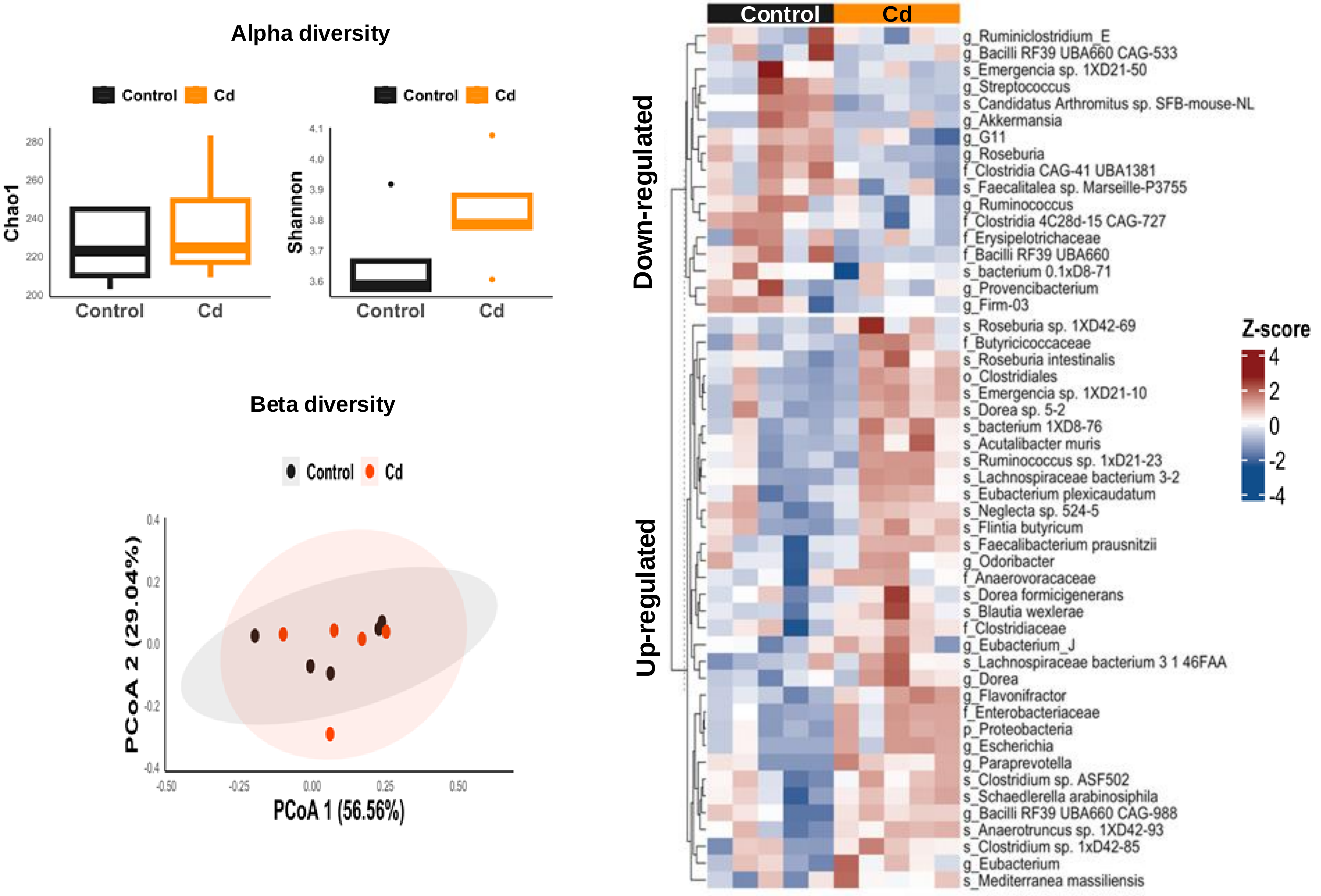

Figure S9

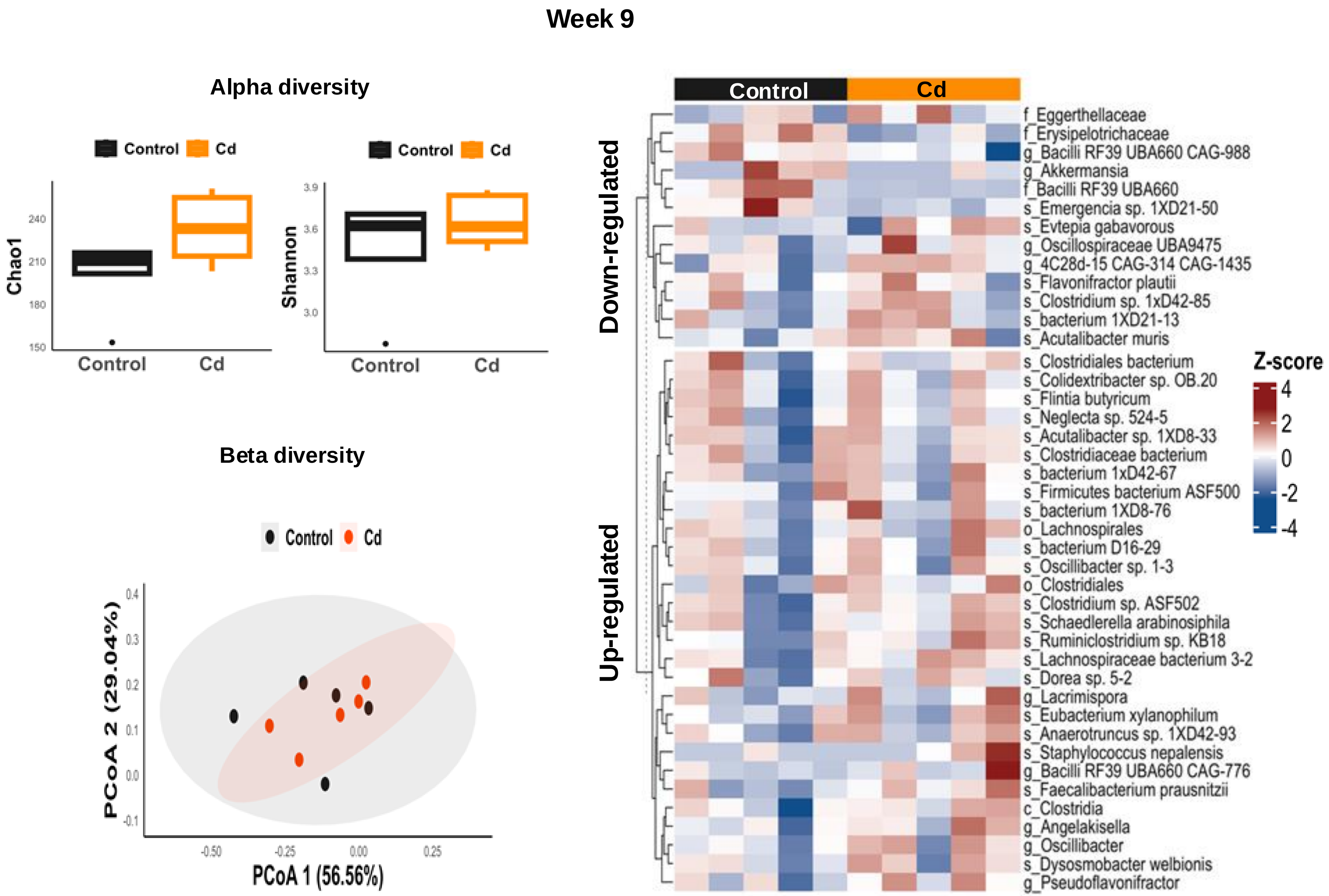

Week 9
Alpha diversity
Control
Cd
Chao1
150
180
210
240
Control
Cd
Shannon
3.0
3.3
3.6
3.9
Beta diversity
Control
Cd
PCoA 2 (29.04%)
PCoA 1 (56.56%)
-0.50
-0.25
0.00
0.25
-0.1
0.0
0.1
0.2
0.3
0.4
Control
Cd
Down-regulated
Up-regulated
Z-score
4
2
0
-2
-4
f_Eggerthellaceae
f_Erysipelotrichaceae
g_Bacilli RF39 UBA660 CAG-988
g_Akkermansia
f_Bacilli RF39 UBA660
s_Emergencia sp. 1XD21-50
s_Evtepia gabavorous
g_Oscillospiraceae UBA9475
g_4C28d-15 CAG-314 CAG-1435
s_Flavonifractor plautii
s_Clostridium sp. 1xD42-85
s_bacterium 1XD21-13
s_Acutalibacter muris
s_Clostridiales bacterium
s_Colidextribacter sp. OB.20
s_Flintia butyricum
s_Neglecta sp. 524-5
s_Acutalibacter sp. 1XD8-33
s_Clostridiaceae bacterium
s_bacterium 1xD42-67
s_Firmicutes bacterium ASF500
s_bacterium 1XD8-76
o_Lachnospirales
s_bacterium D16-29
s_Oscillibacter sp. 1-3
o_Clostridiales
s_Clostridium sp. ASF502
s_Schaedlerella arabinosiphila
s_Ruminiclostridium sp. KB18
s_Lachnospiraceae bacterium 3-2
s_Dorea sp. 5-2
g_Lacrimispora
s_Eubacterium xylanophilum
s_Anaerotruncus sp. 1XD42-93
s_Staphylococcus nepalensis
g_Bacilli RF39 UBA660 CAG-776
s_Faecalibacterium prausnitzii
c_Clostridia
g_Angelakisella
g_Oscillibacter
s_Dysosmobacter welbionis
g_Pseudoflavonifractor

**Figure S10**

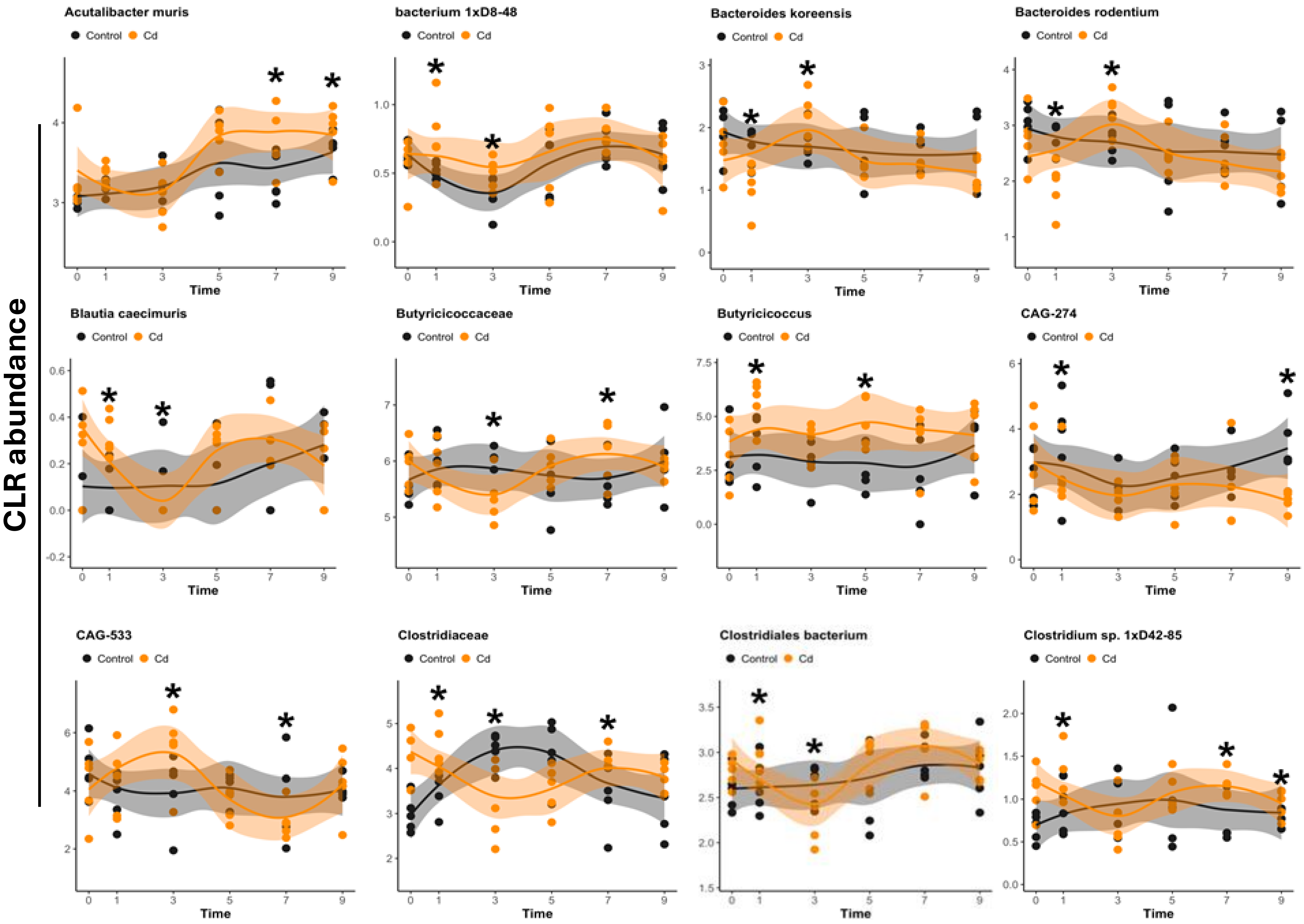
CLR abundance
Acutalibacter muris
bacterium 1xD8-48
Bacteroides koreensis
Bacteroides rodentium
Blautia caecimuris
Butyricicoccaceae
Butyricicoccus
CAG-274
CAG-533
Clostridiaceae
Clostridiales bacterium
Clostridium sp. 1xD42-85
Control
Cd
Time

Figure S11

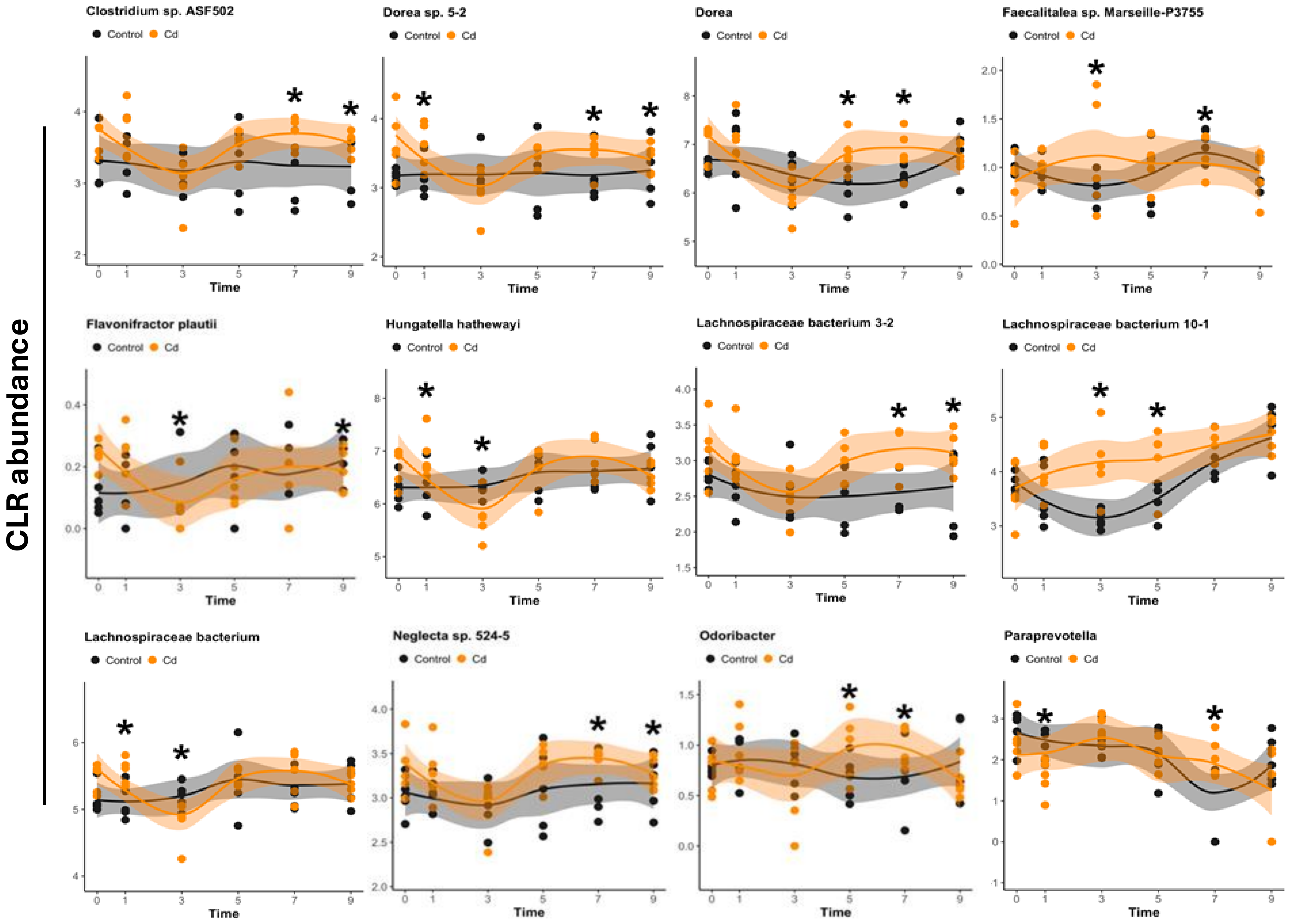
CLR abundance
Clostridium sp. ASF502
Dorea sp. 5-2
Dorea
Faecalitalea sp. Marseille-P3755
Flavonifractor plautii
Hungatella hathewayi
Lachnospiraceae bacterium 3-2
Lachnospiraceae bacterium 10-1
Lachnospiraceae bacterium
Neglecta sp. 524-5
Odoribacter
Paraprevotella
Control
Cd
Time

**Figure S12**

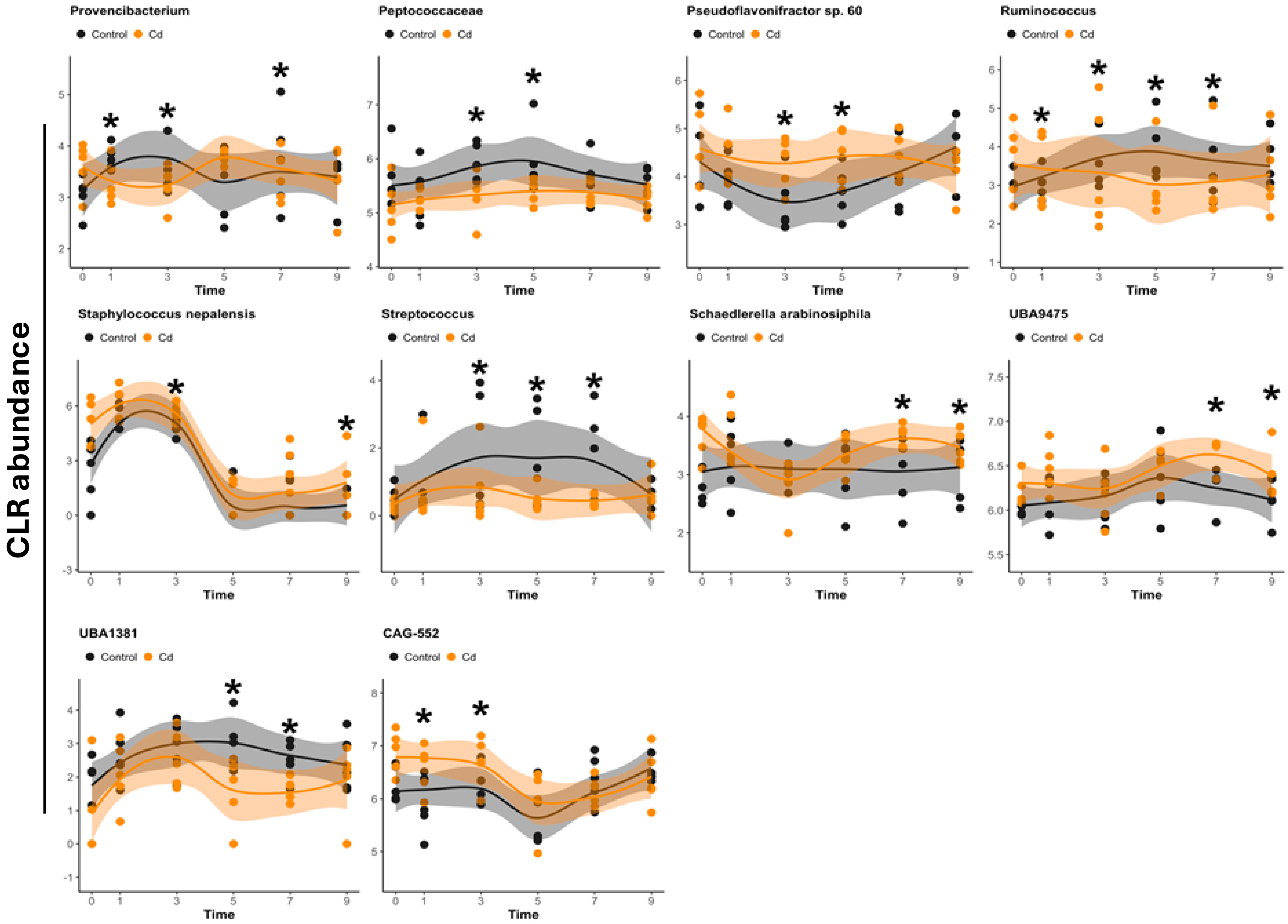

CLR abundance
Provencibacterium
Peptococcaceae
Pseudoflavonifractor sp. 60
Ruminococcus
Staphylococcus nepalensis
Streptococcus
Schaedlerella arabinosiphila
UBA9475
UBA1381
CAG-552
Control
Cd
Time

Figure S13

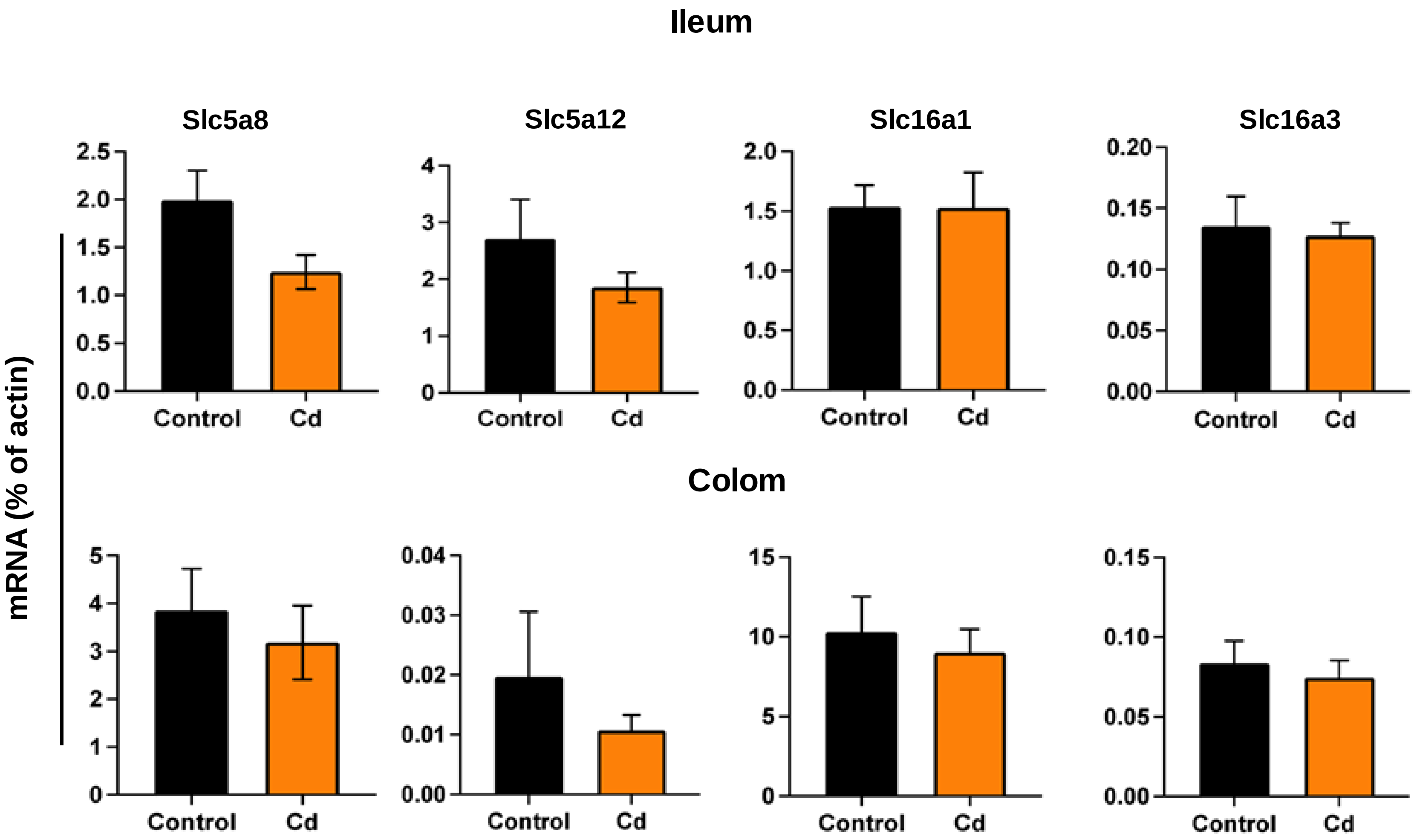

Ileum
Slc5a8
Slc5a12
Slc16a1
Slc16a3
mRNA (% of actin)
2.5
2.0
1.5
1.0
0.5
0.0
4
3
2
1
0
0.20
0.15
0.10
0.05
0.00
Control
Cd
Colom
5
0.04
0.03
0.02
0.01
15
10
0.15

**Figure S14**

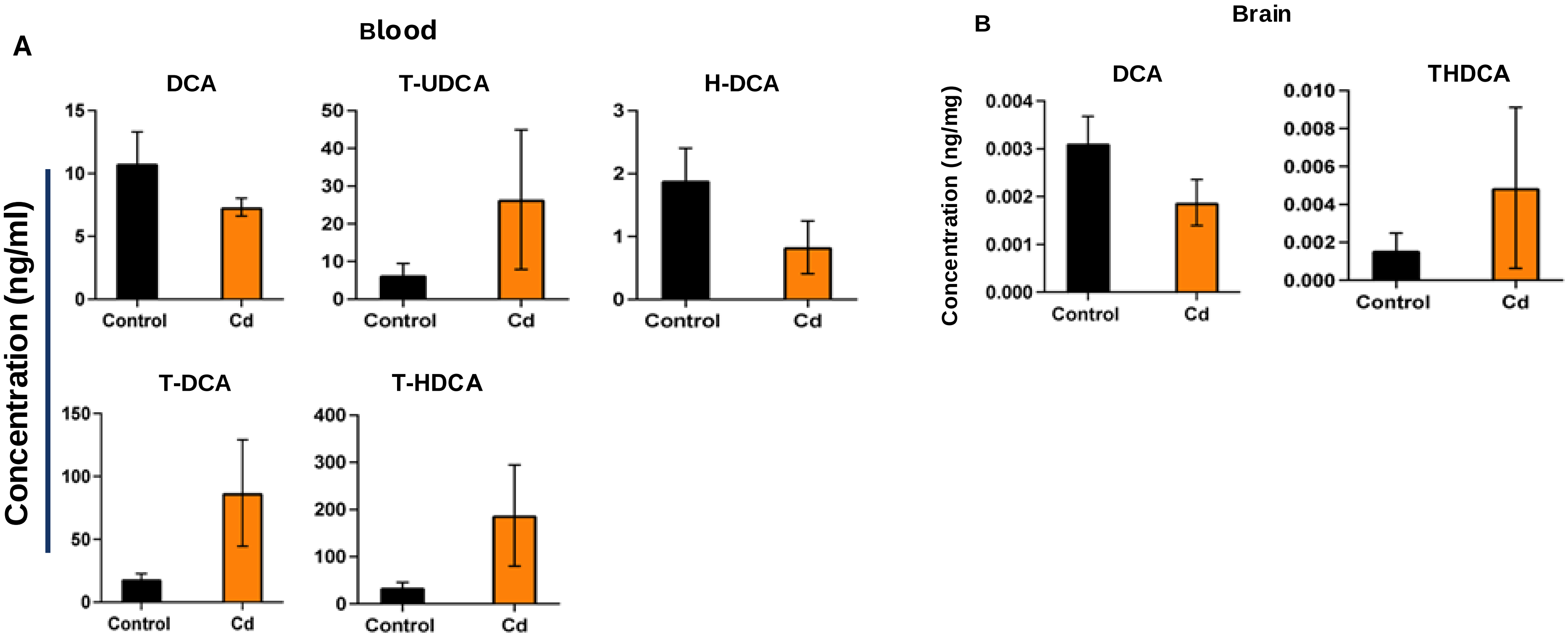
A
Blood
DCA
T-UDCA
H-DCA
T-DCA
T-HDCA
Concentration (ng/ml)
Control
Cd
B
Brain
DCA
THDCA
Concentration (ng/mg)
