## Supplementary material for "Cadmium-induced gut dysbiosis precedes the onset of hippocampus-dependent learning and memory deficits in mice": supplementary data_figure_gut_brain_paper_V11.pdf

**Figure S1. Cd exposure did not affect the body weights of mice.** Data are presented as mean ± SEM. n=8 in each group.

**Figure S2. Cd exposure does not affect the locomotor activity or anxiety of mice in open field test.** For locomotor activity, there was no significant difference the control and Cd-exposed group in Floorplan move time, Floorplan move distance, or speed. For anxiety, there was no significant difference between control and Cd-exposed group in margin move time and distance, center move time and distance, or number of center entry. In addition, both groups had similar center entry times. Data are presented as mean ± SEM. n=8 in each group.

**Figure S3. Animals spent equal amount of time exploring each object/location in the training session of the NOL test (A). Both control and Cd-exposed mice had similar freezing behavior after three foot-shock in the fear conditioning training session (B).** Data are presented as mean ± SEM. n=8 in each group.

**Figure S4. Dysregulated gene expression in the hippocampus of mice after 9 weeks Cd exposure. (A)** Principal component analysis (PCoA) measure separated by Cd treatment. (**B)** Volcano plot of the significant differentially expressed genes (DEGs) in the hippocampus between the control and Cd-exposed mice. (**C)** The heatmap shows the significant DEGs in the hippocampus between the control and Cd-exposed mice. The colors of the heatmap represent the $log_2$-fold change of hippocampus genes of the Cd-exposed mice as compared with the control group. Red indicates upregulated, and Blue indicates downregulated from Cd-exposed mice (Bonferroni-adjusted p-value < 0.05). n = 5 in each group.

**Figure S5. Cd exposure induces dysregulated gut microbiome at one week into exposure.** Cd exposure did not affect Alpha or Beta diversity, but induced alteration of gut microbiome. Red and blue colors show statistically significant increases or decreases in taxa abundance, respectively (adjusted p-value < 0.05). n = 5 in each group.

**Figure S6. Cd exposure induces dysregulated gut microbiome at three weeks into exposure.** Cd exposure did not affect Alpha or Beta diversity, but induced alteration of gut microbiome. Red and blue colors show statistically significant increases or decreases in taxa abundance, respectively (adjusted p-value < 0.05). n = 5 in each group.

**Figure S7. Cd exposure induces dysregulated gut microbiome at five weeks into exposure.** Cd exposure did not affect Alpha or Beta diversity, but induced alteration of gut microbiome. Red and blue colors show statistically significant increases or decreases in taxa abundance, respectively (adjusted p-value < 0.05). n = 5 in each group.

**Figure S8. Cd exposure induces dysregulated gut microbiome at seven weeks into exposure.** Cd exposure did not affect Alpha or Beta diversity, but induced alteration of gut microbiome. Red and blue colors show statistically significant increases or decreases in taxa abundance, respectively (adjusted p-value < 0.05). n = 5 in each group.

**Figure S9. Cd exposure induces dysregulated gut microbiome at nine weeks into exposure.** Cd exposure did not affect Alpha or Beta diversity, but induced alteration of gut microbiome. Red and blue colors show statistically significant increases or decreases in taxa abundance, respectively (adjusted p-value < 0.05). n = 5 in each group.

**Figure S10 – S12. Cd exposure induces persistent alteration of gut microbiomes.** The persistently dysregulated microbiome that are significantly affected by Cd exposure at two or more than two time points. n = 5 in each group. *. adjusted p-value < 0.05.

**Figure S13. Cd exposure did not affect the expression of short-chain fatty acids-related transporters in the large and small intestines.** n = 7-8 in each group

**Figure S14. The levels of secondary bile acids in the blood (A) and brain (B).** n = 6-8 in each group.

**Table S1. Primer sequences used in RT-qPCR**

| Target Genes | Forward Primer Sequence | Reverse Primer Sequence |
|---|---|---|
| *Cldn1* | GGACTGTGGATGTCCTGCGTTT | GCCAATTACCATCAAGGCTCGG |
| *Cldn2* | AGGACTTCCTGCTGACATCCAG | AATCCTGGCAGAACACGGTGCA |
| *Cldn7* | CTGCCTTGGTAGCATGTTCCTG | CCAGCCGATAAAGATGGCAGGT |
| *F11r (Jam1)* | CACCTACTCTGGCTTCTCCTCT | TGCCACTGGATGAGAAGGTGAC |
| *Jam2* | CAGACTGGAGTGGAAGAAGGTG | GCTGACTTCACAGCGATACTCTC |
| *Tjp1* | TTAAGCCTCCGGAAGTAGCA | GGAGCCTGTAGAGCGTTTTG |
| *Tjp2* | AATGGAAAGGTTGGCAACTG | CGTGCTTGTCCTGCTCAATA |
| *Hcar2* | GATCCCCTCCAGTTTCGTGG | TTTGAGTCCCAGATGCACCC |
| *Ffar2* | AATTTCCTGGTGTGCTTTGG | ACCAGACCAACTTCTGGGTG |
| *Ffar3* | CTTCTTTCTTGGCAATTACTGGC | CCGAAATGGTCAGGTTTAGCAA |
| *Abcg2 (Bcrp)* | GAACTCCAGAGCCGTTAGGAC | CAGAATAGCATTAAGGCCAGGTT |
| *Slc5a8* | TTATGGGCGGTCGCAGTATG | CAAAACGGTAGACCTCGGCA |
| *Slc5a12* | CTGCTGAAGTCTACCGCTTTG | CATAGGTGCTTGTGATGCCAG |
| *Slc16a1* | AATGCTGCCCTGTCCTCCTA | CCCAGTACGTGTATTTGTAGTCTCCAT |
| *Slc16a3* | CAGCTTTGCCATGTTCTTCA | AGCCATGAGCACCTCAAACT |
| *Gapdh* | GGCAAATTCAACGGCACAGT | GTCTCGCTCCTGGAAGATGG |
| *Actb* | GGCCAACCGTGAAAAGATGA | CAGCCTGGATGGCTACGTACA |
